# Mitochondria dysregulation contributes to secondary neurodegeneration progression post-contusion injury in human 3D in vitro triculture brain tissue model

**DOI:** 10.1101/2022.09.23.509276

**Authors:** Volha Liaudanskaya, Nicholas J Fiore, Yang Zhang, Yuka Milton, Marilyn F Kelly, Marly Coe, Ariana Barreiro, Victoria K Rose, Matthew R Shapiro, Adam S Mullis, Anna Shevzov-Zebrun, Mathew Blurton-Jones, Michael J Whalen, Aviva J Symes, Irene Georgakoudi, Thomas JF Nieland, David L Kaplan

## Abstract

Traumatic Brain injury-induced disturbances in mitochondrial fission-and-fusion dynamics have been linked to the onset and propagation of neuroinflammation and neurodegeneration. However, cell-type-specific contributions and crosstalk between neurons, microglia, and astrocytes in mitochondria-driven neurodegeneration after brain injury remain undefined. We developed a human three-dimensional *in vitro* triculture tissue model of a contusion injury, composed of neurons, microglia, and astrocytes, and examined the contributions of mitochondrial dysregulation to neuroinflammation and progression of injury-induced neurodegeneration. Pharmacological studies presented here suggest that fragmented mitochondria released by microglia are a key contributor to secondary neuronal damage progression after contusion injury, a pathway that requires astrocyte-microglia crosstalk. Controlling mitochondrial dysfunction thus offers an exciting option for the development of therapies for TBI patients.

## Introduction

Traumatic brain injury (TBI) is a debilitating condition with multifaceted pathobiology that leads to long-lasting disabilities and morbidity. Primary TBI comprises the direct physical injury to the brain and results in acute local neuronal death, excitotoxicity, and edema (1). This is a complete and irreversible event that triggers a series of subsequent effects that can progress for decades. These secondary injury effects are potentially preventable and manifest in chronic glial neuroinflammation, diffused axonal degeneration, and death of the surrounding tissue (2-4). Astrocytes and microglia have dual roles in the progression of brain injuries (5, 6), promoting injurious inflammation and providing homeostatic and regenerative support (3, 6-8). Mitochondria play a crucial role in regulating glial activity and function, through balanced mechanisms of fission and fusion that maintain their growth, shape, distribution, and structure (9). Disruptions in mitochondrial function influence crucial pathways, such as activation of inflammation, energy production, metabolism and calcium homeostasis, phosphorylation of transcription factors, and various death pathways regulation (10, 11). Crosstalk between microglia and astrocytes has been shown to accelerate brain injury progression (12), where microglia push astrocytes toward a neurotoxic state (A1 astrocytes) characterized by increased secretion of pro-inflammatory factors, disrupted mitochondria function, and a failure to nurture neurons or eliminate foreign agents, which collectively leads to neurotoxicity and neuroinflammation (12, 13). In neurodegenerative disorders such as Alzheimer’s (AD), Parkinson’s (PD), Huntington’s diseases (HD,) and Amyotrophic lateral sclerosis (ALS), astrocytes were shown to be activated by fragmented extracellular mitochondria released from microglia stimulated by the damage-associated molecular patterns (DAMP) secreted by diseased neurons (14). A key process in the regulation of mitochondria fragmentation is phosphorylation of dynamin-related protein – 1 (Drp1), and its subsequent relocation to the mitochondrial outer membrane, where it binds to the Fis1 receptor (10). In rodent models of moderate types of brain injuries, blocking excessive Drp1-mediated mitochondria fission with an inhibitor of Drp1 catalytic activity (Mdivi-1) ameliorated early neuronal damage progression and improved behavioral outcomes (15).

While it is known that brain injury induces mitochondria dysregulation in rodents (15), and microglia-astrocyte crosstalk is neurotoxic in neurodegenerative disorders (14), the role of glial mitochondria fragmentation to brain injury-induced secondary neurodegeneration remains to be established. Here we determined crosstalk and cell-type-specific contributions of neurons, microglia, and astrocytes to dysregulated mitochondria-driven neurodegeneration after moderate injury in a human 3D triculture tissue model of the brain. Our results demonstrated that microglia play a pivotal role in the progression of secondary damage post-contusion through increased pathological mitochondria fragmentation and its metabolic dysregulation.

## Results

### *In vitro* 3D human triculture brain tissue model fabrication and characterization

To determine glial cell-specific contributions to injury-induced neuronal network degeneration progression, we developed a 3D *in vitro* triculture human tissue model of a contusion injury, composed of induced neural stem cells, human primary astrocytes, and an HMC3 microglia cell line, or iPSC (induced pluripotent stem cells) derived microglia from two healthy donors (YZ1 and ND41866*C). These human tricultures (neuron, astrocyte, and microglia culture (NAMc)) were generated by seeding neurons, astrocytes, and microglia at a ratio of 2:0.5:0.1 million, respectively, in 3D silk scaffolds and then enveloping them in a collagen type I hydrogel for sustained long-term growth (Fig. 1a, Supplementary Fig. 21). Culture conditions, such as viability, neuronal network density, glial cell-specific markers, were optimized for the survival of all three cell types when combined in the tricultures (Supplementary Fig 1-3, 21). Confocal image analysis, gene and protein expression studies of tri-cultures demonstrated upregulation of quintessential cell-type specific transcription factors (NeuN for mature neurons, Sox9 for astrocytes, and PU.1 for microglia) concomitant with an increase in density of beta-3-tubulin (Tuj1) positive neuronal networks (Fig. 1 b-e, Supplementary Fig. 2-3). Importantly, astrocytes or microglia in di-cultures with neurons have no effect on neuronal network density compared to neuronal monocultures (Supplementary Fig. 3). These results suggest that interactions between the multiple cell types in a 3D environment support cellular growth and maturation (Fig. 1 b-e).

**Fig. 1.**
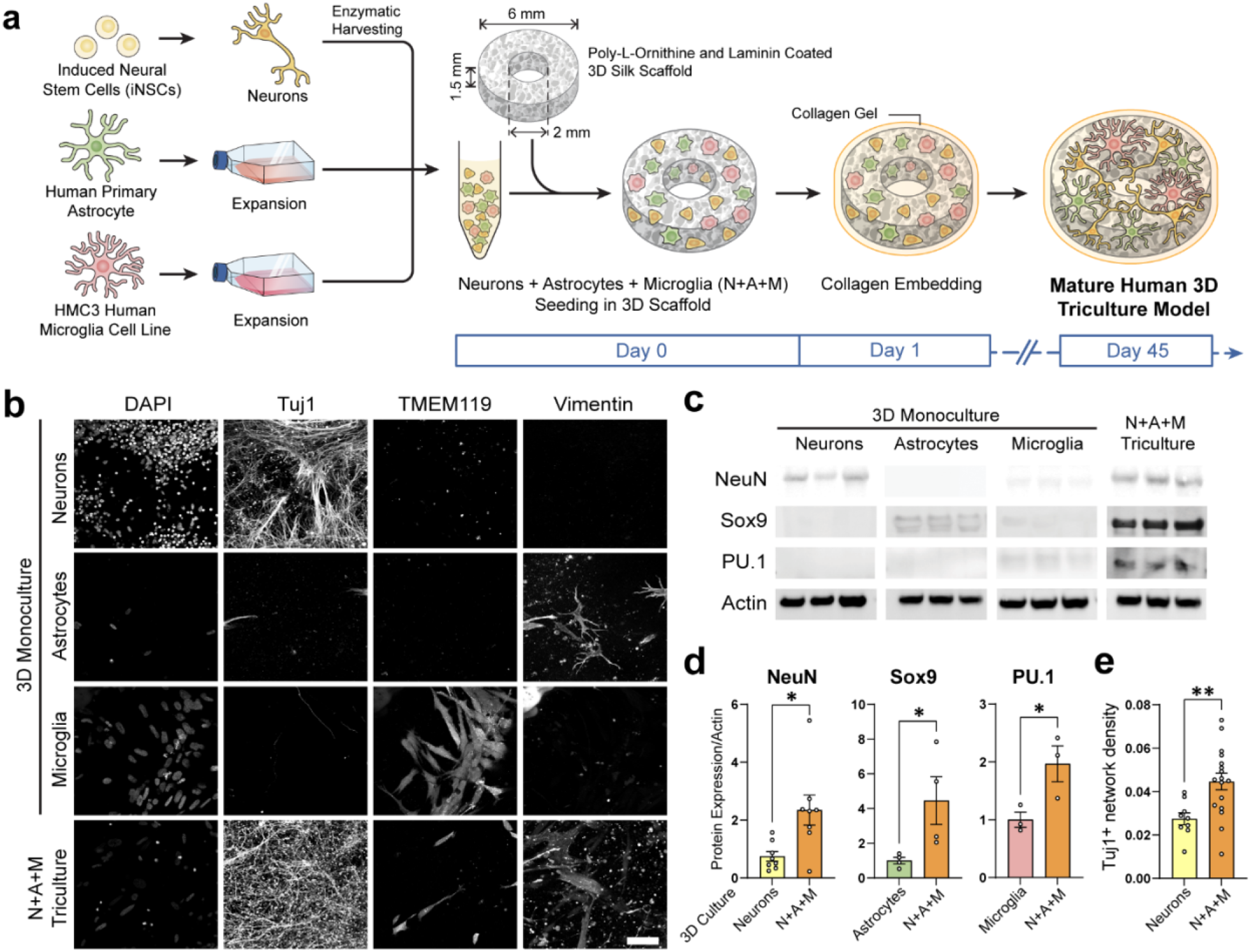
Validation and characterization of human *in vitro* 3D triculture model composed of neurons, microglia, and astrocytes. **a**, Schematic representation of the preparation of 3D tricultures. **b**, Representative images of neuron (Tuj1), microglia (TMEM119), and astrocyte (vimentin) specific cytoplasmic markers in monocultures and tricultures. **c**, Representative images of Western blots stained with antibodies recognizing cell-specific transcription factors (NeuN (neuronal), Sox9 (astrocytic), PU.1 (microglial), or actin as a loading control. **d**, Quantification of the Western blots showing results normalized to actin expression, and **e**, Tuj1 neuronal network density quantified from confocal z-stack images. Scale bar: 50µm. **(d)** Means ± SEM of n=2-4 scaffolds per condition. *, ** indicates a significant difference (p<0.05, 0.01 respectively; t-test with Welch’s correction was used to determine the differences between control and experimental groups). All experiments were replicated at least three times.

Gene expression analysis in tricultures demonstrated a high level of neuronal quintessential (TUBB3, PAX6, SYN1) and maturation markers (MAP2, RBFOX3), cortical layer-specific genes (RELN, CUX1, POU3F2, BCL11B, FOXP2), excitatory and inhibitory neuronal markers (ATP12A2, TH, SLC6A1, ACHE, GRIN2B, ISL2) (Supplementary Fig. 4a); high levels of astrocytes (SOX9, AQP4, GFAP, S100B, ALDOC, SLC1A2, ALDH1A1, ALDH1L1) and microglia pan markers (HEXB, IRF8, RUNX1, SPI1, TAL1); and low to undetectable levels of pluripotency (OCT4, NANOG) or brain endothelial cells markers (VWF, CDH5M, CD34) (Supplementary Fig. 4b) (16). Microglia-specific genes were expressed at a lower level, due to a small percentage of these cells present in the tricultures (4%) in comparison to neurons (77%) or astrocytes (19%). To further analyze the state of neuronal maturation in the 3D triculture in vitro model, the gene expression was compared to the earlier published genetic footprint of iPSC-derived neurons in monocultures, co-cultures with astrocytes, and adult human neurons (Supplementary Fig. 5 a-d) (16). Gene expression of markers associated with adult neurons, and maturation of iPSC-derived neurons (in particular, in co-culture with astrocytes) were expressed at comparable levels in the human 3D in vitro triculture model (Supplementary Fig. 5 b-d).

To ensure a non-inflamed state of glia in co-culture conditions, the production level of inflammatory cytokines, chemokines, and growth factors was evaluated (Supplementary Fig. 6). Among 24 profiled markers, the level of IL-8 chemokine was significantly higher in co-cultures conditions, comparing to neuronal monocultures, and in triculture, IL-8 was not different from NA or NM di-cultures. Other markers were equal between all groups, indicating a homeostatic and stable environment.

### Injury-induced neuronal network degeneration in the 3D human triculture brain tissue model is blocked by P110 peptide, a selective inhibitor of excessive mitochondrial fission and fragmentation

To determine cell-type specific contributions of mitochondrial fragmentation in contusion-induced neurodegeneration, the 3D cultures of different compositions were exposed to P110 peptide, a selective inhibitor of excessive mitochondria fission and fragmentation (Fig. 2 a, b) (17). The P110 treatment protocol was chosen based on the previously published work (14). The P110 peptide selectively inhibits GTPase activity of Drp1 protein (and its translocation to mitochondria) without affecting the activity of fusion proteins (OPA1 mitochondrial dynamin-like GTPase (OPA1) and mitofusin 1 (MFN1)) (17). Human 3D tri-, di-, and monocultures were subjected to contusion injury using a controlled cortical impactor (CCI, here and after “CCI” in figures refers to the model of induced injury) at 45-50 days post-seeding, using parameters established previously *in vitro* and *in vivo* (Fig. 2 b, c) (18). Structural (neuronal network density), biochemical (LDH), and molecular (glutamate) markers of contusion injury were monitored at 4 different time points within 2 weeks of injuries (8, 24, 48 hours, and 14 days) (Fig. 2-4, Supplementary Fig. 7-13). Contusion injury resulted in the release of two brain injury-associated biomarkers, lactate dehydrogenase (19) and glutamate (20), in all mono-, di- and tricultures within 48hr (Supplementary Fig. 7). Only in the triculture conditions did the increased release of LDH and glutamate persisted for a prolonged time (14 days), after 40% drop at earlier time points 24 and 48hr. P110 treatment had no effect on the glutamate and LDH release, which is consistent with other drug treatment protocols, where neuronal protection was not associated with LDH or glutamate concentration (data not shown). Next, we evaluated injury-induced neural network degradation (Fig. 2 d-e, Supplementary Fig. 8-10) by quantitative confocal microscopy analysis developed previously in our group (18).

**Fig. 2.**
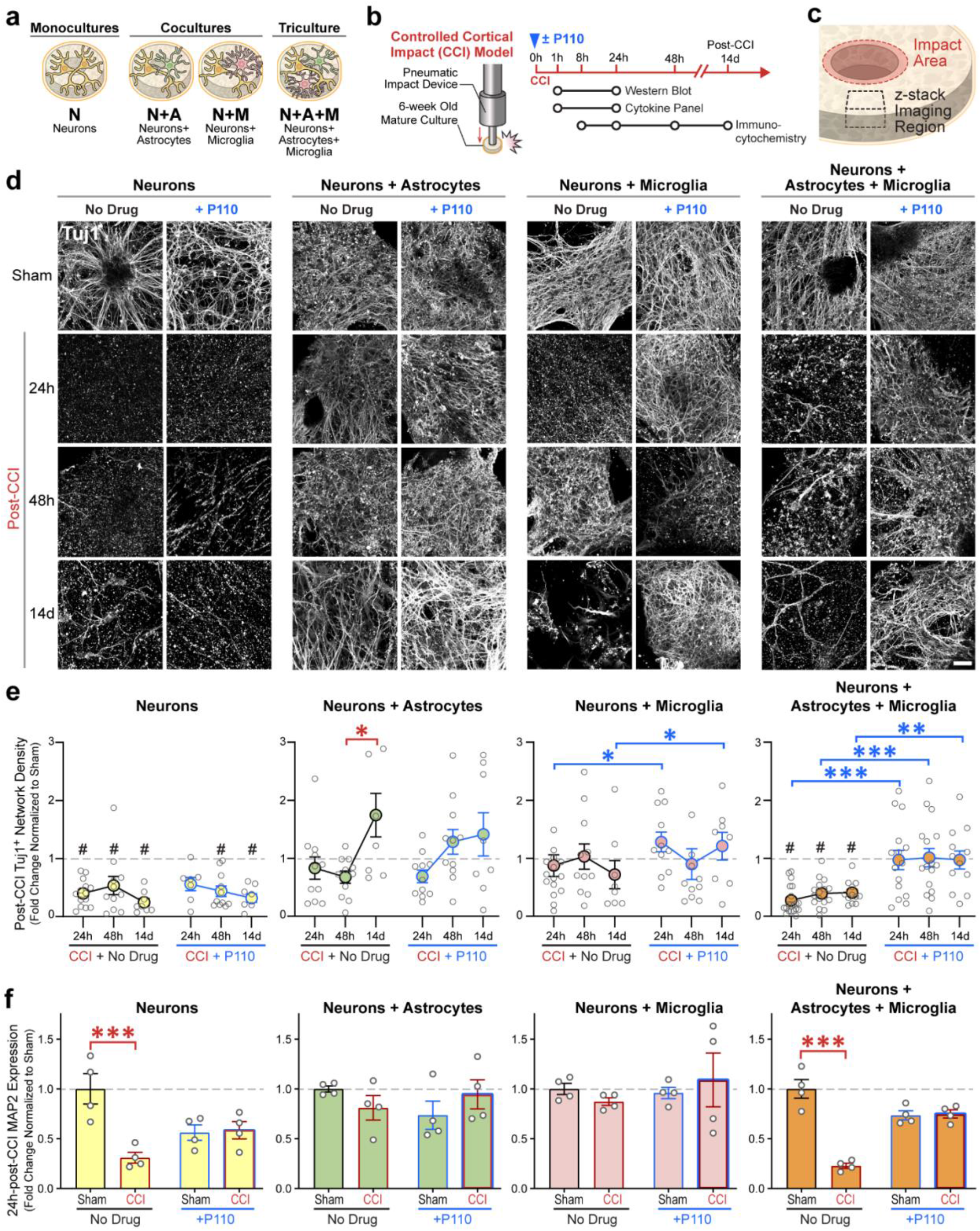
Neuronal network degeneration in tricultures with microglia and astrocytes, but not neuronal monocultures is reversible with mitochondria fission inhibitor P110. **a**, Schematic representation of experimental groups. **b**, Contusion (Controlled cortical impact model (CCI)) injury experimental design. **c**,Schematic representation of the injury area, and image acquisition. **D**, Representative images of Tuj1 neuronal staining at 24, 48 hours, and 14 days after injury in N, NA, NM, and NAM groups. **e**, Quantification of Tuj1 neuronal network density in control and P110 treated groups of N, NA, NM, and NAM at three different time points (24, 48 hours, and 14 days). **f**, Quantification of MAP2 protein Western blots at 1 and 24 hours after injury in all groups. Proteins were isolated from entire scaffolds, without separating the injured area from the penumbra. Data presented in **e** and **(f)** mean ±SEM of three independent experiments with n=2-4 scaffolds per condition. *, **, *** indicate significant differences (p<0.05, 0.01, 0.001 respectively; two-way ANOVA (analysis of variance) (with Tukey’s post-hoc test) between control and experimental groups). Scale bar: 50 µm.

In Nc, NAc, and NMc the neural network in the directly impacted area (3mm in the center of the scaffold, Supplementary Fig. 8) degraded by 90% within 24 hours (p<0.01) in the absence or presence of P110. In NAMc however, we have observed intact neuronal network density in the directly impacted area within 24 hours in P110 treated group, but not the control (Supplementary Fig. 8-9). In the adjacent to the injury site (outside the 3mm of the impacted zone), the proximal area of the neuronal monocultures (Nc), network density declined from 24 hours up to 14 days after the injury (from 50% degeneration at 24 and 48 hours to 60% at 14 days, p<0.01) (Fig. 2 d-e; Supplementary Fig. 10). In contrast, di-cultures of neurons with astrocytes (NAc) or microglia (NMc) showed (90%) network survival after injury, up to 48 hours, that decreased by 30% in NMc at 14 days. Injured NAMc showed the largest (80%) drop in network density at 8 hours after the impact and it remained 70-80% degraded up to 14 days (p<0.05). With P110 treatment, the network was preserved in the NA, NM, and NAM cultures at all time points. However, in neuronal culture (Nc), the drug protected the network from degradation compared to sham only at the 24-hour time point (Fig. 2d, e; Supplementary Fig. 10), however, there was no difference compared to injured samples without drug treatment. The above results were corroborated with Western blot analysis of MAP2 protein, the dendritic marker of mature neurons, 24 hours after injury (Fig. 2f). A reduction of MAP2 was observed in Nc and NAMc conditions (60% drop, p<0.01), with unchanged levels in the P110 treatment group.

Next, we looked at the activation of the necroptotic pathway, a known mechanism of neuronal death post-controlled cortical impact injury in rodents (21). We have confirmed necroptotic death execution in Nc and NAMc (the groups that showed neuronal network degeneration) through phosphorylation of mixed lineage kinase (pMLKL), known as necroptotic executor (Supplementary Fig. 11). While the early marker of necroptotic death – phosphorylation of receptor-interacting serine/threonine-protein kinase 3 (RIP3) has not been detected (Supplementary Fig. 11). At last, P110 peptide had no significant effect on the progression of necroptotic death, indicating that P110 acts downstream of necroptosis activation and progression.

In summary, in the absence of glial cells, neurons underwent rapid network loss through a P110 independent process, implicating a mitochondria fission independent mechanism. Microglia (short-term) and astrocytes (short and long-term) in di-cultures with neurons had protective properties on network degeneration progression after contusion injury. However, neurons were unable to withstand injury when grown together with microglia and astrocytes. P110 drug treatment prevented neuronal network degradation in NAM tri-cultures, suggesting a mitochondria fission-dependent process.

### Mitochondria fission is associated with neuronal network degradation in tricultures after contusion injury

To confirm that mitochondria fission was associated with neuronal network degeneration the mitochondria aspect ratio (AR, a measure of mitochondria width over length; Fig. 3 a, b), the total amount of TOMM20 (translocase of outer mitochondrial membrane 20) mitochondria, and the expression levels of fission (Fig. 3 c-e) and fusion markers (Supplementary Fig. 12) were analyzed. Nc showed a decreased amount of TOMM20 positive mitochondria with a high fragmentation rate of TOMM20 24 hours after injury, which was partially rescued in the presence of the P110 peptide. However, the decrease in the AR was not changed after the P110 treatment. In control or P110 treated injured NMc, NAc, and NAMc there was no decline in the amount of TOMM20 positive mitochondria (Fig. 3 c). In the injured NAc and NMc with/without P110 treatment, the aspect ratio and the level of mitochondria fission, and fusion remained equal to shams. While the total amount of mitochondria remained unchanged in NAMc, the mitochondria underwent a significant transition to the fragmented state after contusion, showing a decreased mitochondria aspect ratio, and increased expression of pDRP1 (dynamin-related protein-1 phosphorylated at Serine 616) and Fis1 (mitochondria fission 1 protein), two factors responsible for mitochondria fission (Fig 3). However, expression of mitofusin 1,2 (MFN 1,2), a fusion marker was not decreased (Supplementary Fig. 13). Mitochondria fragmentation was reversed in the P110 treated tricultures (NAMc), showing sham levels of pDRP1 (Fig. 3 d) and Fis1 expression (Fig. 3 e), and sham level of aspect ratio (Fig. 3 a, b). These results indicated the gain-of-toxic function of microglia and astrocytes in tricultures, that induced mitochondria fragmentation-associated neurodegeneration progression after injury.

**Fig. 3.**
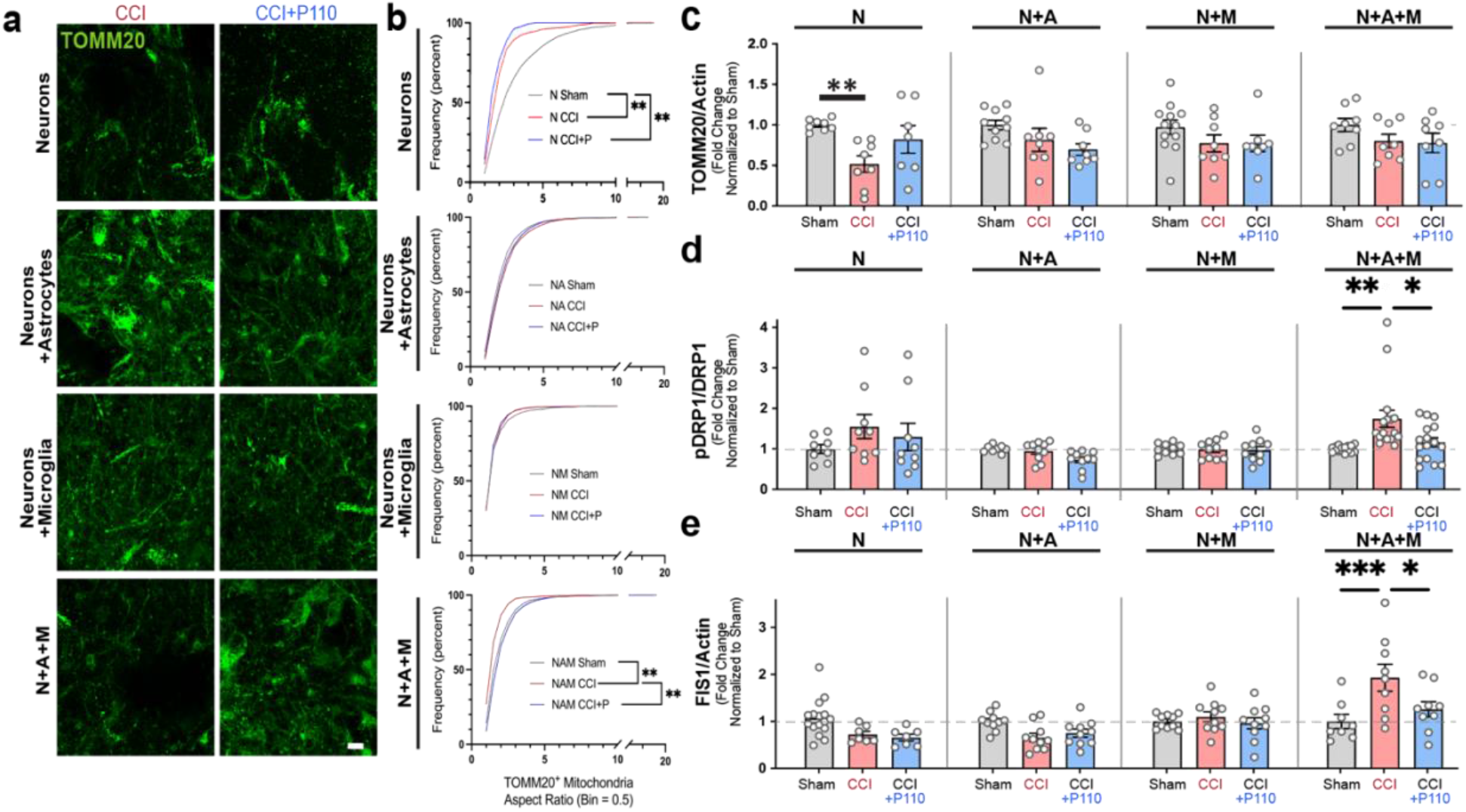
Injury-induced neurodegeneration in tricultures, but not neuronal monocultures was associated with increased Drp1-Fis1 driven mitochondria fission. **a**, Representative images of TOMM20 mitochondria at 24 hours after contusion in N, NA, NM, and NAM groups. **b**, Frequency distribution (%) of TOMM20-positive mitochondria aspect ratio in control and P110 treated groups of N, NA, NM, and NAM. **c**, TOMM20 protein quantification by Western blot 24 hours after injury in all groups. **d, e**, pDRP1/DRP1, and FIS1 protein quantification by Western blot 24 hours after the injury in all groups. Proteins were isolated from entire scaffolds, without separating the injured area from the penumbra. Data presented in **(b-e)** mean ±SEM of three independent experiments with n=2-4 scaffolds per condition. *, **, *** indicates significant difference (p<0.05, 0.01, 0.001 respectively; two-way ANOVA (with Tukey’s post-hoc test) between control and experimental groups).

### Increased mitochondria fission post-contusion in tricultures is associated with induced neuroinflammation

Human and rodent *in vivo* studies of brain injury showed an increased risk of acute and chronic neuroinflammation, often associated with worsening the patients’ outcomes (1, 5, 22). Having linked an increased mitochondria fission to secondary neuronal network degeneration progression in tricultures, we next evaluated if mitochondria fragmentation influenced acute neuroinflammation in NAM tricultures 24 hours after the injury (Fig. 4, Supplementary Fig. 13).

Increased expression of key A1 astrocyte phenotype activators (12, 14, 23) interleukin-1 alpha, beta (IL-1α, IL-1β), tumor necrosis factor-alpha (TNFα), and ciliary neurotrophic factor (CNTF) was mitigated in the presence of P110 after injury. Additionally, C-C motif chemokine ligand 2 (CCL2), interleukin-6 (IL-6), interferon-gamma (IFNγ), and other acute inflammatory markers (Supplementary Fig. 8), were upregulated after injury and mitigated with P110 treatment. TNF-α and Fas ligand (FasL) has been linked to worsened outcomes after CCI in mice and proposed as contributors to secondary damage progression (24). Here, we found upregulated expression of both TNFα and FasL, which was rescued with P110 treatment. Some TBI biomarkers, such as brain-derived neurotrophic factor (BDNF), glia cell-derived neurotrophic factor (GDNF) (22), granulocyte-macrophage colony-stimulating factor (GM-CSF) (25), and S100 calcium-binding protein B (S100B) (26) were upregulated after injury, and the response was mitigated in the presence of P110 (Fig. 4, Supplementary Fig. 8).

**Fig. 4.**
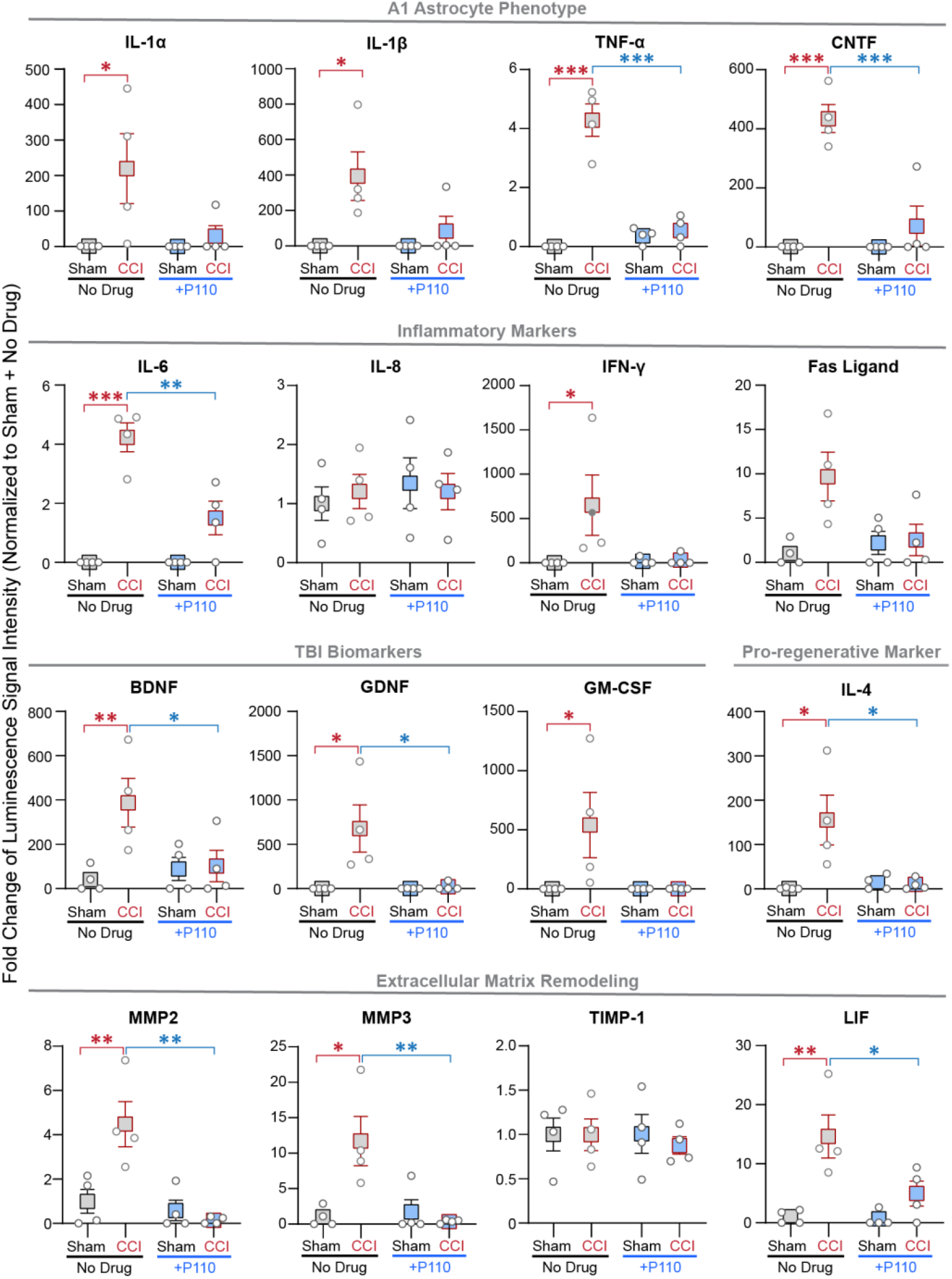
Increased secondary neurodegeneration was associated with neuroinflammation in tricultures 24 hours after injury. Quantification of cytokine array for A1 associated cytokines: IL-1α, IL-1β, TNFα, and CNTF; Inflammatory markers: IL6, IL8, IFNγ, Fas ligand; Neurotrophic and Growth factors: BDNF, GDNF, GM-CSF; Pro-regenerative marker: IL-4; Extracellular matrix remodeling proteins: MMP2, MMP3, LIF, and TIMP1 secreted 24 hours after injury in triculture NAM cultures with/-out P110 treatment. Data presented are the mean ± SEM of four independent experiments with each data point being the average for n=2-4 scaffolds per condition. *, **, *** indicate significant differences (p<0.05, 0.01, 0.001 respectively; two-way ANOVA (with Tukey’s post-hoc test) between control and experimental groups).

The brain undergoes changes in the extracellular environment after injury (7). Here we found an increase in matrix metalloproteinases-2 and -3 (MMP2 and MMP3) with the injury that was mitigated with P110 treatment, while tissue inhibitors of metalloproteinases (such as TIMP1) remained unchanged in control and P110 treated cultures (Fig. 4). Some TBI markers, such as interleukin-8, -18 (IL-8, IL-18), vascular endothelial growth factor (VEGF) and T-cell directed CC Chemokine (TARC) remained unchanged in response to contusion or in the presence of P110 (Fig. 4, Supplementary Fig. 8) (27). Additionally, some known pro-regenerative markers such as interleukin -4, -10 (IL-4, IL-10), and transforming growth factor beta (TGFβ) were upregulated after contusion but remained at sham levels in the P110 treated group. Together these results indicated that mitochondria fission played a significant role in regulating neuroinflammation post-injury.

### Dysfunctional mitochondria released from microglia contributed to injury-induced neurodegeneration and neuroinflammation progression post-contusion

We further investigated the cell-specific contributions of mitochondria dysregulation to secondary damage progression after injury through isolation and transfer of mitochondria (Supplementary Fig. 14 a, b) from injured cultures to monocultures that were not exposed to contusion (Fig. 5 a-g; Supplementary Fig. 14). Conditioned media or mitochondria isolated from injured neurons were not sufficient to induce neurodegeneration after transfer to naïve neuronal monocultures (N-N transfer; N-Nt) (Supplementary Fig. 15); however, if conditioned media from injured neurons was transferred to naïve microglia, and then extracellular mitochondria from these cultures were added to naïve neuronal monocultures, we observed about 50% of network degradation (p<0.05; N-M-Nt) (Supplementary Fig. 14 c, d). This effect was exacerbated by 75% of network degradation (p<0.01; N-M-A-Nt, please see Fig. 5a for the schematic of the experiment) when naïve neurons were treated with mitochondria isolated from astrocyte cultures (that were treated for 24h with extracellular mitochondria from microglia, activated by conditioned media from injured neurons) (Fig. 5 b-c). Increased network degeneration was concomitant with a significant drop in neuronal mitochondria aspect ratio (Fig. 5 d-e). To evaluate if there is a cell type-specific contribution to injury progression, NAc were treated with dysfunctional mitochondria released from activated microglia, and NMc with astrocytic mitochondria (Supplementary Fig. 15). We observed massive neuronal network degeneration only in NAc treated with mitochondria from microglia, and the degeneration was prevented with P110 treatment (Supplementary Fig. 15).

**Fig. 5.**
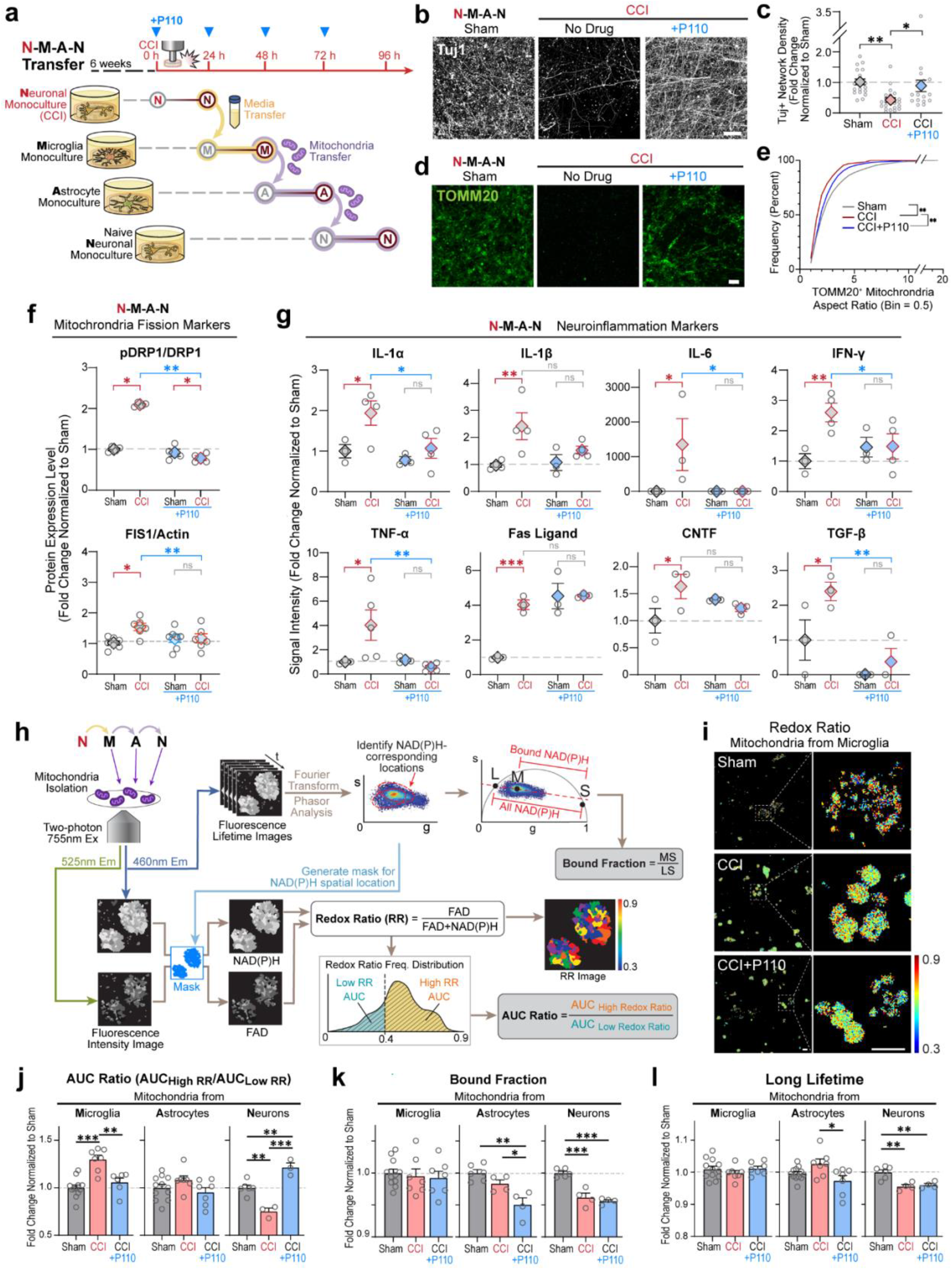
Mitochondria dysregulation in microglia was crucial for secondary neurodegeneration and neuroinflammation progression post-contusion injury. **a**, Schematic representation of transfer experiment from neurons to microglia to astrocytes, and neurons with/without P110 compound performed at each transfer step. Representative images of **b**, Tuj1, and **d**, TOMM20 staining of naïve neurons treated for 24 hours with mitochondria isolated from astrocytes activated with mitochondria isolated from microglia that were treated with conditioned media collected from 24h injured neurons with quantification of **c**, Tuj1 positive neuronal network density, and **e**, TOMM20 positive mitochondria aspect ratios. **f**, Quantification of the pDRP1/DRP1 and Fis1 proteins expression in N-M-A-N control and P110 treated groups. **g**, Inflammatory cytokine secretion in N-M-A-Nt control and P110 treated groups. **h**, Schematic representation of the metabolic image analysis. **i**, Representative redox ratio maps of mitochondria isolated from microglia activated by conditioned media from injured neurons. Quantification of **j**, redox ratio, **k**, NADH bound fraction, and **l**, a long lifetime in live mitochondria isolated from microglia, astrocytes, and neurons treated with mitochondria isolated from injured neurons, activated microglia, or astrocytes. Proteins were isolated from entire scaffolds, without separating the injured area from the penumbra. Data presented in **(c, e, f, g, j, k, l)** mean ±SEM of n=2-10 scaffolds per condition. *, **, ***, **** indicates significant difference (p<0.05, 0.01, 0.001, 0.0001 respectively; one-way or two-way ANOVA (with Tukey’s post-hoc test) between experimental groups). Experiments were replicated at least three times. Scale bar in **b, d**: 50 µm; in **i**: 10 µm.

To examine the role of crosstalk between astrocytes and microglia in mitochondrial-induced neuroinflammation and neurodegeneration, we focused our study on the injured neuron to microglia to astrocyte to neuron transfer (N-M-A-Nt), please see Fig 5a for experimental design (Fig. 5 b-g). We first tested if inhibition of mitochondrial fission was able to rescue the neuronal damage (Fig. 5 b, c). P110 treatment restored the AR of mitochondria TOMM20 networks and prevented neuronal network damage (Fig. 5 d, e). Conditioned media from injured neurons induced pDRP1 and Fis1-associated mitochondria fission in microglia cultures (N-**M**-A-Nt, second transfer step, Fig. 5a), which was prevented by P110 treatment. Similar results were obtained in naïve neurons treated with extracellular mitochondria isolated from astrocytes (N-M-A-**N**t, fourth transfer step, Fig. 5a), but not in astrocytes treated with extracellular mitochondria isolated from microglia (N-M-**A**-Nt, the third transfer; Supplementary Fig. 14). We thus confirmed the contribution of extracellular mitochondria to fission associated secondary neurodegeneration progression in neurons after brain injury (Fig. 5 f).

To correlate the effects of mitochondria dysregulation on neuroinflammation in tricultures (NAMc) (observed in Fig. 4 and Supplementary Fig. 16), the expression of the same panel of cytokines and chemokines was evaluated in transfer experiments. Similar patterns were detected between N-M-A-Nt transfer (Fig. 5g, Supplementary Fig. 16) and NAMc culture after injury (Fig. 4): elevated expression of pro-inflammatory markers IL-1α, IL1-β, IL-6, IFN-γ, TNF-α, FasL, MMP2, and MMP3. The increase of these markers was mitigated in the presence of the P110 peptide. Our findings indicated that excessive mitochondria fission in microglia was crucial to the progression of neurodegeneration, and the effect was exacerbated with microglia-astrocyte neuroinflammatory crosstalk.

We next characterized the function of mitochondria released from neurons, astrocytes, and microglia in transfer experiments (N-M-A-Nt). The number of released mitochondria (DNA quantification) at each transfer step and its physiological function -ATP production and oxygen consumption (Supplementary Fig. 17) were examined. Activated microglia and astrocytes (by conditioned media from injured neurons, or mitochondria released from microglia) showed an increased amount of released mitochondria with decreased levels of ATP production and oxygen consumption (Supplementary Fig.17). The P110 treatment mitigated the effect of injury on the number of released mitochondria and their function.

The number of mitochondria released by activated naïve neurons was not different from the sham; however, with lower ATP production rate, and increased oxygen consumption. P110 treatment induced increased secretion of mitochondria with a lower oxygen consumption rate (Supplementary Fig. 16). Next, we examined the total amount of released mitochondria over time (from M to A to N) in sham and injury separately and discovered an overall increase in the number of mitochondria at the last transfer step in sham groups, however, in injured group, the number of mitochondria was significantly decreased over time, indicating potential re-uptake or degradation of mitochondria by neuronal cells, but not glial. The mitochondria amount was comparable to sham when samples were treated with P110 peptide (Supplementary Fig. 18).

To corroborate the results of mitochondria characterization with biochemical methods, real-time imaging of isolated mitochondria was used to assess metabolic function based on the endogenous two-photon excited fluorescence of NAD(P)H (reduced nicotinamide adenine dinucleotide and reduced nicotinamide adenine dinucleotide phosphate) and FAD (flavin adenine dinucleotide). The intensity images were used to calculate the redox ratio, defined as FAD/(NAD(P)H+FAD), while lifetime images were analyzed to extract the fraction of NAD(P)H in bound form and it’s corresponding (long) lifetime (Fig. 5 h-l)(28). Extracellular microglial mitochondria showed an increased redox ratio following injury, which was mitigated in the presence of P110 (Fig. 5 i, j); in astrocytic mitochondria, the redox ratio remained at the sham level in both control and P110 groups (Fig. 5 j), while neuronal mitochondria showed a significant drop in redox ratio after injury in the untreated group and increase in P110 treated group (Fig. 5j). The bound fraction of NADH was consistently lower in P110 treated astrocytes and neurons (Fig. 5 k). In the injury group, it was lower only in neurons and no change was observed in the microglia group. The long NAD(P)H lifetime was lower in the P110 group in astrocytes and neurons (Fig. 5 l), and for the untreated injury group only in neurons (same as what was observed with bound fraction measurement); no change was detected in microglia following contusion injury with or without P110 treatment. Our data demonstrated that extracellular mitochondria derived from microglia that had been activated by conditioned media from injured neurons are dysfunctional and trigger a cascade of metabolic changes in astrocytic and neuronal mitochondria.

### Mitochondria dysregulation-induced neurodegeneration and inflammation are independent of the microglia cell source

To test if the fragmented mitochondria-driven neurodegeneration is inherent to microglia, and not just the HMC3 cell line, we replicated the most critical experiments with iPSC-derived microglia from two healthy donors (YZ1 and ND41866*C) (Fig. 6-7, Supplementary Fig. 19-31), and further characterized the genetic response of 3D in vitro tricultures to contusion injury, using mRNA sequencing analysis. Microglia were differentiated following an established protocol (29, 30) and characterized for specific marker expression after each round of differentiation (Supplementary Fig. 19). We prepared 3D triculture tissues with iPSC-derived microglia following the protocol described above (Fig. 1, Supplementary Fig. 1-3) and performed viability staining at specific timepoints to ensure healthy, symbiotic growth of the three cell-types in NAMc (Supplementary Fig. 20).

**Fig. 6.**
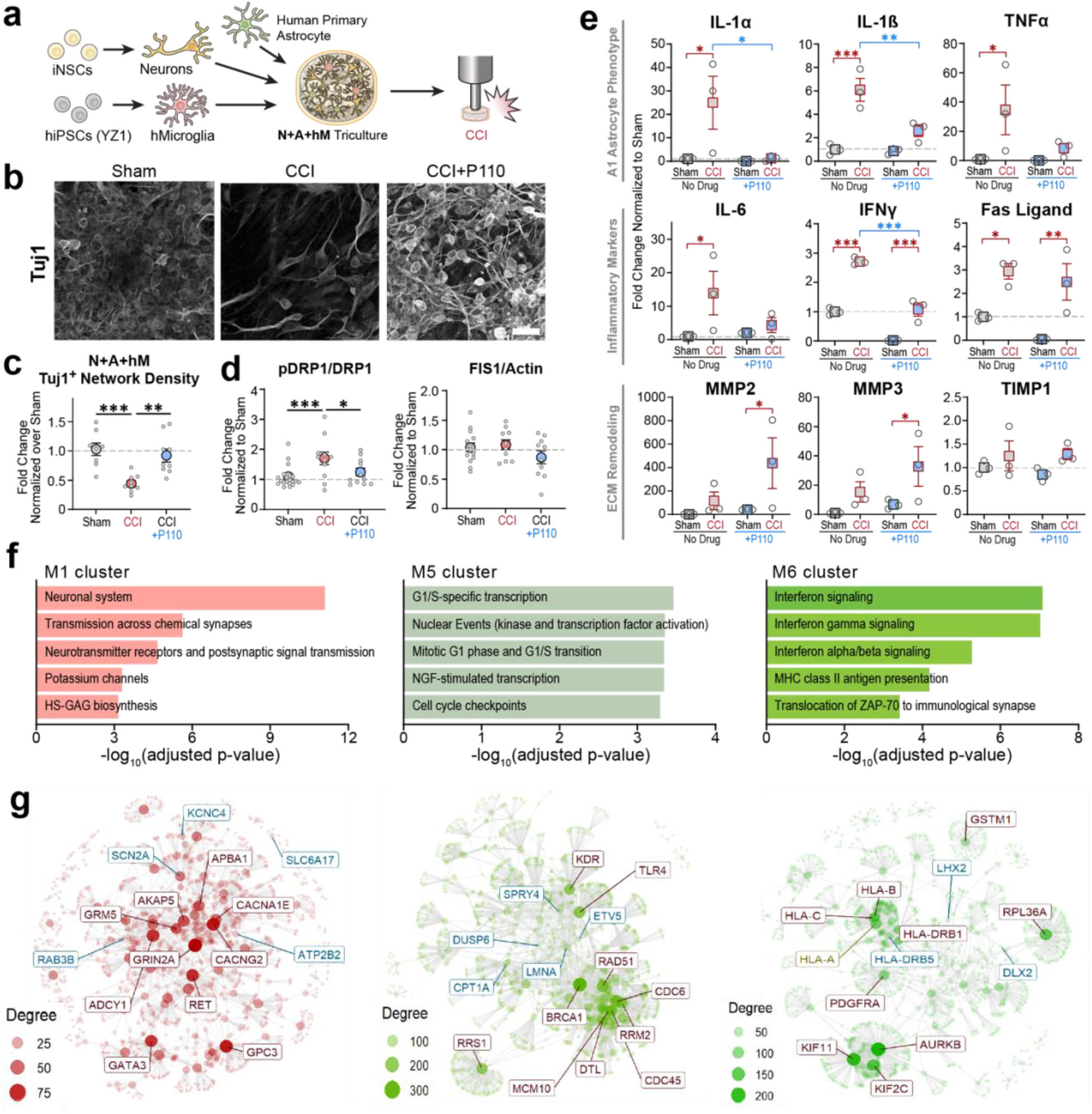
Dysfunctional mitochondria released from iPSC-derived microglia (YZ1 line from a healthy donor) induced neurodegeneration and neuroinflammation progression 24 hours after contusion injury. **a**, Schematic representation of the experimental design. **b**, Representative images of Tuj1 network density in tricultures with iPSC derived microglia. **c**, Quantification of the Tuj1 neuronal network density in control and P110 treated groups of NAMc. **d**, Mitochondria fission associated pDRP1 and FIS1 proteins expression in control and P110 treated NAM cultures. Proteins were isolated from entire scaffolds, without separating the injured area from the penumbra. **e**, Inflammatory cytokines secretion post-injury in YZ1 triculture groups. **f**, Gene set enrichment analysis revealed distinct expression modules associated with M1 - neuronal systems (negatively associated with a contusion in CTRL; rescued with P110); M5 - replication and repair (positively associated with a contusion in CTRL and P110); M6 - interferon signaling (positively associated with a contusion in CTRL; unchanged in P110). **g**, Critical regulators of molecular mechanisms associated with M1, M5, and M6 modules. The full map of 7 detected modules and corresponding pathways can be found in Supplementary Fig. 25, 26, and 29. Data presented in **(c-e, g-i, k-m)** mean ±SEM of n=2-10 scaffolds per condition. *, **, *** indicates a significant difference (p<0.05, 0.01, 0.001 respectively; two-way ANOVA (analysis of variance) or one-way ANOVA between control and experimental groups) (with Tukey’s post-hoc test). Experiments were replicated at least three times; metabolic images were replicated two times. Scale bar: 50 μm in **b, f, h**; 10 μm in **j**

**Fig. 7.**
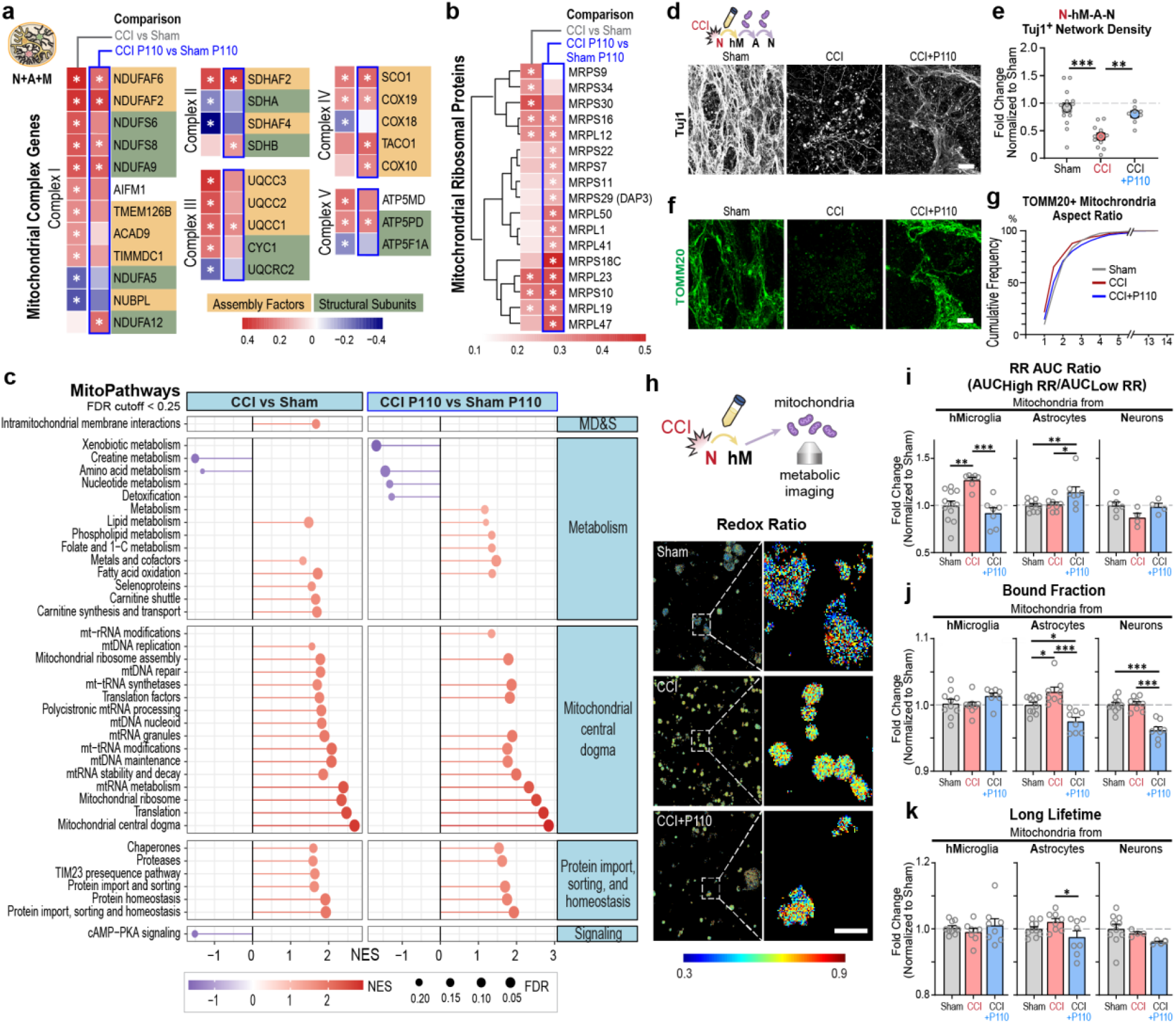
Contusion injury induced dysregulation of intra- and extracellular mitochondria. Differentially expressed genes (DEGs) associated with **a**, mitochondria complex genes; and **b**, mitochondria ribosomal proteins were detected (abs(Log2(Fold Change))> 0.585 and q < 0.01) in human 3D in vitro triculture model composed of induced neural stem cells derived neurons, primary astrocytes, and iPSC-derived microglia from two healthy donors YZ1 and ND41866*F (n=6) and summarized in over-representation analysis of **c**, mitochondria pathways. **d, f**, Representative images and **e-g**, quantifications of Tuj1 neuronal and TOMM20 mitochondria networks in naïve neurons treated with mitochondria isolated from astrocytes activated with mitochondria isolated from YZ1 microglia that had been treated with conditioned media collected from 24h injured neurons. **h**, Representative redox ratio maps of mitochondria isolated from microglia that were activated by conditioned media from injured neurons. Quantification of **i**, redox ratio, **j**, bound fraction, and **k**, a long lifetime in live mitochondria isolated from iPSC microglia, astrocytes, or neurons following N-M-A-Nt experimental design (Fig. 5a). Data presented in **(e, g, i-k)** mean ±SEM of n=6-10 scaffolds per condition. *, **, *** indicates a significant difference (p<0.05, 0.01, 0.001 respectively; two-way ANOVA (analysis of variance) or one-way ANOVA between control and experimental groups) (with Tukey’s post-hoc test). Experiments were replicated at least three times; metabolic images were replicated two times. Scale bar: 50 μm in **d, f**; 10 μm in **h**

After 6 weeks, the 3D tricultures went through the injury protocol and the effect of contusion injury was observed after 24 hours (Fig. 6; Supplementary Fig. 14-15). The neuronal network degraded by 50% (p<0.05) in cultures containing microglia cells from either donor and the effect was reversed in the presence of P110 (Fig. 6 a-c; Supplementary Fig. 21 a, b). The degradation of the neuronal network was associated with increased mitochondria fission marker expression, pDRP1 (both lines), however, we did not observe an increase in Fis1 expression in either of the lines (Fig. 6 d; Supplementary Fig. 21 c). In NAMc made with either of the iPSC microglia lines, the expression of pDRP1 was reversed in the presence of P110. Unlike in tricultures with the HMC3 cells, levels of Mitofusin 1,2 protein were downregulated in the iPSC tricultures after injury, while the treatment with P110 peptide was not significant from either sham or injury group without treatment (Supplementary Fig. 22), suggesting decreased fusion. Similar to HMC3 tricultures (Fig. 4-5), we observed mitochondria fission-induced neuroinflammation; A1 astrocyte-associated cytokines IL-1α, IL-1β, and TNFα were upregulated in untreated injured cultures and downregulated to the levels of a sham when treated with P110 (Fig. 6 e, Supplementary Fig. 23, 24). Increased levels of ECM remodeling markers MMP2 and MMP3 were reduced by P110 in injured HMC3 but not in YZ1 or ND41866*C tricultures (Fig. 6e, Supplementary Fig. 24). mRNA sequencing cluster analysis revealed significant changes in ECM assembly and function, further suggesting ECM contribution in injury progression (Supplementary Fig. 25-26). Expression of the apoptosis marker FasL was similarly upregulated in all microglial tricultures; however, the levels were reduced by P110 treatment only in HMC3 tricultures. mRNA sequencing cluster analysis revealed enrichment of type I interferon response and major histocompatibility complex (MHC) class I antigen presentation in response to contusion injury (Fig. 6 f, g – cluster 6, Supplementary Fig. 25, 27), which correlates with genetic signature observed in rodent studies of brain injuries (31). We next looked at the expression of glia-specific gene-contributors to interferon type I response activation, known to be dysregulated post fluid percussion injury model in rodents (25 top DEGs), and detected similar changes in our contusion human *in vitro* injury model (Upregulated - SLFN5, OAS3, PARP12, TRIM25, CCL2, PARP9, ISG15, PAPR14, IFIT2, 3, IFI16, OAS1, SP100, GDP2, EIF2AK2, IFIH1, ATP10A, MLH1; downregulated - MTHFR, C3, XDH, IRGM, SDC3, NAMPT) (Supplementary Fig. 27). We have found greater transcriptional activation of microglia specific markers, compared to astrocytes, which has been shown *in vivo* rodent injury studies as well (31). Some of the markers were not detected in our model (some examples are IFI209, OASL2, IFI27I2A, IRF9, and H2-Q4), due to species differences, and limitations of our model (lack of macrophages, blood vessels, and other factors) (31). Among upregulated genes in astrocytes were GBP3, IFIT3; and downregulated MAFB. Microglia genes upregulated were PARP14, PHF11, IFIT3, IFI16, MF213, MLH1, SRP54, MYO1E, and downregulated SDC3 (Supplementary Fig. 27).

At last, we have discovered that astrocytes specific pan-markers were significantly downregulated, together with neuron-specific markers (Fig. 6 f-g, cluster M1, Supplementary Fig. 25, 28-29), while microglia markers were mostly unchanged, or upregulated (Supplementary Fig. 28). Indicating, the potential death of neurons and astrocytes, but not microglia. Additionally, cluster analysis of cell cycle gene expression revealed activation of cell cycle and DNA repair progression (Fig. 6 f, g – cluster 5).

### Contusion injury induced mitochondria intra- and extracellular changes in human 3D triculture model of brain injury

To further characterize mitochondria health post-injury, the expression profile of mitochondria-specific markers was examined (Fig. 7 a-c). mRNA expression analysis detected increased expression of genes associated with mitochondria oxidative phosphorylation pathway (complex I and III), while P110 treatment mitigated the increase in some of these markers (Fig. 7 a). Unlike OXPHPOS markers, the genes responsible for mitochondria ribosomal protein expression (names) were significantly upregulated in the P110 treated group (Fig. 7 b), compared to control injury or sham groups. At last, a significant downregulation in the expression of glycolysis genes was observed in both control and P110 treated groups (Supplementary Fig. 30). Overall, a significant decrease in the mitochondria metabolism of creatinine and amino acids was observed regardless of the drug treatment, however in the control condition, increased expression of carnitine metabolism genes was detected, potentially providing an alternative path for NADH generation for OXPHOS pathway (32). Increased expression of the mitochondria genes responsible for mitochondria DNA replication and repair in the control group is indicative of damage progression, while in both control and P110 treated groups increased transcription and mitochondria protein assembly genes may indicate potential mitochondrial degeneration.

At last, to confirm microglial mitochondria fission contribution to neurodegeneration, the transfer experiments were replicated in iPSC microglia-based models (Fig. 7 d-k; Supplementary Fig. 21 e, f, 31). Structural damage and mitochondrial fragmentation were observed in neurons that received extracellular mitochondria from microglial exposed to conditioned media isolated from injured neurons (50% of network degraded in both iPSC lines; N-M-Nt). P110 treatment counteracted these effects (Supplementary Fig. 31). Extracellular mitochondria isolated from astrocytes (after they were treated for 24 hours with microglial mitochondria activated with conditioned media from injured neurons, please see Fig. 5a for experimental design) exacerbated neurodegeneration in transfer experiments with the ND41866*C line to 75% of network degradation (N-M-A-Nt; Supplementary Fig. 21 e, f) and remained similar in YZ1 cultures (50% of network degradation) (Fig. 7 d-g) These effects were associated with increased levels of mitochondria fragmentation in both lines (Fig. 7 f, g; Supplementary Fig. 21 e, f).

We examined the metabolic profile of mitochondria released from neurons, astrocytes, and iPSC microglia (YZ1 line) after exposure to conditioned media from injured neurons (Fig. 7 h-k; N-M-A-Nt). Similar to transfer experiments with the HMC3 line, we observed an increased level of redox ratio in microglial mitochondria following the transfer of conditioned media from injured neuronal cultures. P110 treatment mitigated this increase. Mitochondria isolated from astrocytes that had been activated with microglial mitochondria showed an increased redox ratio for the P110 injury group. No change was detected after the last transfer step to neurons (Fig. 7 i). The NADH bound fraction was consistently lower in P110 treated astrocytes and neuronal groups, with no changes in the microglia group (Fig. 7 j). However, the astrocytes control injury group showed an increase in NAD(P)H bound fraction, and no change was observed in the neuronal control group, unlike in HMC3 experiments. The long lifetime measurements were similar between the HMC3 and the YZ1 line. Together these findings confirmed cell-line independent effects of microglial mitochondria fission on secondary neurodegeneration and neuroinflammation progression after contusion injury.

## Discussion

The temporal profile of neuroinflammation and neurodegeneration after brain injury has been studied in animal models of TBI, as well as in brain tissue, cerebrospinal fluid, and plasma samples from TBI patients (5). While rodent models are helpful in exploring behavioral changes and multicellular interactions between the central and peripheral nervous systems, limitations include innate differences between rodents and humans (such as white/grey matter ratio, microglia activation early after TBI in rats vs persistence for years in humans, brain edema lasts for a few days in rats vs weeks in humans, genetics) (33), screening limits due to animal use and costs, and the lack of options for long-term studies (years). Human clinical samples are a useful source for the detection of biomarkers and understanding of late stages of disease progression; however, they cannot be used to study molecular triggers and disease onset or to perform exploratory research of underlying mechanisms. To overcome these limitations, we developed a preclinical 3D human tissue model that would permit insights into molecular and cell-specific contributions to the progression of TBI. The *in vitro* model consists of multiple neural cell types (neurons, astrocytes, and microglia) for the study of cell-specific contributions to injury progression. The engineered tissues withstand the mechanical insult and recapitulated TBI primary injury hallmarks (cell death, excitotoxicity, acute neuroinflammation) and secondary damage progression (neurodegeneration, chronic neuroinflammation, cell death).

### 3D bioengineered triculture human tissue model of brain injury recapitulates native injury progression in rodents and humans

When compared to corresponding 3D monocultures, the triculture conditions enhanced the expression of cell-specific transcription factors (neuronal NeuN, astrocytic Sox9, and microglial PU.1) and increased neuronal Tuj1 positive network density (Fig.1). Improved cell maturation may be due to cellular crosstalk that stimulates the secretion of (neuro)trophic factors (23) or enhances neuronal survival through (physical) support from glia (e.g., glutamate reuptake and synaptic pruning)) (7). We showed that our tissue model replicated many critical aspects of human brain injuries (1, 19, 20, 34), such as progressive neuronal network damage (Fig. 2, 6; Supplementary Fig. 7-10, 21), glutamate release (Supplementary Fig. 7), increased membrane permeability, as witnessed by LDH release into the media, (Supplementary Fig. 7), and secretion of pro-inflammatory and neurotrophic factors (IL-1α, IL-1β, IL-6, TNF-α, BDNF, NGF, as well as others) (Fig. 4, 6; Supplementary Fig. 13, 21, 23-24). Additionally, we observed some differences in the inflammatory markers’ secretion and the response to P110 treatment between tricultures with iPSC microglia and HMC3 cells, indicating the importance of the genetic contribution to the severity of injury progression and potential drug treatments.

### Crosstalk between glial cells drives neuroinflammation and secondary damage progression after injury through activation of excessive mitochondria fission

*In vivo* rodent experiments suggest that dysfunction in mitochondria leads to brain tissue loss and poor behavioral outcomes (e.g., spatial and motor learning, anxiety, fear, activity) in moderate and severe TBI (35). However, the role of mitochondrial dysregulation in damage progression after brain injury, particularly the contributions of multicellular crosstalk, remained unexplored. We observed a high rate of neurodegeneration in neuronal monocultures that was accelerated in tricultures (compare 8 h timepoints, Supplementary Fig. 10), suggesting a toxic gain-of-function of glial cells, possibly through the secretion of inducers of the A1 astrocytic phenotype (12) (IL-1β, IL-1α, C1q, TNFα) and other pro-inflammatory molecules (Fig. 4 and Supplementary Fig. 13). The underlying mechanisms of injury-induced neurodegeneration are in part dependent on mitochondria acquiring a pathological state, characterized by excessive fission, that can be rescued by a specific inhibitor (P110). In contrast, the loss of TOMM20 staining coupled with the absence of fission markers and a lack of response to P110 in neuronal monocultures suggests that a failure in mitochondria function is associated with neurodegeneration in the absence of glia. Altogether, these results suggest that neurodegeneration in neuronal monocultures is independent of mitochondrial fission pathways, while neurons in crosstalk with glial cells experience mitochondria fission-dependent damage (Fig. 2-3).

### Contusion injury induces metabolic changes in the mitochondria

Human brain mitochondria undergo substantial metabolic changes after the injury, including a short-term (12 hours) increase in glycolysis, followed by prolonged glycolytic depression (up to a few weeks) (36); strong decrease in the oxygen metabolism through oxidative phosphorylation cycle in the long-term (weeks)(37, 38); high lactate to pyruvate ratio in the brain (an indicator of the patient’s poor outcome) (39). In our in vitro model of a contusion injury, we observed decreased expression of glycolysis-associated genes, however, further experiments are required to confirm if glycolysis is compromised. Intriguingly, despite of decreased level of glycolysis (the main producer of pyruvate needed for the Krebs cycle to generate NADH, that fuels the OXPHOS), expression of genes associated with OXPHOS was increased 24 hours after the injury, potentially using pyruvate produced through increased metabolism of carnitine, instead of classic glycolysis cycle. An alternative source of pyruvate could be lactate released by astrocytes in response to injury and converted into pyruvate by LDH (40). In our system, we observed a decrease in glycolysis, the classic pathway of pyruvate generation, however, we detected only in triculture conditions a decrease in LDH after the initial contusion-induced spike (Supplementary Fig. 7), which potentially could explain the increase in OXPHOS to compensate the lack of glycolysis.

### Dysfunctional mitochondria released from injury-activated microglia are a driving force for secondary neurodegeneration progression after TBI

Rodents and 2D *in vitro* models have shown an association between fragmented mitochondria and the progression of several neurodegenerative disorders in addition to TBI (14, 15). In 2D models of Amyotrophic lateral sclerosis (ALS), Alzheimer’s (AD), Parkinson’s (PD), and Huntington’s disease (HD), these effects can be mimicked in neural cell cultures by exposure to mitochondria released from the glia (14), however, the link between cell-specific mitochondria dysregulation to neuronal damage has never been shown in TBI. Our results indicated that mitochondria released from microglia (cell line or iPSC-derived) that had been activated by conditioned media from injured neurons were a driving force of secondary neural damage progression (N-M-Nt). The mechanism of mitochondrial release and the factors released by injured neurons to stimulate microglia remain to be characterized but could involve active signaling, dying cells, and DAMPS (14, 41). In transfer experiments including astrocytes (N-M-A-Nt), induction of mitochondria fission and neuroinflammation was coupled to accelerated neurodegeneration, suggesting a feedback loop of injury-induced damage progression and crosstalk between the three cell types. Label-free, optical assessments of mitochondrial metabolism suggest that extracellular microglial mitochondria shift to a more oxidized state, possibly as a result of oxidative stress (Fig. 5 h-l and 7 h-k) (42-44). In P110 treated astrocyte and neuronal groups, the combined decrease in the NAD(P)H bound fraction and long lifetime in extracellular mitochondria may be attributed to enhanced utilization of NADPH in the glutathione pathway to combat injury-induced oxidative stress (45).

A major advantage of our bioengineered 3D tissues over other *in vitro* models (e.g., 2D cultures, brain-on-chip) is the ability to inflict physiologically relevant mechanical injuries and monitor the primary and secondary injury progression over an extended time frame (46). Cells in this 3D system remain functional for over 2 years, providing opportunities to understand cellular and genetic contributions to injury progression and pharmacological interventions (47, 48). The key finding of the present TBI study is the pivotal role of glial-neuronal crosstalk in the progression of secondary damage and neuroinflammation and the role of mitochondria function therein. These data suggest that selective inhibition of pathological mitochondrial fission mediated by Drp1-Fis1 could be a potential therapeutic target for TBI patients.

## Disclaimer

The opinions and assertions expressed herein are those of the authors and do not necessarily reflect the official policy or position of the Uniformed Services University, or the Department of Defense.

## Acknowledgments

The authors thank the NIH (R01AG061838 and P41EB027062), the Department of Defense Technology for the Warfighter Program (HU0001-20-2-0015), the NIH Research Infrastructure grant (NIH S10OD021624) for the purchase of the microscope, Dr. Yu-Ting L. Dingle, of Pipette & Stylus LLC, for assistance with illustrations and figures, Dr. Dana Cairns, for supplying hiNSC cells, Rushil Bakhshi for testing the mitochondria isolation protocol. And Dr. Dustin Snapper for help with the discussion of the results.

## Authors contribution

V.L., M.J.W., A.J.S., I.G., T.J.F.N., D.L.K conceived the project, designed, interpreted the experiments, and wrote the manuscript. V.L. conducted all *in vitro* experiments. V.L. isolated mitochondria, Y.Z. collected metabolic images of mitochondria, Y.Z. and I.G. performed the analysis. V.L., Y.M., M.C, A.M., A.S.-Z. expanded cells for triculture seeding experiments, V.L. and M.C. differentiated and analyzed iPSC-derived microglia with M.B.-J. guidance, M.F.K., Y.M., A.S.-Z. performed sample maintenance (media changes). V.L., Y.M., M.F.K., V.K.R., and M.R.S. conducted western blots. V.L., Y.M., M.F.K., V.K.R., M.R.S., A.B., A.S.- Z. performed biochemical assays, immunofluorescent staining, and with A.M. fabrication of silk scaffolds. V.L. collected all immunofluorescent images. V.L. and M.F.K. performed and analyzed protein arrays. V.L. and M.F.K. performed mRNA isolation, N.J.F. performed mRNA sequencing analysis.

## Materials and methods

### 3D Silk Scaffold Preparation

#### Silk Processing and Scaffold Fabrication

Silk scaffolds were prepared from silk fibroin following a previously established protocol (Rockwood et al., 2011). *Bombyx mori* cocoons were boiled for 30 minutes in a sodium carbonate solution (0.02M) to isolate silk fibroin and remove sericin. The fibroin fibers were dried overnight in a fume hood. The following day, the silk fibers were dissolved in 9.3 M lithium bromide (Sigma) by immersion for 4 hours at 60°C. Dialysis of the silk in deionized water at room temperature was performed for 3 days in 3,500 molecular weight cut-off dialysis tubing (Invitrogen) to remove the lithium bromide from the silk. The resulting silk solution was centrifuged twice at 9,000 rpm for 20 minutes and a strainer with 100 µm pores was used to filter out the remaining debris. Then, 1 mL of the silk solution was dried at 60°C overnight to determine the weight/volume (w/v) concentration. The silk solution concentration was adjusted to 6 mg/mL and combined in a 10 cm dish with 400 to 500 µm sodium chloride (Sigma) particles at a ratio of 1:2 (v/w) and left for 2 days at room temperature to trigger pore formation and crystallization for stability in aqueous solution and to further refine the 3D sponge structures, the solution was incubated for one hour at 60°C. After separating the sponge from the dish, residual sodium chloride was removed by dialysis in deionized water for 2 days at room temperature with 6 water changes total. The sponges were cut into individual donut-shaped scaffolds using biopsy punches (Integra) with a 6 mm outside and 2mm inside diameter. The scaffolds were trimmed to a height of 1.5 mm using scissors and a razor blade. Scaffolds were then autoclaved in excess deionized water for 20 minutes with the liquid cycle, cooled to room temperature, and then processed for coating as described below. Silk scaffolds were used for experiments within 3-months of preparation.

#### Coating Silk Scaffolds with Extracellular Matrix

Scaffolds were coated with 10µg/mL poly-ornithine (PLO) (Sigma, cat.no. A004C) and 5µg/mL laminin (Fisher, cat.no. 501003381) to aid in cell adhesion. Up to 50 scaffolds were placed in a single well of a 6-well plate using fine forceps and incubated overnight at 37°C in 7 ml of a 10ug/ml PLO coating solution in distilled water. The next day, the PLO was aspirated and washed three times with PBS with 5-minute incubations between washes. The scaffolds were then incubated overnight at 4°C in 0.5 mg/mL 100µL laminin stock aliquots with Dulbecco’s Modified Eagle Medium/Nutrient Mixture (DMEM/F12, Invitrogen) phenol red-free media (PRF) in a 1:100 ratio and then the scaffolds were retained in the laminin solution. The scaffolds were incubated at 37°C for 3 hours before use.

### Cell Culture

#### Mouse embryonic fibroblasts (MEFs)

Following ATCC guidelines, MEFs were used as a feeder layer for induced neuronal stem cells. MEFs were used at passages 1-3. DMEM (Invitrogen) supplemented with 10% Fetal Bovine Serum (FBS) (Invitrogen) and 1% Antibiotic-antimycotic (Anti-Anti) (Invitrogen) were used to maintain the MEFs. The media was changed every 3 days. Cells were passaged at 80% confluency. In brief, cells were washed with PBS and then treated with 0.25% Trypsin-EDTA solution for 3 minutes at 37°C. Once the Trypsin-EDTA (Invitrogen) was deactivated with MEF media, the cells were centrifuged for 5 minutes at 1,000 rpm. The MEFs were further split from 1 dish into 3 dishes (15cm^2^).

As a feeder layer, the MEFs were allowed to reach 100% confluency and then inactivated with 10 µg/mL Mitomycin C (Sigma, M4287) for 3 hours, followed by three washes with PBS. After the last PBS wash, media was added to the plates, and the cultures were maintained until the hiNSC seeding (7 days maximum).

#### Induced human neural stem cells (hiNSCs)

The hiNSC cell line was previously generated from dermis-derived human fibroblasts, isolated from human neonatal foreskin fibroblasts through a direct reprogramming (49). hiNSCs were handled following a previously established protocol (3). The cells were maintained in media composed of KnockOut DMEM (Invitrogen) supplemented with 1% GlutaMax (Invitrogen); 20% KnockOut Serum Replacement (Invitrogen); 1% Anti-Anti (Invitrogen); 0.2% β-mercaptoethanol (Invitrogen); 800 µL of 10 µg/mL basic fibroblast growth factor (Invitrogen) added proportionally to aliquots of media upon use. Media was changed daily. In brief, hiNSCs were expanded and subcultured on top of inactivated mouse embryonic fibroblasts. When cells reached 70-80% confluency, they were lifted from the plates via incubation with TrypLE solution for 1 min at 37°C, followed by quenching with 6mL of hiNSCs expansion media. Cells were then collected and pelleted by centrifugation at 3,000 rpm for 2 minutes. For further expansion, pellets were gently disrupted with a 10 ml pipette, and the colony suspension was transferred to an inactivated MEFs layer; for 3D seeding, the pellet was rigorously pipetted to obtain single-cell suspensions. For expansion, hiNSCs colonies were seeded from 1 dish into 20 (15cm^2^).

#### Human primary astrocytes

Human primary astrocytes (Sciencell research laboratories, cat.no. 1800) were expanded on 20 µg/mL poly-L-lysine (Sigma, P4832) coated surfaces up to passage 5 following established protocols from the manufacturer. Astrocytes were maintained in astrocyte media supplemented with astrocyte growth factors (1%), Fetal Bovine Serum (2%), and Anti-Anti (1%) from Sciencell research laboratories (cat.no. 1801); media was changed every 3 days. In brief, when astrocytes were 70-80% confluent, they were lifted from the plate via incubation in 0.25% Trypsin/EDTA for 3 minutes at 37°C, followed by quenching with astrocyte complete media. Next, the cell suspension was collected from the plates and pelleted by centrifugation at 1,000 rpm for 5 minutes. Cells were re-seeded at 350,000 cells/cm^2^.

#### HMC3 microglia cell line

The HMC3 microglia cell line was obtained from ATCC and maintained following established protocols (50). In brief, microglia were expanded in 15 cm^2^ dishes in EMEM media (ATCC) supplemented with 10% Fetal Bovine Serum (Invitrogen), and 1% Anti-Anti (Invitrogen). The media was changed every 3 days. When cells reached 70-80% confluency, they were lifted from plates with 0.25% Trypsin/EDTA incubation for 3 minutes at 37°C. After quenching with microglia complete media, cells were pelleted with centrifugation at 1,000 rpm for 5 minutes. Cells were seeded at 350,000 cells/cm^2^.

#### Induced pluripotent stem cells

The hiPSC line ND41866*C was obtained from the Coriell Biorepository (Camden, NJ, USA). The YZ1 hiPSC line was obtained from Guisepena Tesco courtesy of the University of Connecticut-Wesleyan Stem Cell Core (UCSCC, Farmington, CT, USA). YZ1 cell line is derived from healthy female lung fibroblasts, and the ND41866*C cell line is derived from healthy male skin fibroblasts.

#### Induced pluripotent stem cells derived microglia

Two iPSC lines from healthy donors (YZ1 and ND418664) were differentiated into microglia following a previously established protocol (29, 30). iPSCs were expanded in colonies on a Matrigel layer in mTeSR1 Embryonic stem cell media (StemCell Technologies, cat.no. 85850) as directed by batch instructions from the manufacturer. Each differentiation round required one vial of frozen iPSCs. The iPSCs were thawed and incubated in mTeSR1 media supplemented with 0.5 µM Rock Inhibitors (StemCell Technologies, cat.no. 72304) for 24 hours at 37°C, with daily media changes until iPSC colonies were near each another. To lift the colonies, 2 mL of ReLeSR (StemCell Technologies, cat.no. 05872) solution was applied for 30 seconds at room temperature, followed by 3 minutes incubation at 37°C after the ReLeSR was aspirated. Media was added to the plates and the side of the plates was gently tapped to detach the cells. After collecting and centrifuging the cells at 300xg for 5 minutes, the cells were split into a 1:6 ratio and re-plated in mTeSR1 media. The following day, the mTeSR1 media was replaced with E8 media (StemCell Technologies, cat.no. 05990), preferable for microglia differentiation. A second splitting followed the same passaging protocol summarized above following the addition of the E8 media at least once before differentiation.

To initiate iPSC differentiation to hematopoietic stem cells, the iPSCs were seeded in 6-well plates coated in phenol-red free Matrigel with hematopoietic differentiation media (StemCell Technologies, cat.no. 05310). The media was changed following the manufacturer’s protocol. After 11 days, early progenitors of hematopoietic cells were collected and counted, and then seeded in 6-well plates coated with phenol-red free Matrigel (∼400,000 cells/well); 2 mL of microglia differentiation media composed of DMEM-F12 phenol red-free media supplemented with 2% B-27 and 0.5% N2 supplements, 1% Glutamax, 5µg/mL Insulin, 2% Insulin-Transferin-Selenite, 1% Non-essential amino acids, and 400µM monothioglycerol was added to each well with freshly infused 100 ng/mL interleukin-34 (IL-34), 25 ng/mL macrophage colony-stimulating factor (MCSF), and 50 ng/mL transforming growth factor beta (TGFβ) growth factors. Then 1 ml of fresh media was supplemented with IL-34 (R&D systems, cat. No. 5265-IL-010), MCSF (Invitrogen, cat.no. PHC9501), and TGFβ (Fisher, cat.no. 5018578) growth factors every other day. On days 12 and 24 of growth, 5 mL of the 6 mL of media in each well was removed and centrifuged to remove the media with accumulated waste. The cells were resuspended in 1 mL of media and replated back into the respective well. On day 25 of growth, the media was supplemented with 100 ng/mL fractalkine (Invitrogen, cat.no. 10636H08H50) and 100 ng/mL OX-2 membrane glycoprotein (CD200) (R&D systems, cat.no. 27712CD050) growth factors in addition to the previously described growth factors.

Hematopoietic stem cells and iPSC-derived microglia were characterized for cell-specific marker expression after each round of differentiation. Live cells were stained with a panel of markers: clusters of differentiation 43, 34, 235, 41, 45, 117 (CD43, CD34, CD235, CD41, CD45, c-kit), triggering receptor expressed on myeloid cells 2 (TREM2), transmembrane protein 119 (TMEM119), and C-X3-C motif chemokine receptor 1 (CX3CR1) and imaged with a Leica SP8 Confocal microscope (Leica Microsystems).

#### 3D in vitro human triculture brain tissue model fabrication

The system for the generation of 3D human tricultures was modified from our earlier published protocols of 3D rat, mouse, and human neuronal monocultures (18, 51, 52). When hiNSCs, primary astrocytes, and HMC3s reached nearly 100% confluency, they were collected and pelleted as described in the individual culture methods, and then combined to achieve the following ratio: 2:0.5:0.1 million neurons, astrocytes, and microglia, respectively. The same ratio was used for cocultures for the respective cell types. Immediately before seeding the triculture cell solution, previously prepared PLO- and laminin-coated silk scaffolds were placed in 96-well plates and a vacuum manifold was used to remove excess liquid. Calculations were performed so that each 40µL of cell suspension contained 2:0.5:0.1 million neurons, astrocytes, and microglia, respectively. Then 40µL of the cell suspension was pipetted onto the semi-dried scaffolds and incubated for 30 minutes at 37°C to allow the cells to attach. After this, 150µL of Neuronal media was added to each well. Neuronal media was composed of NeuroBasal medium supplemented with 2% B-27, 1% Anti-Anti, 1% Glutamax, and 1% astrocytes growth factors. The scaffolds were placed in a tissue culture incubator (37°C, 5% CO_2_ in a humidified atmosphere). The following day, the cell-seeded scaffolds were transferred to new 96-well plates. To each scaffold, 100µL of collagen type I solution (Corning or R&D systems, 3 mg/mL with a pH adjusted to 7.0-7.2 with NaOH) was added, and then incubated for 30 minutes at 37°C to allow the collagen gel to crosslink. Next, 150µL of neural media was added to all scaffolds, followed by incubation for 24 hours at 37°C. The following day, the brain-like tissues were moved into 48-well plates with 1 ml of Neuronal media in each well. Media was changed every fourth day until the tissues were used for injury studies, as controls, or for analysis 6 weeks after seeding.

### 3D Model Injury and Drug Treatment

#### Contusion injury (Controlled cortical impact model (CCI))

At six weeks, the 3D brain-like tissues were placed on a flat weigh boat and subjected to contusion injury using a pneumatic cylinder with a 5 mm flat tip impactor at a velocity of 6m/s, penetration depth 0.6mm, and a dwell time duration of 200 ms (18). Sham tissues were handled in a similar fashion to the injured tissues but without receiving the contusion injury. Following injury or sham protocol, the tissues were incubated at 37°C in 5% CO_2_ in a humidified atmosphere until the indicated time marks for analysis of lactate dehydrogenase (LDH) and glutamate release in the medium, cell death, neural network stricture, western blot, and mitochondria health.

#### Drug treatment

Dynamin-related protein 1 (Drp1) – mitochondria fission 1 protein (Fis1) peptide inhibitor P110 was purchased from Tocris Bioscience (cat. No.68971) (17). For each independent experiment, a new vial of P110 was freshly reconstituted in deionized water at 1 mg/mL. Immediately, following injury or sham treatment, tissues were treated with either 2.4µL of: a) P110 (reconstituted) to achieve 1 µM concentration or b) control treatment (deionized water). Tissues were then incubated at 37°C in 5% CO_2_ and a humidified atmosphere until the indicated time marks for analysis of LDH and glutamate release in the medium, cell death, neural network structure, protein expression, and mitochondrial health.

### Cell Health and TBI Progression Analysis

#### Lactate dehydrogenase activity assay

The cellular viability of the brain-like tissues was tested using a lactate dehydrogenase assay (LDH) (Sigma, cat.no. MAK066-1KT) following the manufacturer’s protocol. LDH assays measure LDH activity via 1,4-dihydro nicotinamide adenine dinucleotide (NADH) formation from nicotinamide adenine dinucleotide (NAD). Cell culture media was collected at the indicated experimental time points and stored at −80°C until all media samples were ready for analysis. Then, 50µl of LDH assay supernatant was added to each sample and the NADH levels were read with a SpectraMax M3 TECAN plate reader, at 450nm. LDH activity was calculated in the following three steps: a) a calibration curve was generated for each individual plate read from standards; b) absorbances were compared to the calibration curve to calculate NADH levels; c) LDH activity was based on the reaction time and the applied dilution.

#### Glutamate assay

Free glutamate in the media was measured with a Glutamate assay kit (Abcam, cat.no ab83389) following the manufacturer’s protocol. Cell culture media was collected at the indicated experimental time points and stored at −80°C until analysis. Then 50µl of sample supernatant and 50µl of glutamate reaction mix (enzyme mix) were combined. The reaction between the enzyme mixture and glutamate substrate was measured with a SpectraMax M3 TECAN plate reader, at 450nm. Calibration curves were generated from standards for each individual plate. Glutamate concentrations were derived via comparison to the calibration curve.

#### Viability staining

3D tissue viability was checked at 2 and 6 weeks after seeding following the established Calcein AM (Invitrogen, C1430) staining protocol. In brief, scaffolds were transferred to clean 48 well plates prior to staining, and 300-500 µl of 2 µM Calcein AM solution was added to each 3D tissue. The samples were then incubated for 30 minutes at 37°C in the dark. When 30 minutes passed, samples were immediately moved for imaging with Leica SP8 Confocal at 37°C using 494/517nm excitation/emission wavelengths.

#### Immunofluorescence Staining and Analysis

A solution composed of 4% sucrose, and 4% paraformaldehyde (PFA, Electron Microscopy Sciences) in PBS was used to fix the 3D brain-like tissues. After fixing, the tissues were washed five times with PBS and then permeabilized for 1 hour with 0.2% TritonX-100 supplemented with 4% goat serum (Thermo Fisher). The permeabilization solution was also used to generate a primary antibody solution. Tissues were incubated in the primary antibody at 4°C overnight, then washed with gentle shaking in PBS five times (5 minutes each wash). After washes tissues were incubated with secondary antibodies diluted in PBS for 1 hour. 4’,6-Diamidino-2-Phenylindole (DAPI) was applied to the tissues for 5 minutes following secondary antibody staining. Five additional PBS washes were used to dispose of any unbound antibodies or DAPI. For all figures, fluorescent image stacks of the stained 3D brain-like tissues were acquired on a Leica SP8 FLIM confocal microscope (Leica Microsystems, Fluotar VISIR 40x/0.95 WATER objective, 290.62 µm x 290.62 µm x 41.81 µm, each maximum projection had 100 z-steps with 0.422 µm in depth each). Images in the figures represent maximum intensity projection and were collected with the same PMT gain settings and laser power between at least three independent experiments. Tuj1 neuronal network density was analyzed using a custom MATLAB code published earlier in our group (18).

Primary antibodies included: anti-TOMM20 (ab186735 rabbit; 1:1000); anti-TOMM20 (ab56783 mouse; 1:1000); anti-beta-tubulin III (ab18207 rabbit; 1:1000); anti-beta-tubulin III (ab78078 mouse; 1:1000); purchased from Abcam.

Secondary antibodies included: goat anti-mouse IgG1, IgG2 (A-21240, A21135), anti-rabbit (A11037) or anti-chicken (A21449) Alexa 488 and 568, 647 (1:500; Thermo); goat anti-chicken (A10040) Alexa 546 (1:500; Life Technologies).

#### Mitochondria Aspect Ratio analysis

The ratio of mitochondria length vs width (Aspect ratio) was measured using available macro from NIH ImageJ software, following published protocol (53). In brief, maximum projection images of TOMM20 positive mitochondria, acquired with Leica SP8 confocal (1024 × 1024 pixels, Fluotar VISIR 40x/0.95 WATER objective) were analyzed by ImageJ Marco. The measured AR for each mitochondrion in at least 4 images (individual samples) was plotted as a distribution graph for comparison with other groups.

#### Western Blot

To extract cellular protein lysate, entire scaffolds previously frozen at -80°C were incubated in 1X RIPA lysis buffer (Sigma, 20-188) supplemented with a protease and phosphatase inhibitor cocktail (Invitrogen, A32961) and sonicated at 20% amplitude for 20 pulses (1 second on, 1 second off). According to the manufacturer’s instructions, the protein concentration was quantified at a 1:200 dilution using a Bradford assay (Invitrogen, 23200).

For the analysis, 5μg of extracted protein was separated on a 4-12% Bolt Bis-Tris gel (Invitrogen, NW04125BOX) and transferred using an iBlot2 Transfer system (Invitrogen) to PVDF membranes (Invitrogen, IB242002) at 23V for 3 minutes. Membranes were blocked for 30 minutes at room temperature in 5% BSA or 5% non-fat dry milk (NFDM) (Jackson Immuno Research Labs, 001-000-161) in TBST (Boston BioProducts, IBB-181) and then incubated with primary antibodies diluted in 5% BSA or NFDM at 4°C overnight on a rocker. To remove excess antibodies, the membranes were washed twice in TBST followed by 3 10-minute washes at 4°C. The membranes were then incubated in secondary antibody solution for 1 hour in 5% BSA or NFDM at 4°C. Following three 10-minute washes at 4°C, the membranes were incubated for 5 minutes in the dark with SuperSignal™ West Pico Plus substrate ((Invitrogen, 34577) was used for actin and Tuj1 detection) or SuperSignal™ West Atto Ultimate Sensitivity substrate ((Invitrogen, A38555) used for detection of the rest of the proteins) to identify the proteins of interest. The membranes were serially imaged using a BioRad ChemiDoc MP. The images were analyzed using ImageJ Software.

Primary antibodies included: anti-DRP1 (ab184247 rabbit; 1:1000); anti-Fis1 (ab156865 rabbit; 1:1000); anti-MAP2 (ab5392 rabbit; 1:10000); anti-TOMM20 (ab186735 rabbit; 1:1000); anti-TOMM20 (ab56783 mouse; 1:1000); anti-beta-tubulin III (ab18207 rabbit; 1:1000); anti-beta-tubulin III (ab78078 mouse; 1:1000); anti-VDAC1 (ab154856 rabbit; 1:1000); anti-mitochondrial (ab92824 mouse; 1:1000); anti-actin HRP conjugated (ab20272 mouse; 1:1000); purchased from Abcam, and anti-DRP1 (phosphor S616) (3455S rabbit; 1:1000) purchased from Cell Signal Technologies.

Secondary antibodies included: HRP – conjugated (anti-rabbit IgG 7074P2; anti-mouse IgG 7076S) purchased from Cell Signal Technologies.

#### Protein Array

For the production level of 30 proteins associated with the neuronal state in health and disease, we used the Human Neuro Discovery Array (AAH-NEU-2-8, Ray Biotech) following the manufacturer’s protocol. In brief, 1 ml of conditioned media (pooled from 3 independent samples) was used per one membrane, with 3-4 total samples per condition (Sham and CCI, P110 untreated/treated), each from one independent experiment. The membranes were imaged using a BioRad ChemiDoc MP. The images were analyzed using ImageJ Software. The presented data is the ratio of CCI (contusion injury) over Sham (untreated).

#### Mitochondria isolation and treatment of 3D in vitro brain cultures

Mitochondria isolation and transfer experiments were conducted following established protocols (14). In brief, the media from 3D *in vitro* cultures (neurons, microglia, or astrocytes – depending on the experimental set-up) was collected into the 1.5 ml Eppendorf tubes and centrifuged for 10 minutes at 1,000 x g to pellet the nuclei. The post-nuclear supernatant was further centrifuged at 13,000 x g for 25 minutes to collect a mitochondria-enriched fraction. After mitochondria pellets were generated, the mitochondria were reconstituted in 100 µl of neuronal medium and transferred to 900 µl of media in naïve scaffolds seeded with neurons, microglia, or astrocytes (depending on the experimental set-up) for 24 hours of incubation. The treatment groups received a fresh dose of the P110 peptide at each transfer step. The residual mitochondria-free media after centrifugation was stored at -20°C for LDH and Glutamate release analysis. Isolated mitochondria were characterized for mitochondria-specific markers expression: mitochondria, TOMM20, and VDAC1 (Supplementary Fig. 14 a-b).

### RNA Isolation Protocol

Scaffolds were collected and stored at -80°C. Upon isolation, samples were thawed on ice and incubated with 600µL of Lysis Buffer from RNeasy Kit (Qiagen, Catalog No. 74106). During incubation, samples were cut up using small scissors and continued to sit in a lysis buffer for approximately 20 minutes. The lysed samples were placed onto QIAshredder Mini Spin (Qiagen) columns and spun at 10,000rpm for 2 minutes. An equal amount of 70% ethanol was added to the supernatant and mixed thoroughly. The ethanol-containing mixture was spun in two rounds through the RNeasy Mini Spin column (Qiagen) at 10,000rpm for 1 minute each; the flow through was discarded. In a new collection tube, 400µL of RNA wash from SurePrep RNA/DNA/Protein Purification Kit (Fisher Scientific, Catalog No. BP2802-50) was added to the RNeasy column and spun at the same setting for 1 minute; the flow through was discarded. After the first RNA Wash, 30µL of DNase and RDD buffer (1:7 ratio) containing solution from the RNase-Free DNase Set (Qiagen, Catalog No. 79254) was incubated on the columns at room temperature for 15 minutes before a second and third round of RNA was performed as well as a final spin to ensure the column was dry. The column was placed into a 1.5mL microcentrifuge tube and 22uL of RNA Elution Solution (Fisher Scientific) was added to the center of the column and spun at 12,000rpm for 1 minute. Samples were then read on the NanoDrop 2000 spectrophotometer and stored at -80°C before sequencing.

### Messenger RNA (mRNA) sequencing and bioinformatic analysis

Approximately 1μg of total RNA for each sample underwent RNA quality control analyses before library preparation and bulk mRNA sequencing as previously described (54). RNA integrity numbers (RIN) were determined with a Bioanalyzer 2100 (Agilent Technologies, Santa Clara, CA, USA) which indicated minimal RNA degradation. Illumina Novaseq sequencers generated paired-end reads (150bp) at a read depth of 20 million reads (54).

Raw sequencing reads (.fastq files) were uploaded to the Tufts University Galaxy Server (55) for analysis, first utilizing FastQC (https://www.bioinformatics.babraham.ac.uk/) for quality control before read alignment with RNA STAR (56) (reference genome GRCh38.p13, https://www.ncbi.nlm.nih.gov/) and quantification using featureCounts (57). Differentially expressed gene lists were created using the DESeq2 R package (58) by comparing injury and Sham conditions, with or without P110 treatment. Gene set enrichment analyses (GSEA) were performed using the desktop software (https://www.gsea-msigdb.org/gsea/index.jsp) (59, 60) using MitoCarta3.0 (61) and custom curated mitochondrial pathway (62) databases. Heatmaps were generated using the pheatmap R package. Gene module co-expression analyses were performed with the CEMiTool R package (63, 64), GSEA using Reactome pathway database (https://reactome.org/), and signed interaction network analysis utilizing high confidence STRING protein-protein interactions (STRING score > 700).

### Metabolic imaging and Data analysis

Images were acquired using a Leica TCS SP8 confocal microscope equipped with a tunable (680 to 1300 nm) femtosecond laser (InSight Deep See; Spectra-Physics; Mountain View, California) and a water-immersion 40× objective (NA 1.1). Two-photon excited fluorescence (TPEF) images (1024 × 1024 pixels, 290.6 × 290.6 μm) were acquired at 755 nm excitation and two emission ranges at 460 ± 25 nm and 525

± 25 nm. Location co-registered NAD(P)H lifetime data were then acquired at the same excitation and emission wavelengths with a 1-minute integration time (512 × 512 pixels) using a Picoquant Picoharp 300 time-correlated single photon counter and SymPhoTime analysis software. A three-level Otsu’s threshold and a combination of two-dimensional (2D) discrete Fourier transform and power spectral density (PSD) methods were applied on binarized images to segment out the mitochondria regions (65). 2D pixel-wise redox ratio maps were generated based on the fluorescence intensities from two different emission channels with the expression of ⑤②, with the 525 nm channel for FAD fluorescence, and the 460 nm channel for NAD(P)H fluorescence.

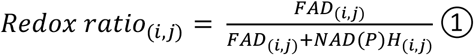

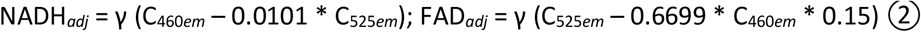

Lifetime data were saved into 261-time bins with equal intervals (12.5ns in length) and processed using a phasor approach as previously described (66). Briefly, this approach does not require prior knowledge of the fluorophores within the samples. It has a high sensitivity to environmental perturbations and can provide complementary information on the nature of fluorophores to the intensity data. In this study, we introduced a phasor-guided redox ratio distribution analytical approach, where we considered the redox ratio from NAD(P)H-containing pixels by mapping the location of specific regions of the phasor distribution data to the original lifetime images and then to the co-registered pixels in the NAD(P)H and FAD intensity images. After Fourier transformation, the pixel-wise lifetime information was converted to a set of g and s cos and sin components, respectively. The redder hues in the phasor map represent more pixels with similar frequency components, while the bluer hues have fewer ones (Fig. 5h and Supplementary Figures 32-33). A linear fit was applied to all the g and s phasor values corresponding to the lifetime spectra from all pixels of a field. The two points where the line intersected the universal semicircle represented the short and long NAD(P)H lifetimes; the bound NAD(P)H fraction was estimated based on the projected location of each phasor point on the fitted line, according to the equation ③ as shown in Supplemental Fig. 21 (66).

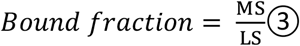

The pixel-wise redox ratio values were extracted by mapping back the NAD(P)H phasor locations to the original 2D images and referring to the co-registered NADH and FAD intensity images. All redox values from each sample were sorted into 10 bins ranging from 0 to 1, with 0.1 intervals. A metric AUC ratio, calculated according to equation ③, was used to quantify the redox distribution differences among different treatment groups (CCI vs Sham, P110 CCI vs P110 Sham).

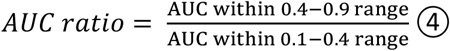

#### Region selection

Autofluorescence lifetime spectra from all the samples of this study were mapped into four main regions of the phasor space (Supplementary Fig. 32). Data within the heavily overlapped Q1 and Q2 regions were present in all fields and accounted for most of the pixels (>95%, Supplementary Fig. 34 G), while lifetime spectra that were mapped within the Q3 and Q4 regions could only be occasionally detected with varying prevalence. Redox distributions from the Q1, Q2, and overlapped regions were almost identical (Supplementary Fig. 34 F), indicating they may be corresponding to NADH and NADPH (67). The redox ratio distributions of the pixels mapped to the Q3 and Q4 phasor spaces were more right-shifted, indicating distinct origin, possibly from retinol and retinoic acids (68). Therefore, to avoid artifacts introduced by other fluorophores besides NAD(P)H, redox analysis in this study, focused on pixels with lifetime spectra that were mapped within the Q1, Q2, and their overlapped regions.

All the intensity images were power and detector gain normalized before analysis according to equation *⑤*. Incident power was measured with a power meter before every experiment session before the objective.

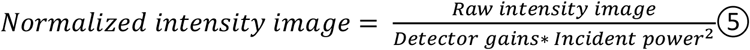

Two cell lines, HMC3 and iPSCs-derived microglia, each with two independent rounds, three dependent transfers, and four treatment conditions were imaged. For each treatment, 2-4 samples, each with 3-4 locations were acquired. In total, for each cell line, results were from at least 5 samples from each treatment condition (*N*_*Sham*_=5-6, *N*_*P*110,*Sham*_=5-6, *N*_*P*110,*CCI*_=7-8, *N*_*P*110,*Sham*_=7-8). All the CCI (contusion injury) data were normalized to corresponding shams.

#### Statistical analysis

Statistical analysis was performed between and within experimental groups using GraphPad Prism 9 software. Two-tailed t-tests were used to compare values within an experimental group and between two experimental groups. One-or two-way ANOVA analysis of variance was used to compare multiple groups. Tukey’s post hoc tests were used to assess computed significant differences between experimental and control groups. Any p-value less than 0.05 was considered statistically significant. Each experiment was repeated at least three times; technical replicates (2-6, depending on the experiment) were used for every assay. Quantified data graphically presented the mean and standard error of the mean (SEM) of each group.

