## Supplementary document for "Mitochondria dysregulation contributes to secondary neurodegeneration progression post-contusion injury in human 3D in vitro triculture brain tissue model"

### Supplementary Results

Initially, cultures were incubated in a Neurobasal medium supplemented with 1% Anti-Anti, 1% Glutamax, and 2% B27 supplement, and cellular viability was checked weekly with calcein (live) staining. Two weeks after seeding, astrocytes in monocultures and co-cultures started to die rapidly (Figure Supplementary 1a, red arrows indicate dying cells). To prevent astrocytic death, 1% astrocytes growth factors (ScienCell Research Laboratories) were introduced in the previously described neuronal medium, and viability was checked weekly. Astrocytes in monocultures showed rapid growth and proliferation. In co-cultures and tricultures, we observed high neuronal network density formation without the obvious presence of dead cells (Supplementary Fig. 1 b). For all consequent experiments in this manuscript, cultures were incubated in the Neurobasal medium supplemented with astrocytes growth factors in addition to B27, Glutamax, and anti-anti.

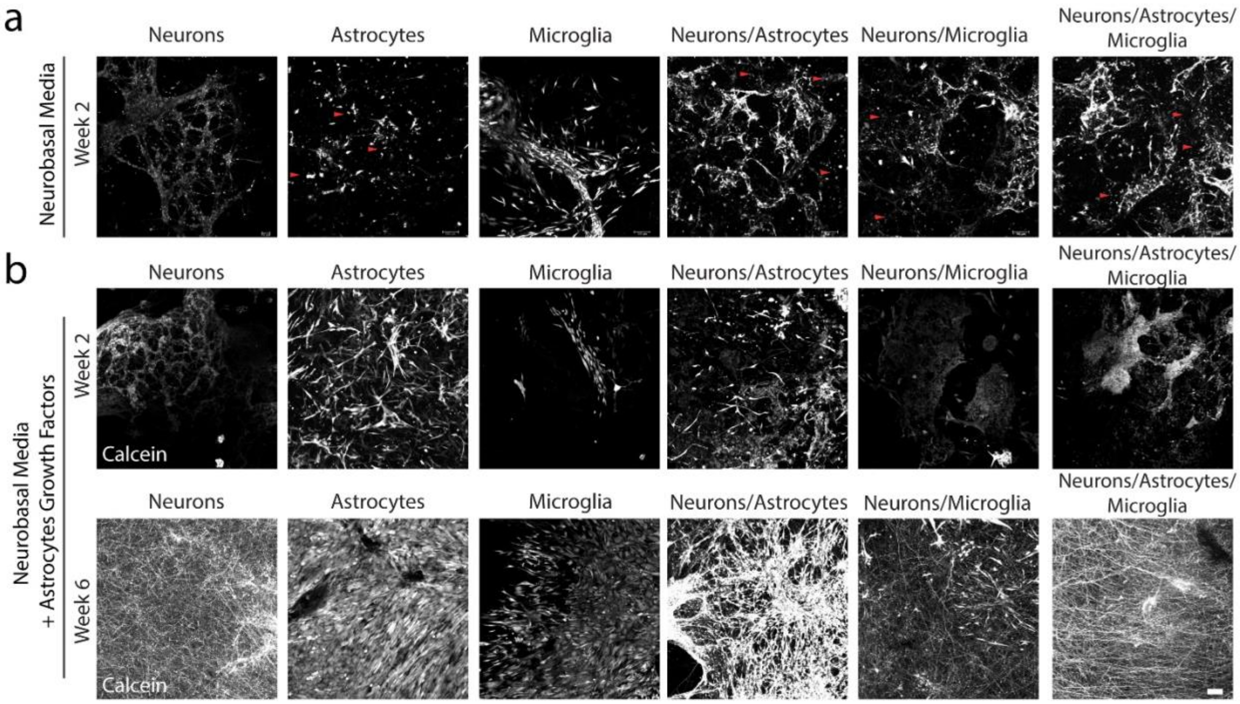


**Supplementary Fig. 1. Astrocyte growth factors were crucial in maintaining the viability of 3D tissues over time. a,** Representative images of calcein-stained live samples at 2 after seeding. Astrocytes and co-cultures with astrocytes demonstrated a high death rate when incubated in a neurobasal medium without astrocytes growth factors 2 weeks after seeding. Red arrows point to dead cells. **b,** Addition of astrocytes growth factors into the Neurobasal medium supported all cultures' growth and development at 2 and 6 weeks after seeding. Scale bar: 100μm.

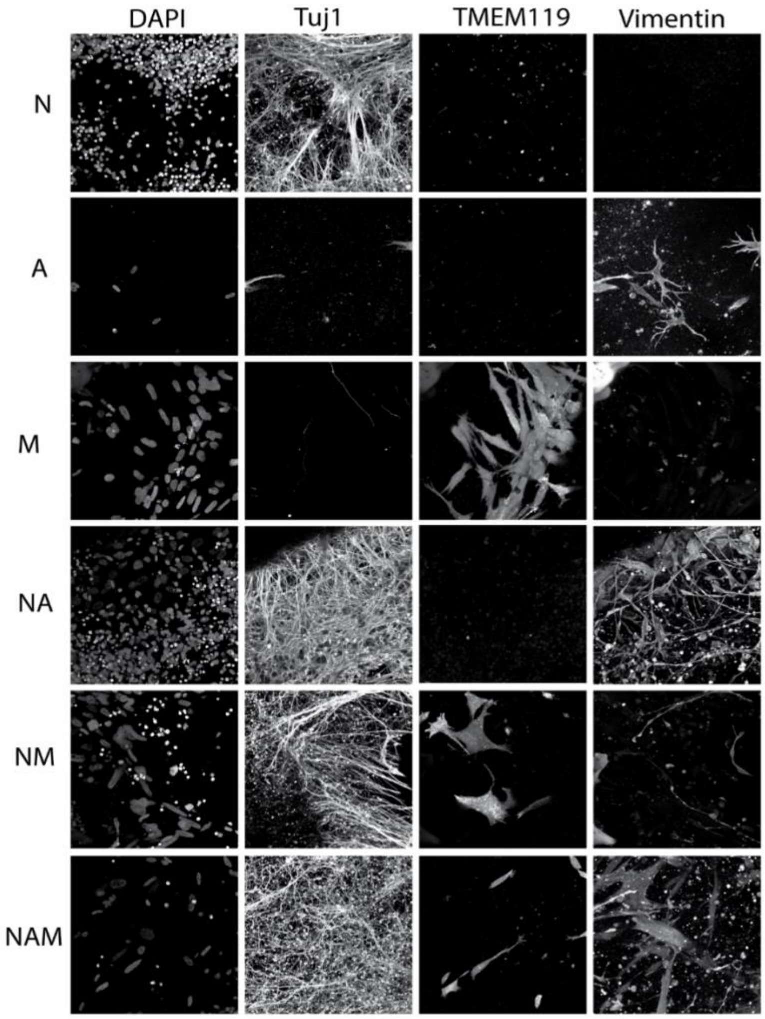


**Supplementary Fig. 2 Human in vitro mono-, di- and triculture model composed of neurons, microglia, and astrocytes.** Representative images of neuronal (Tuj1), microglia (TMEM119), and astrocytes (vimentin) specific cytoplasmic markers in mono-, di- and tricultures. Scale bar: 50µm.

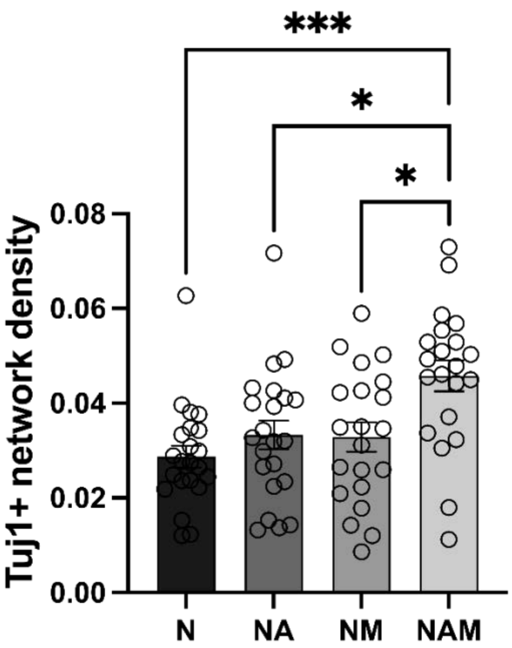


**Supplementary Fig. 3 Human in vitro mono-, di- and triculture model composed of neurons, microglia, and astrocytes.** Quantification of Tuj1 positive network from Supplementary Fig. 2. Datta presented as mean ±SEM of three independent experiments with n=4 scaffolds per condition. *, *** indicates significant differences ($p<0.05$, 0.001 respectively; two-way ANOVA (analysis of variance) (with Tukey's post-hoc tets) between control and experimental groups).

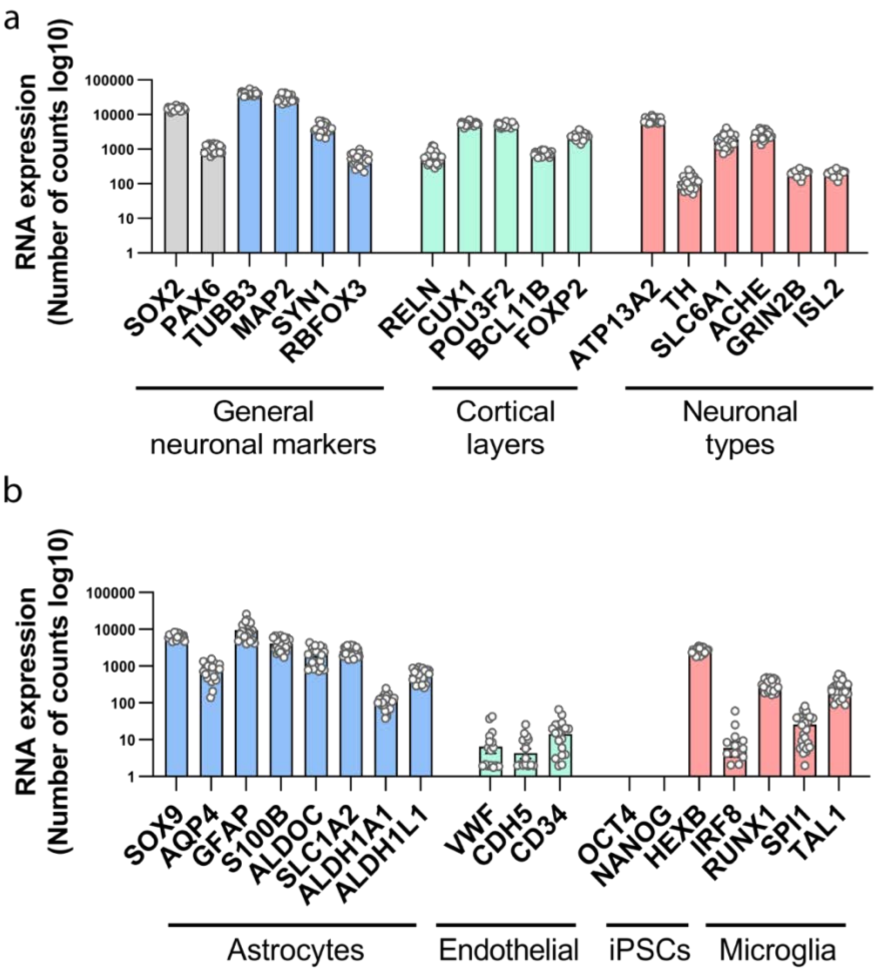


**Supplementary Fig. 4 Human 3D in vitro triculture model characterization.** mRNA expression (count number via bulk mRNA sequencing) of **a,** neuron-specific makers (SOX2, PAX6 - progenitor state; TUBB3 – pan-marker; MAP2 – dendritic; SYN1 – synaptogenesis; NeuN – transcriptional factor of neuronal maturation): cortical layers specific (RELN – Layer I; CUX1 – Layer II-IV; POU3F2 – Layer V-VI; BCL11B – Layer V, striatal neurons; FOXP2 – Layer V-VI); and neuronal subtypes specific (ATP12A2, TH – Dopaminergic neurons; SLC6A1 – GABAnergic; ACHE – Cholinergic; GRIN2B – Glutaminergic; ISL2 – motor neurons)); **b,** astrocyte-specific (SOX9, AQP4, GFAP, S100B, ALDOC, SLC1A2, ALDH1A1, ALDH1L1); endothelial specific (VWF, CDH5, CD34); iPSC (OCT4, NANOG); microglia specific (HEXB, IRF8, RUNX1, SPI1, TAL1) markers. Each marker shows n=12 samples isolated from tricultures composed of induced neural stem cells derived neurons, primary astrocytes, and iPSC-derived microglia from two healthy donors, YZ1 and ND418664*F.

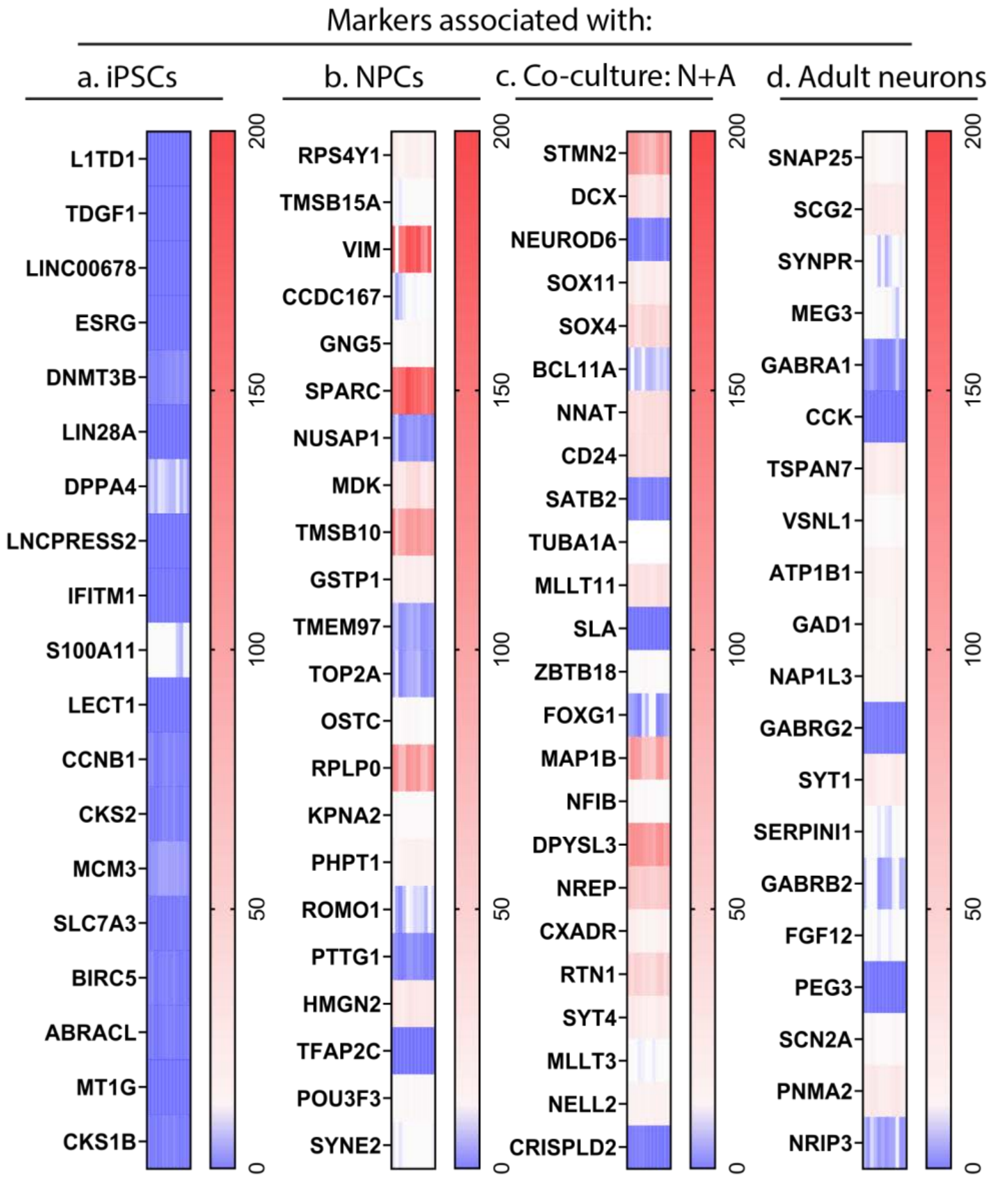


**Supplementary Fig. 5 Neuronal Maturation markers are highly expressed relative to hiPSC markers in the human 3D in vitro triculture model.** mRNA expression (count number via bulk mRNA sequencing) of **a,** induced pluripotent stem cell (iPSCs); **b,** neural progenitor cells (NPCs); **c,** iPSCs-derived neuronal + astrocytes co-culture; and **d,** human adult neuronal markers. Each marker shows n=12 samples isolated from tricultures composed of induced neural stem cells derived neurons, primary astrocytes, and iPSC-derived microglia from two healthy donors, YZ1 and ND418664*F.

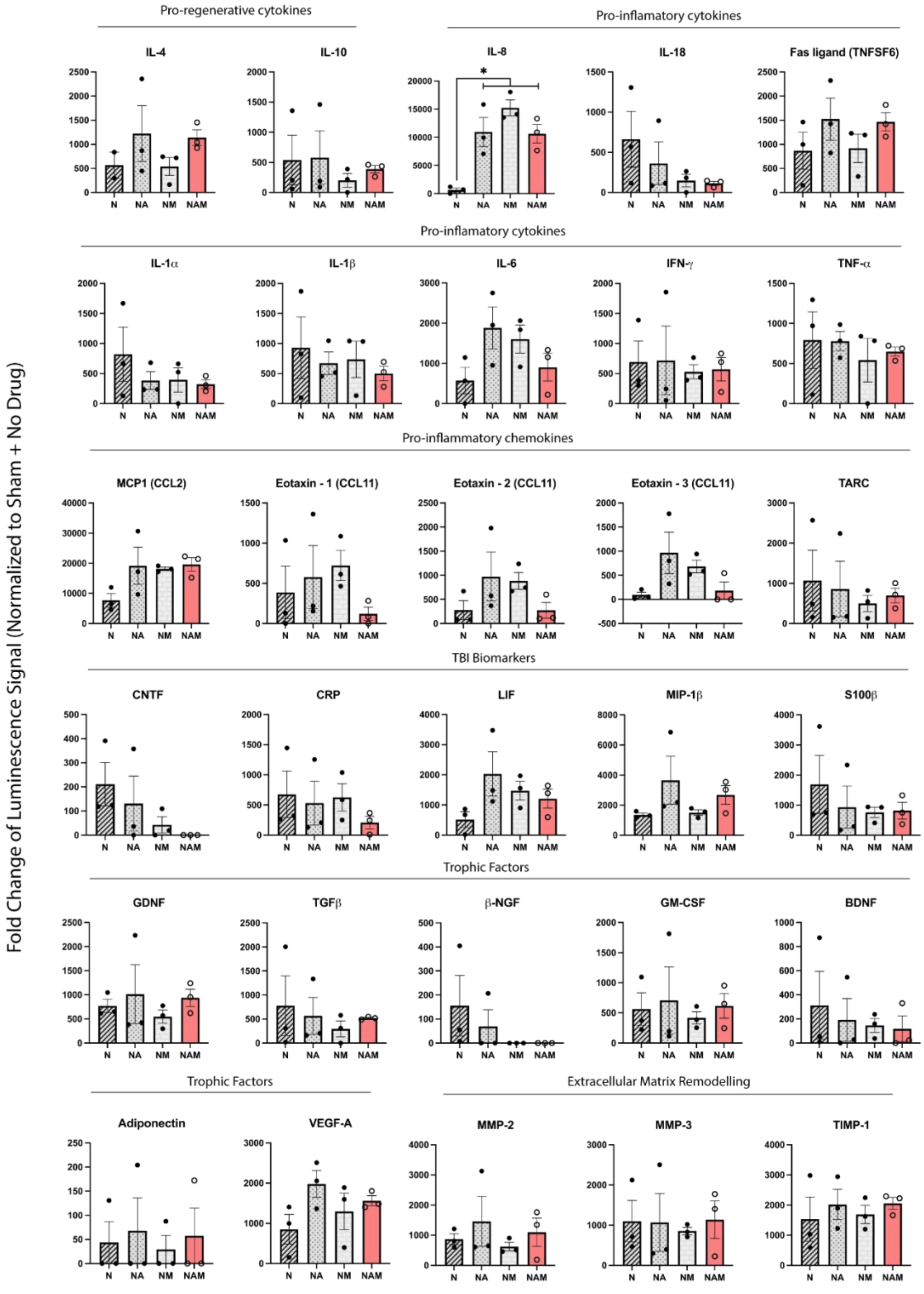
Pro-regenerative cytokines
Pro-inflamatory cytokines
IL-4
IL-10
IL-8
IL-18
Fas ligand (TNFSF6)
Pro-inflamatory cytokines
IL-1α
IL-1β
IL-6
IFN-γ
TNF-α
Pro-inflammatory chemokines
MCP1 (CCL2)
Eotaxin - 1 (CCL11)
Eotaxin - 2 (CCL11)
Eotaxin - 3 (CCL11)
TARC
TBI Biomarkers
CNTF
CRP
LIF
MIP-1β
S100β
Trophic Factors
GDNF
TGFβ
β-NGF
GM-CSF
BDNF
Trophic Factors
Extracellular Matrix Remodelling
Adiponectin
VEGF-A
MMP-2
MMP-3
TIMP-1
N
NA
NM
NAM
Fold Change of Luminescence Signal (Normalized to Sham + No Drug)

**Supplementary Fig.6.** Neuronal co-culture with astrocytes and/or microglia shows immune homeostasis 6 weeks after seeding. The production level of Pro-regenerative and pro-inflammatory cytokines and chemokines, TBI biomarkers, Trophic factors, and ECM remodeling proteins. Data presented as mean ± SEM of three independent experiments with each data point is the average for n=4 scaffolds per condition. *, **, ***, **** indicate significant differences (p<0.05, 0.01, 0.001, 0.0001 respectively; two-way ANOVA (analysis of variance) between control and experimental groups).

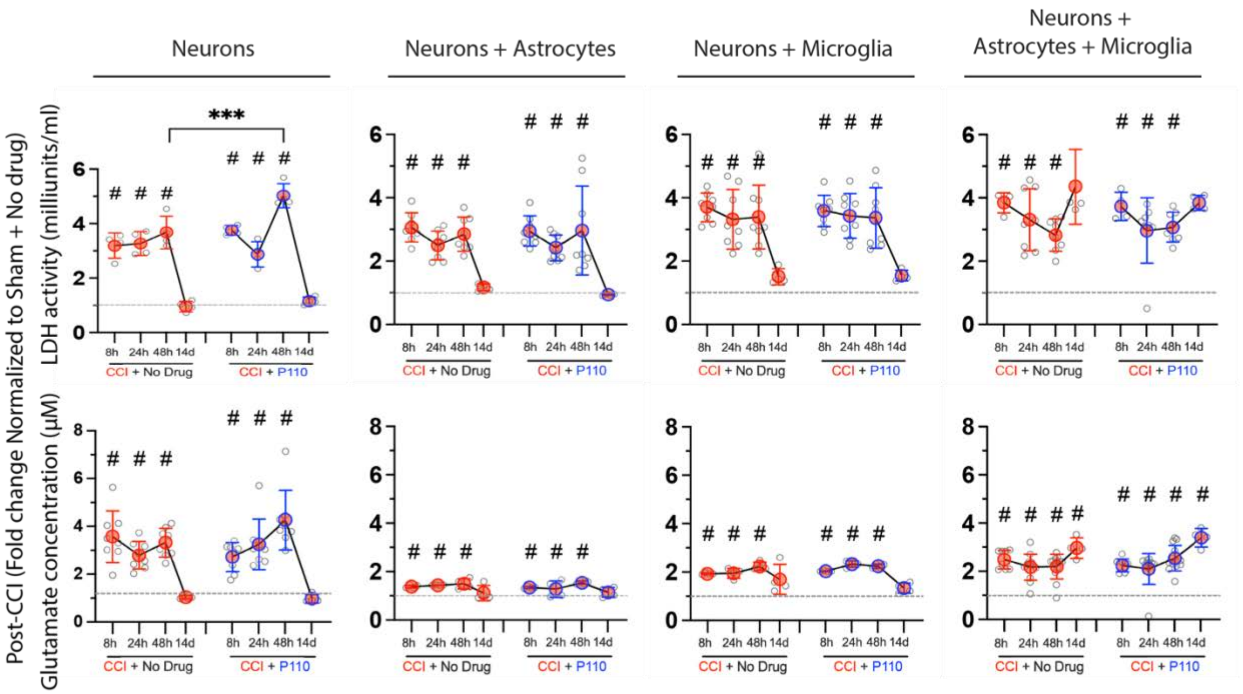


**Supplementary Fig. 7. Biochemical response to TBI in human 3D in vitro model. Top,** LDH and **Bottom,** Glutamate release after controlled cortical impact injury in human 3D cultures of neurons, microglia, and astrocytes in mono-, di- and tricultures at 4-time points: 8, 24, 48 hours, and 14 days. Data presented as mean ±SEM of three independent experiments with n=2-4 scaffolds per condition. *, **, ***, **** indicate significant differences (p<0.05, 0.01, 0.001, 0.0001 respectively; two-way ANOVA (analysis of variance) (with Tukey's post-hoc test) between control and experimental groups).

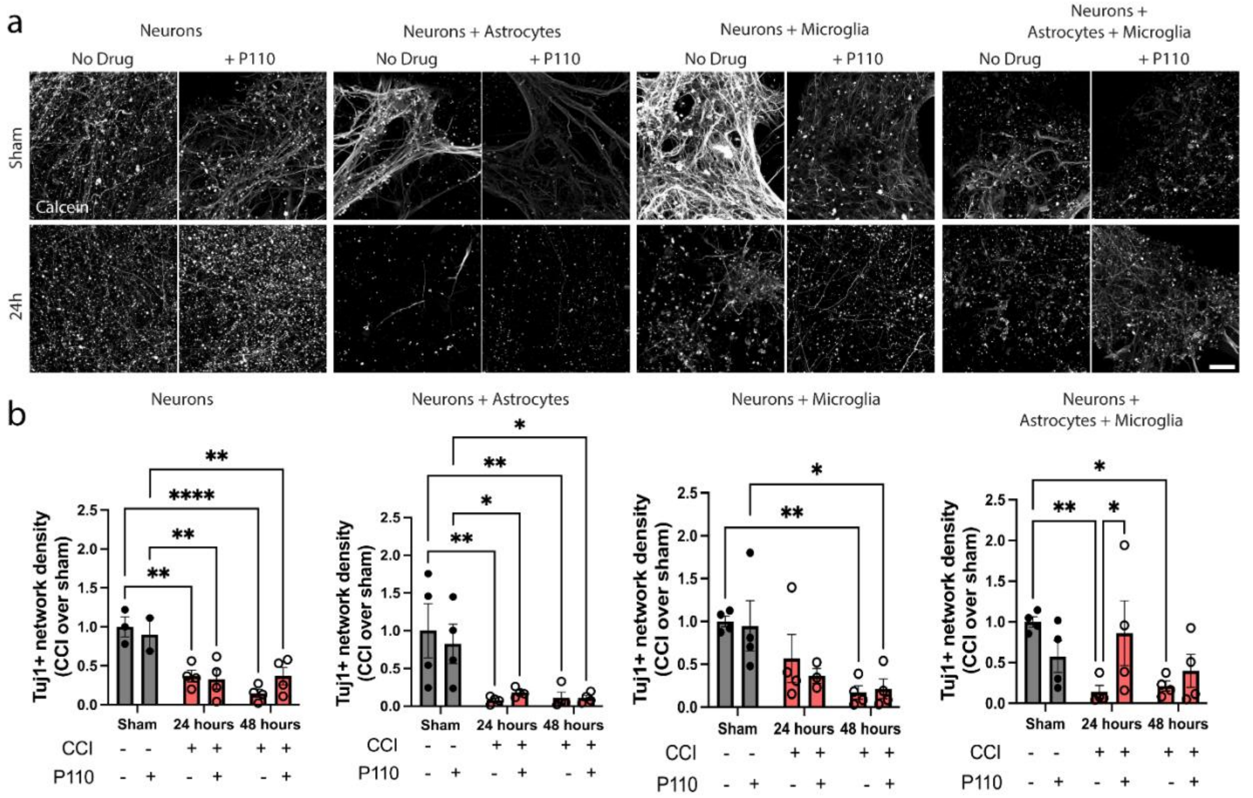


**Supplementary Fig. 8 Neuronal network degeneration in the injury site 24 hours after controlled cortical impact injury. a,** Representative images of Tuj1 neuronal staining at 24 hours after CCI in N, NA, NM, and NAM groups. **b,** Quantification of Tuj1 neuronal network density in control and P110 treated groups of N, NA, NM, and NAM at two different time points (24 and 48 hours). Data presented as mean ±SEM of three independent experiments with n=2-4 scaffolds per condition. *, **, ***, **** indicate significant differences (p<0.05, 0.01, 0.001, 0.0001 respectively; two-way ANOVA (analysis of variance) between control and experimental groups). Scale bar: 50 µm.

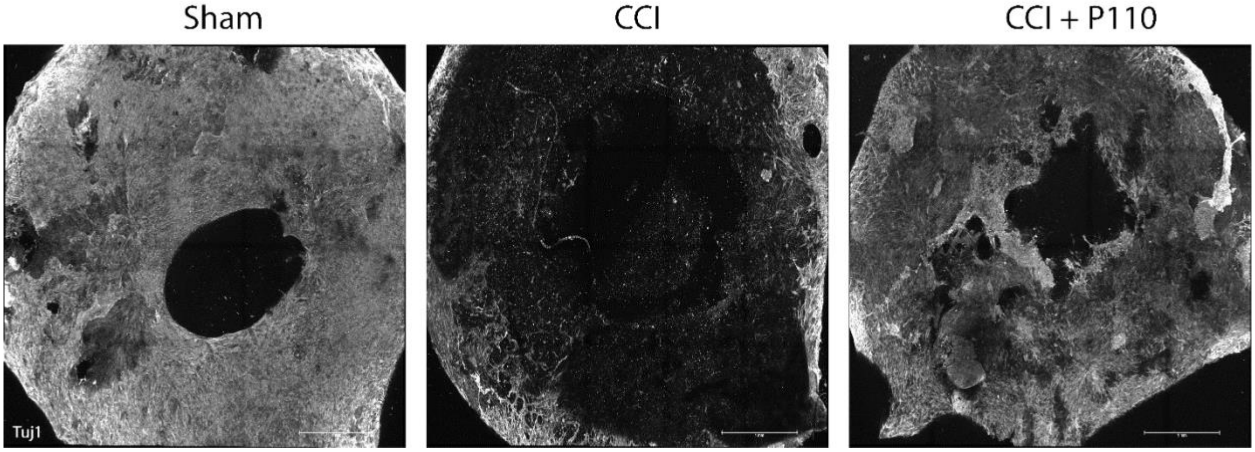


**Supplementary Fig. 9. Contusion injury induces mitochondria fission-dependent progressive neuronal network degradation in the human 3D triculture model.** Representative images of Tuj1 neuronal network staining at 24 hours after the injury in NAM groups with and without P110 treatment. Scale bar: 1mm.

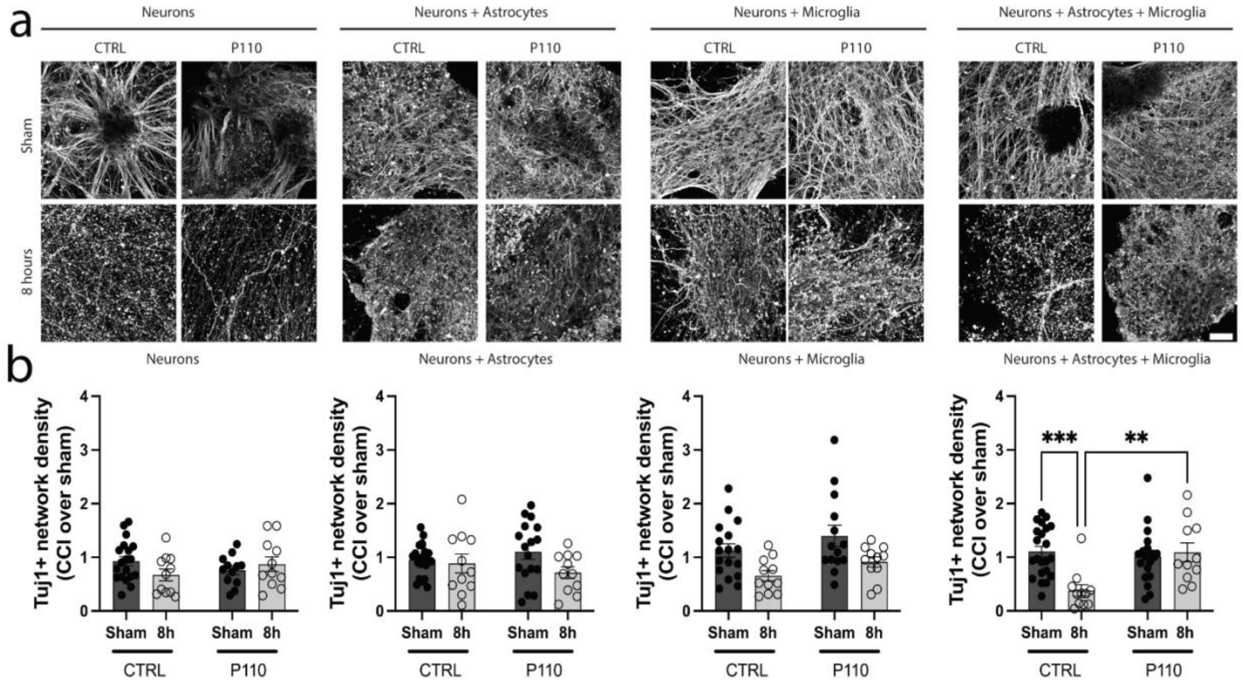


**Supplementary Fig. 10. Crosstalk between astrocytes and microglia had a detrimental effect on neuronal survival after brain injury and was reversible with mitochondria fission inhibitor P110. a,** Representative images of Tuj1 neuronal staining at 8 hours after contusion in N, NA, NM, and NAM groups in the proximal area. **b,** Quantification of Tuj1 neuronal network density in control and P110 treated groups. Data presented as mean ±SEM of three independent experiments with n=2-4 scaffolds per condition. *, **, ***, **** indicate significant differences ($p<0.05$, 0.01, 0.001, 0.0001 respectively; two-way ANOVA (analysis of variance) between control and experimental groups).

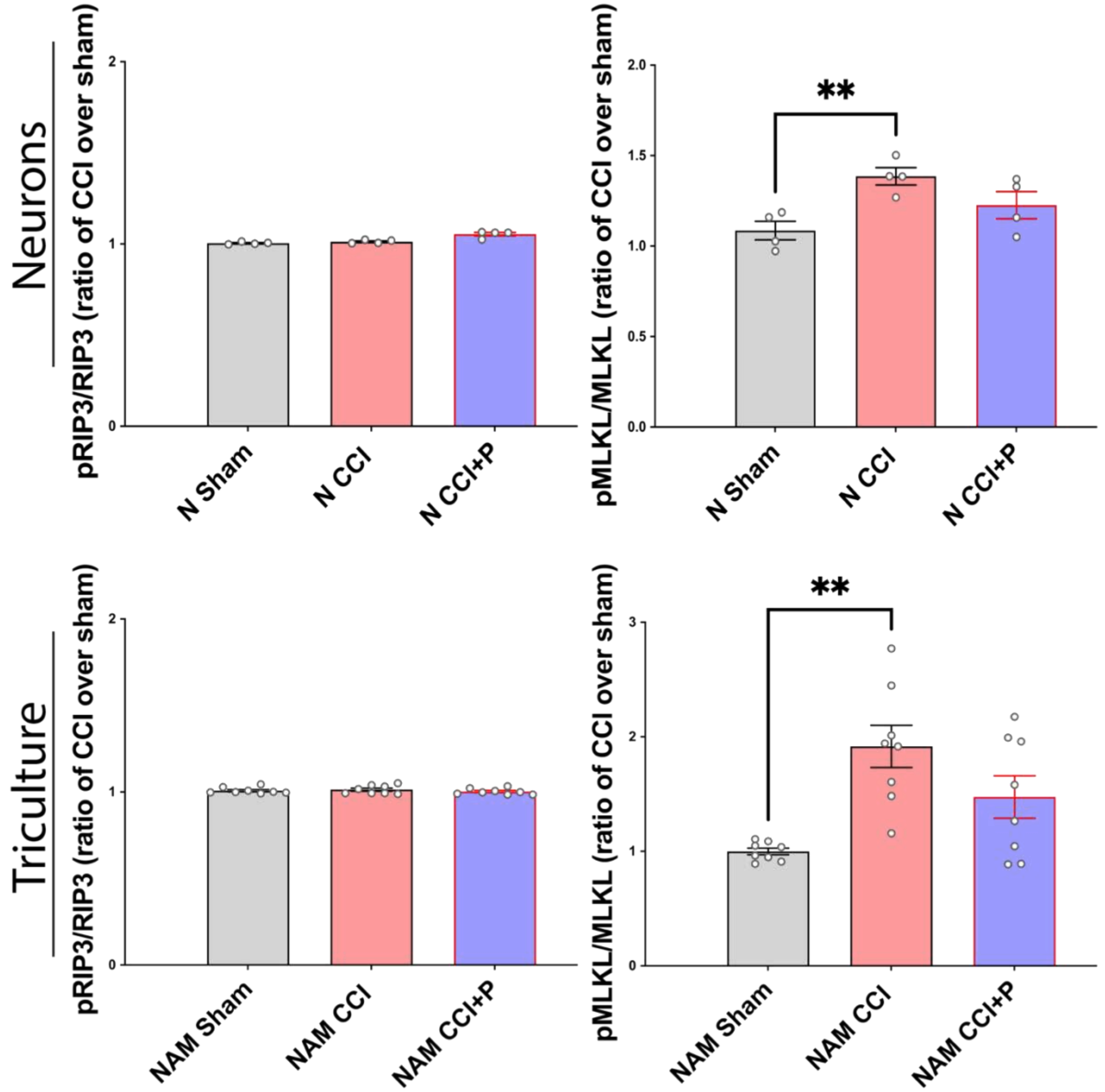


**Supplementary Fig. 11 Contusion injury induces necroptotic death progression independent of mitochondria fission in neuronal monoculture and tricultures.** Densitometry analyses of phospho-RIP3 (p-RIP3) and p-MLKL protein amount 24 hours after contusion injury in control and P110 treated groups. Data presented as mean ±SEM of two independent experiments with n=2-4 scaffolds per condition. *** indicates a significant difference (p< 0.01; one-way ANOVA (with Tukey post-hoc test) between control and experimental groups).

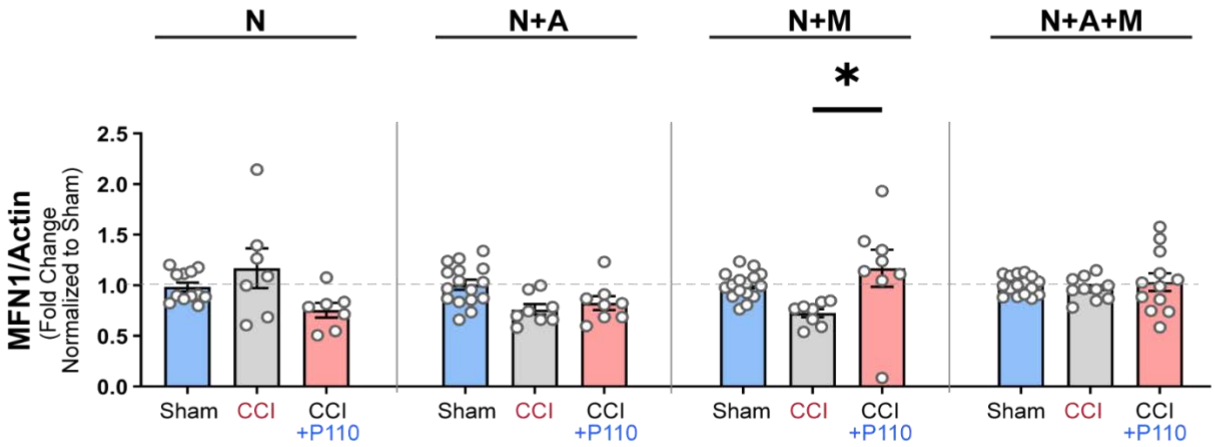


**Supplementary Fig. 12. Secondary neurodegeneration in tricultures was associated with increased mitochondria fission but not fusion.** Mitofusin 1, 2 protein amount quantification of Western Blots 24 hours after injury in control and P110 treated groups. Data presented as mean ±SEM of three independent experiments with n=2-4 scaffolds per condition. *, **, ***, **** indicates significant difference (p<0.05, 0.01, 0.001, 0.0001 respectively; one-way ANOVA (analysis of variance) between control and experimental groups).

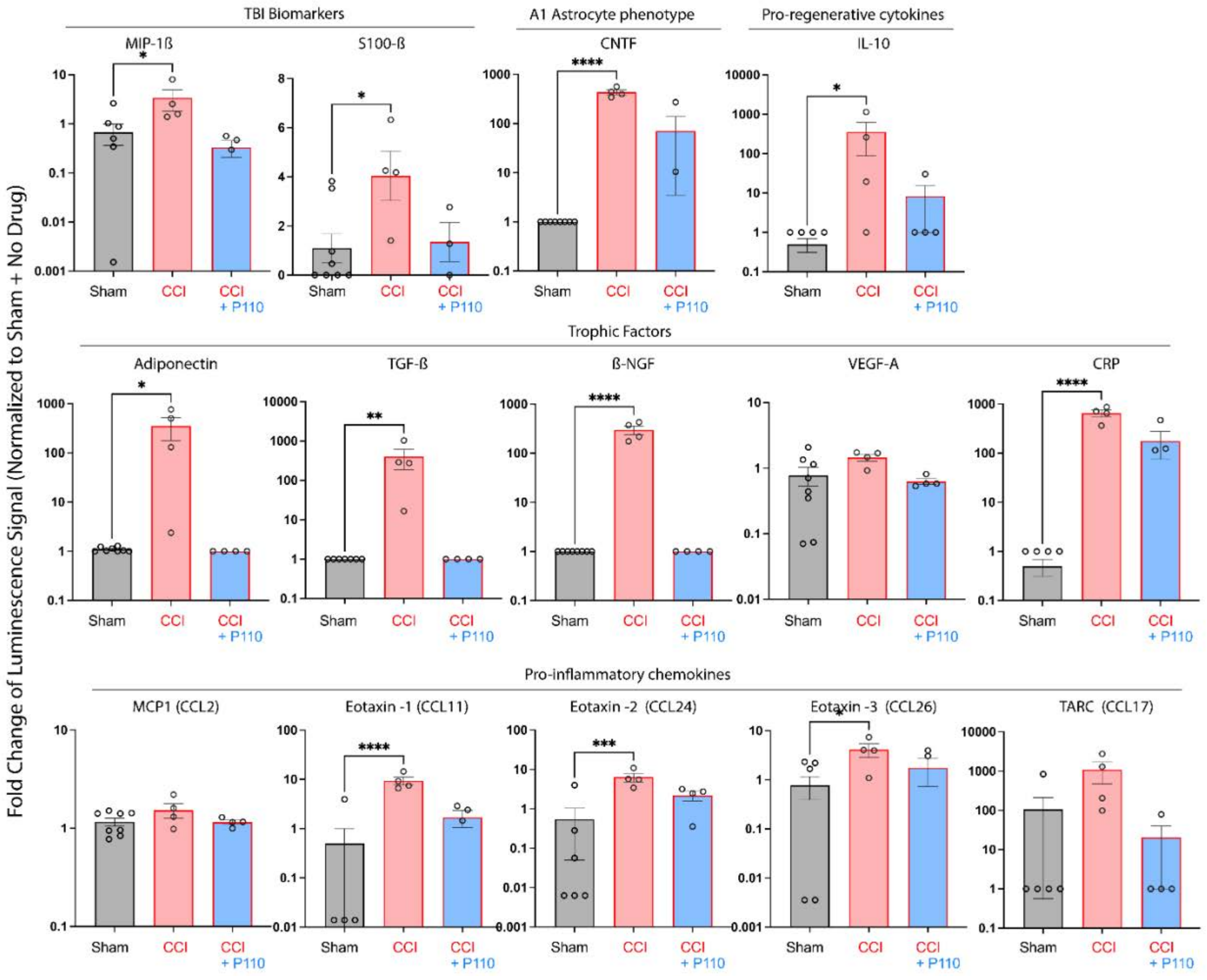

**Supplementary Fig. 13. Increased mitochondria fission was associated with neuroinflammation in tricultures 24 hours after the injury.** The production level of TBI biomarkers, A1 Astrocyte associated markers, Pro-regenerative cytokines, Trophic factors, and Pro-Inflammatory chemokines. Data presented as mean ± SEM of four independent experiments with each data point is the average for n=2-4 scaffolds per condition. *, **, ***, **** indicate significant differences ($p<0.05$, 0.01, 0.001, 0.0001 respectively; two-way ANOVA (analysis of variance) between control and experimental groups).

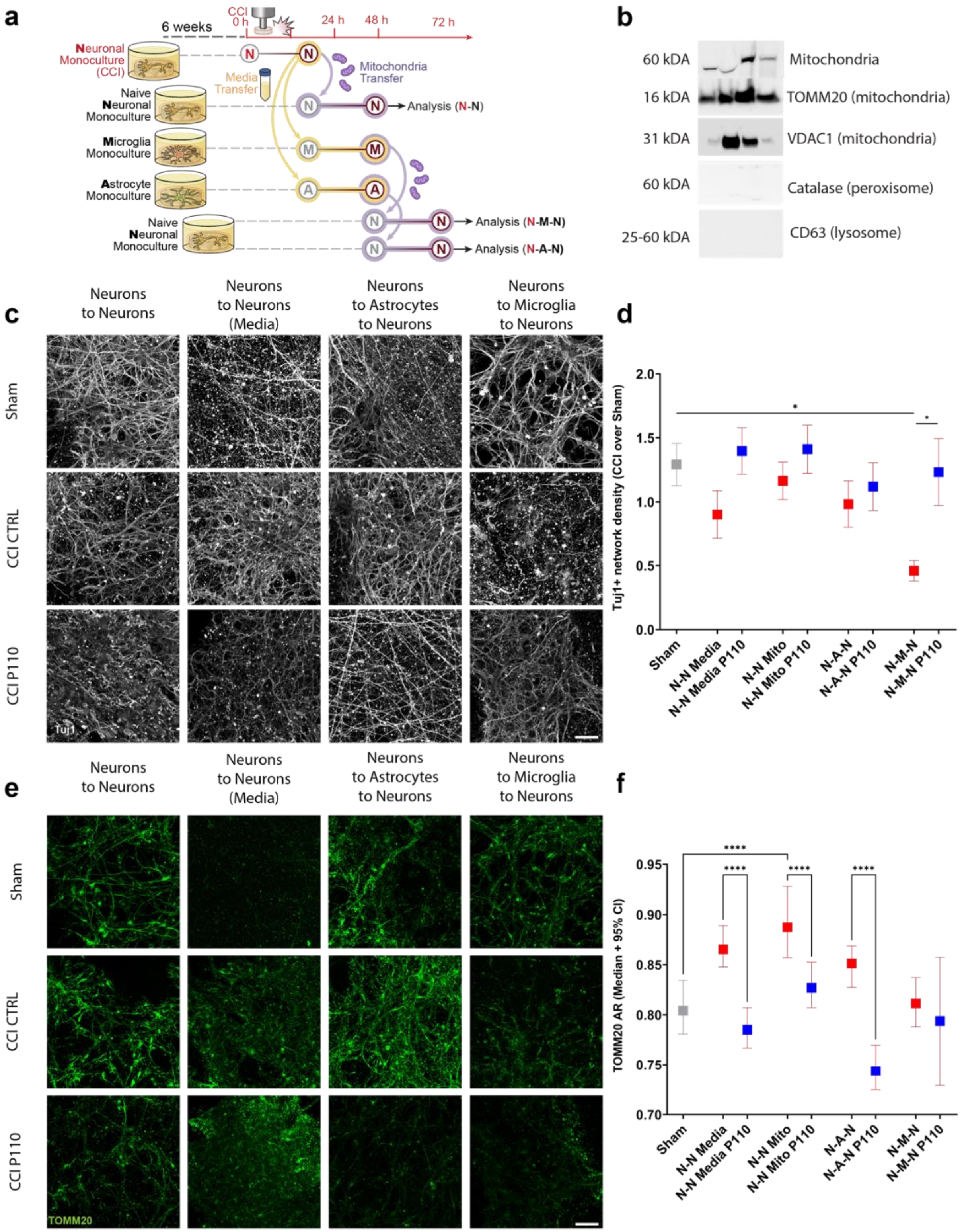


**Supplementary Fig. 14. Mitochondria fragmentation in microglia was crucial for secondary neurodegeneration and neuroinflammation progression post-CCI. a,** Schematic representation of transfer experiments from neurons to neurons, or astrocytes, or microglia. **b,** Western blot analysis of mitochondria-specific proteins expression: anti-mitochondria, TOMM20, and VDAC1and of peroxisomes (catalase) and lysosomes (CD63) to ensure the purity of mitochondria isolation. Representative images of **c,** Tuj1, and **e,** TOMM20 staining of naïve neurons treated for 24 hours with mitochondria isolated from microglia treated with conditioned media collected from 24h injured neurons with quantification of **d,** Tuj1 positive neuronal network density, and **f,** TOMM20 positive mitochondria aspect ratios. Data presented in **(d, f)** mean ±SEM of n=2-10 scaffolds per condition. *, **, ***, **** indicates significant difference (p<0.05, 0.01, 0.001, 0.0001 respectively; one-way or two-way ANOVA (analysis of variance) between experimental groups). Experiments were replicated at least three times. Scale bar: 50 µm.

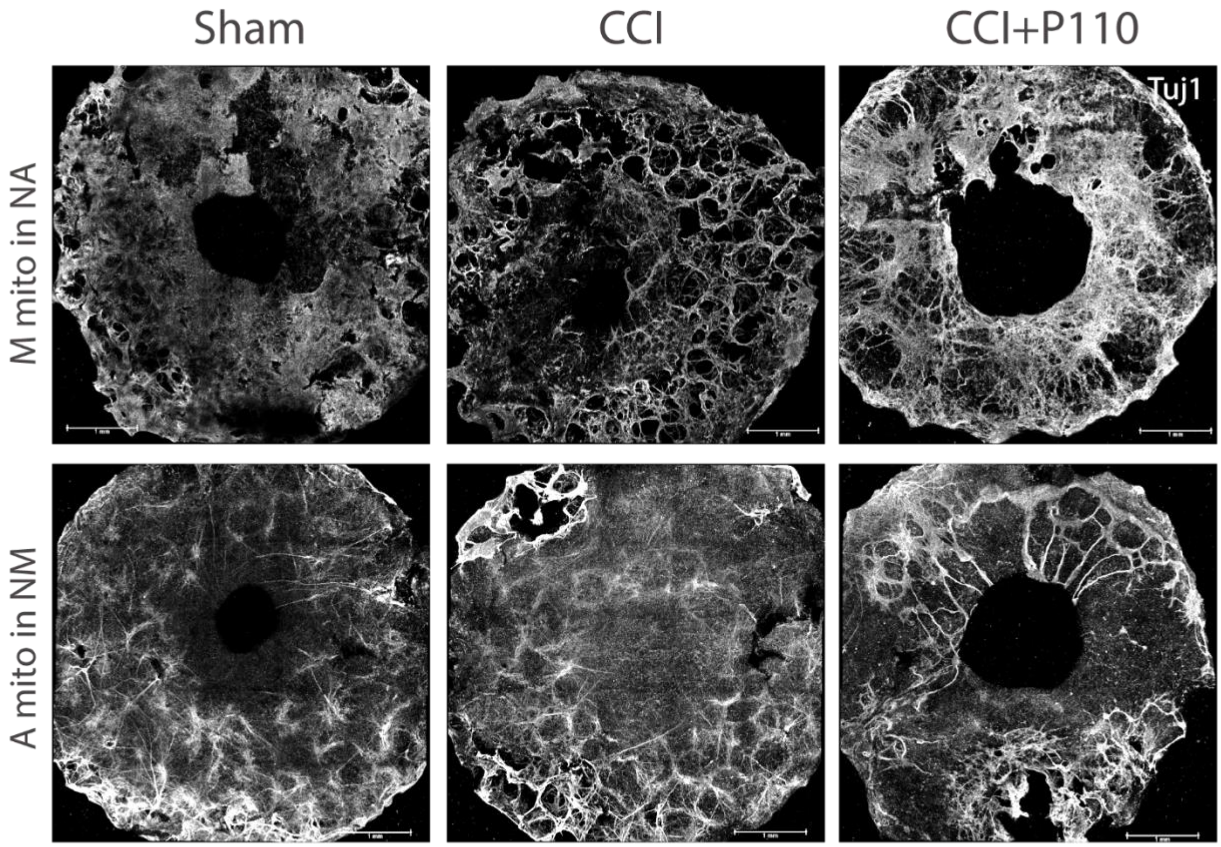


**Supplementary Fig. 15. Mitochondria released from microglia activated by conditioned media from injured neurons induced neuronal network degeneration in naïve co-culture of neurons with astrocytes in mitochondria fission dependent manner, but not by mitochondria released from astrocytes activated by conditioned media from injured neurons induced neuronal network degeneration in naïve co-culture of neurons with microglia.** Representative images of Tuj1 neuronal network staining 24 hours after treatment in NAM groups with and without P110 treatment. Scale bar: 1mm.

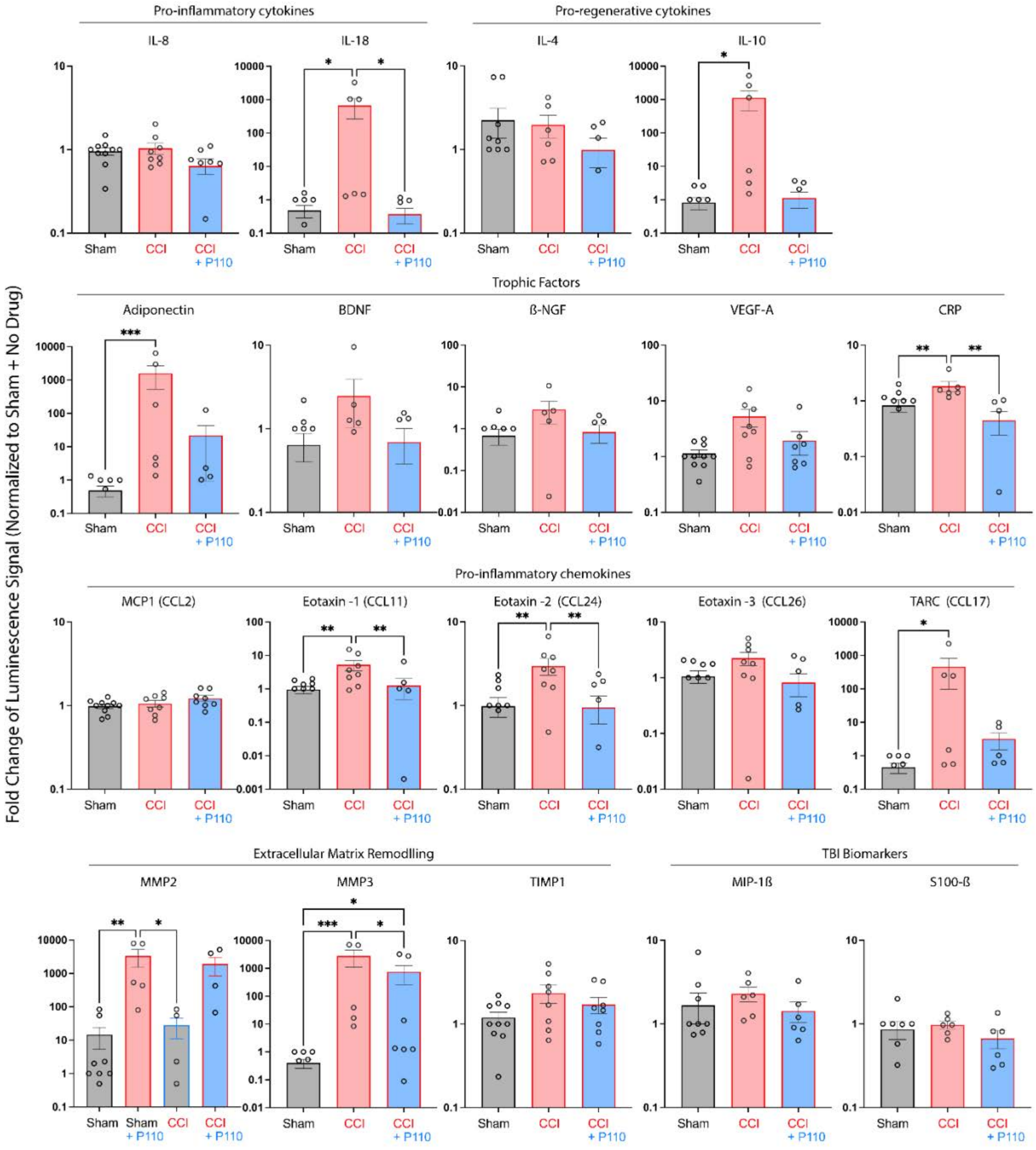


**Supplementary Fig. 16. Mitochondria dysregulation in microglia was crucial for secondary neurodegeneration and neuroinflammation progression post-CCI.** The production level of Pro-inflammatory and pro-regenerative cytokines, trophic factors, pro-Inflammatory chemokines, extracellular matrix remodeling proteins, and TBI biomarkers. Data presented as mean ± SEM of four independent experiments with each data point is the average for n=2-4 scaffolds per condition. *, **, ***, **** indicate significant differences (p<0.05, 0.01, 0.001, 0.0001 respectively; two-way ANOVA (analysis of variance) between control and experimental groups).

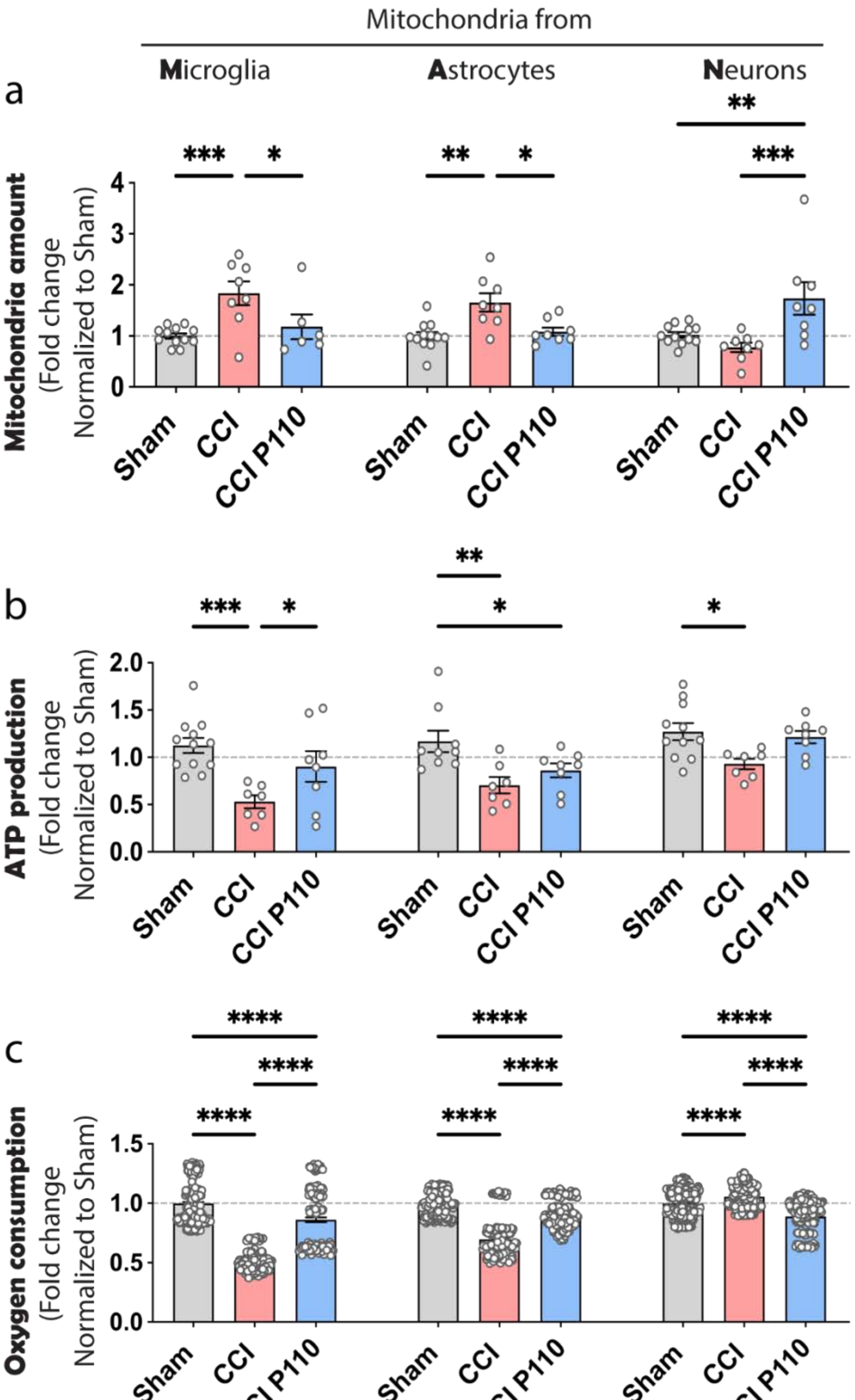


**Supplementary Fig. 17 Conditioned media from injured neurons induced the release of dysfunctional mitochondria from microglia, astrocytes, and neurons. a,** DNA quantification; **b,** ATP production; **c,** Oxygen consumption by the mitochondria released from microglia, astrocytes, and neurons activated by treatment with conditioned media from injured neurons for 24 hours. Data presented as mean ±SEM of n=4-8 scaffolds per condition from 2 independent experiments. *, **, ***, **** indicates significant difference (p<0.05, 0.01, 0.001, 0.0001 respectively; one-way or two-way ANOVA (with Tukey's post-hoc test) between experimental groups).

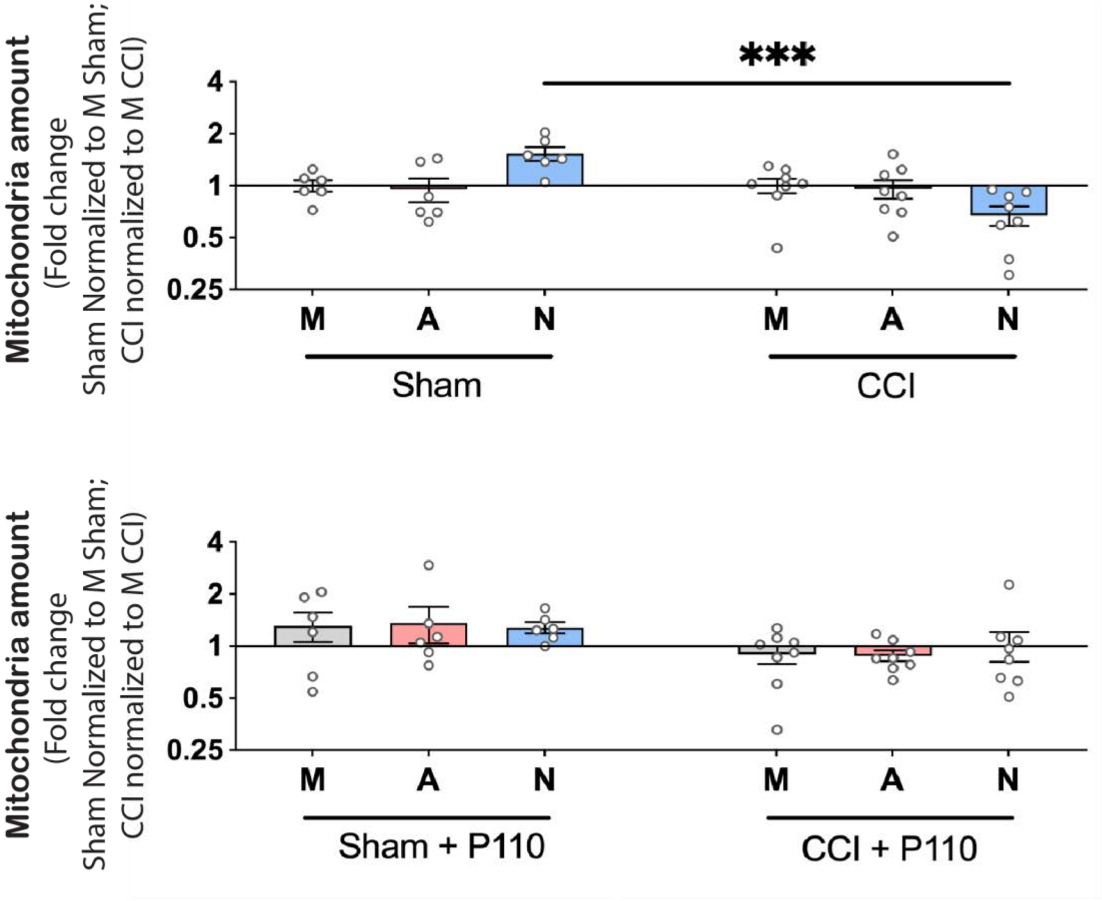


**Supplementary Fig. 18 Activated naïve neurons re-uptake dysfunctional mitochondria from the extracellular environment.** DNA quantification of mitochondria released from activated by treatment with conditioned media neurons for 24 hours. Data presented as mean ±SEM of n=6-8 scaffolds per condition from 2 independent experiments. *** indicates a significant difference (p< 0.001; one-way or two-way ANOVA (with Tukey's post-hoc test) between experimental groups).

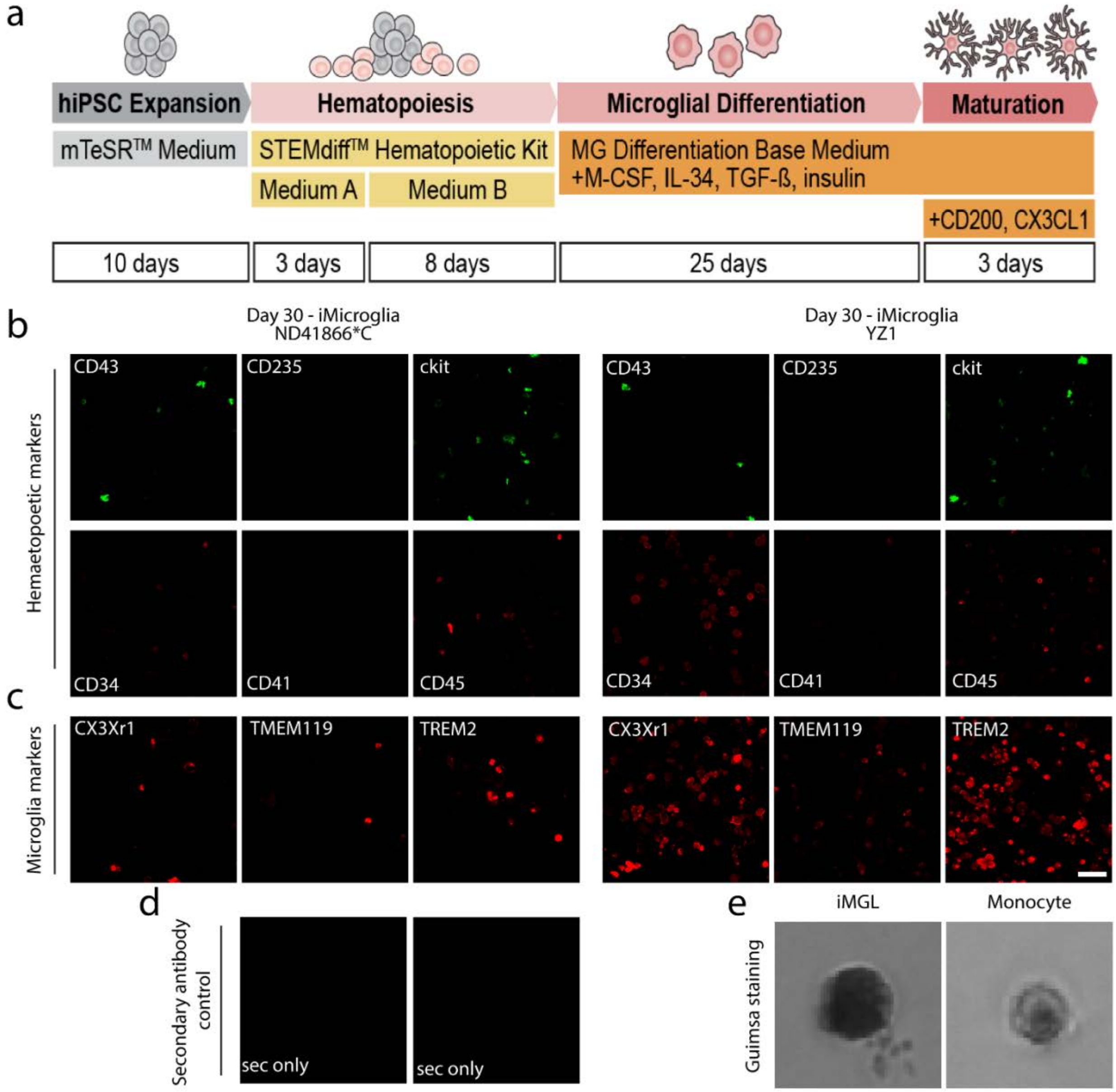


**Supplementary Fig. 19. iPSC-derived microglia characterization. a,** Schematic representation of the microglia differentiation protocol from induced pluripotent stem cells of two healthy donors (YZ1 and ND41866*C). **b,** Hematopoietic markers: CD43, CD34, CD235, CD41, and c-kit; **c,** Microglial markers: CD45, CX3CR1, TMEM119, TREM2; **d,** Secondary antibody control; **e,** Guimsa staining in comparison to human monocytes. Scale bar: 50µm.

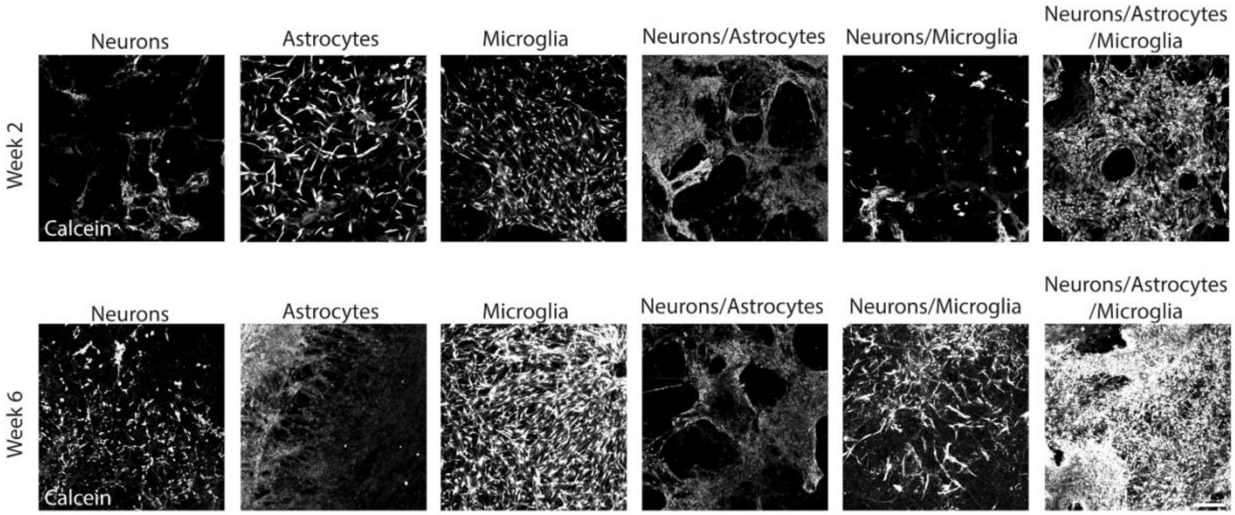


**Supplementary Fig. 20. Tricultures with iPSC derived microglia were alive at 6 weeks before controlled cortical impact injuries.** Calcein staining of live samples at 2 and 6 weeks after seeding. Scale bar: 100μm.

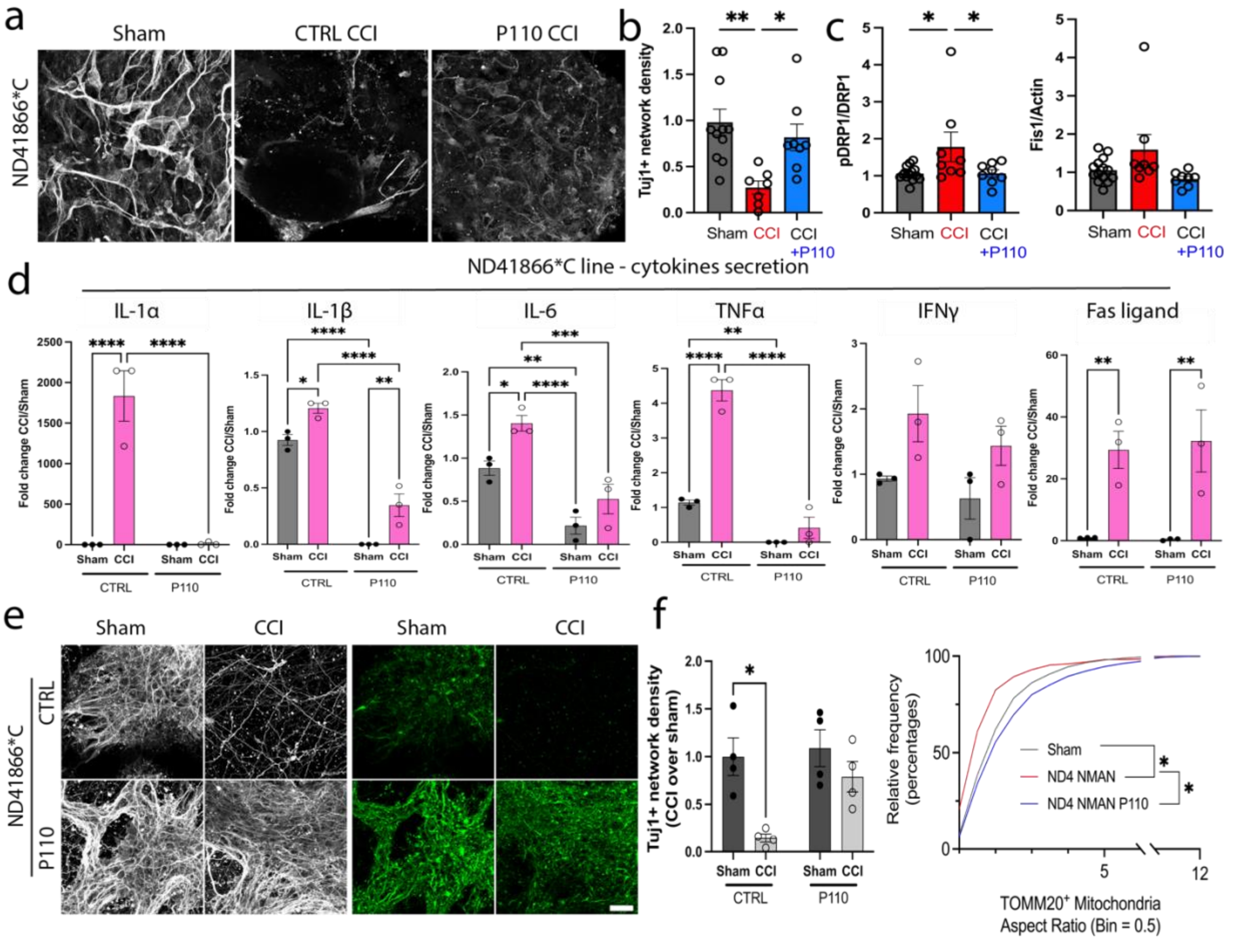

**Supplementary Fig. 21. Fragmented mitochondria released from iPSCs derived microglia (ND41866*C line from a healthy donor) induced neurodegeneration and neuroinflammation progression 24 hours after controlled cortical impact injury. a,** Representative images of Tuj1 network density in tricultures with iPSC derived microglia. **b,** Quantification of the Tuj1 neuronal network density in control and P110 treated groups of NAM group. **c,** Mitochondria fission associated pDRP1 and FIS1 proteins expression in control and P110 treated NAM groups. **d,** Inflammatory cytokines secretion post-CCI in YZ1 triculture groups. **e,** Representative images, and **f,** Quantifications of Tuj1 neuronal and TOMM20 mitochondria networks in naïve neurons treated with mitochondria isolated from astrocytes activated with mitochondria isolated from YZ1 microglia treated with conditioned media collected from 24h injured neurons. Data presented as mean ±SEM of n=2-10 scaffolds per condition. *, **, ***, **** indicates significant difference (p<0.05, 0.01, 0.001, 0.0001 respectively; two-way ANOVA (analysis of variance) or one-way ANOVA between control and experimental groups). Experiments were replicated at least three times. Scale bar: 50µm

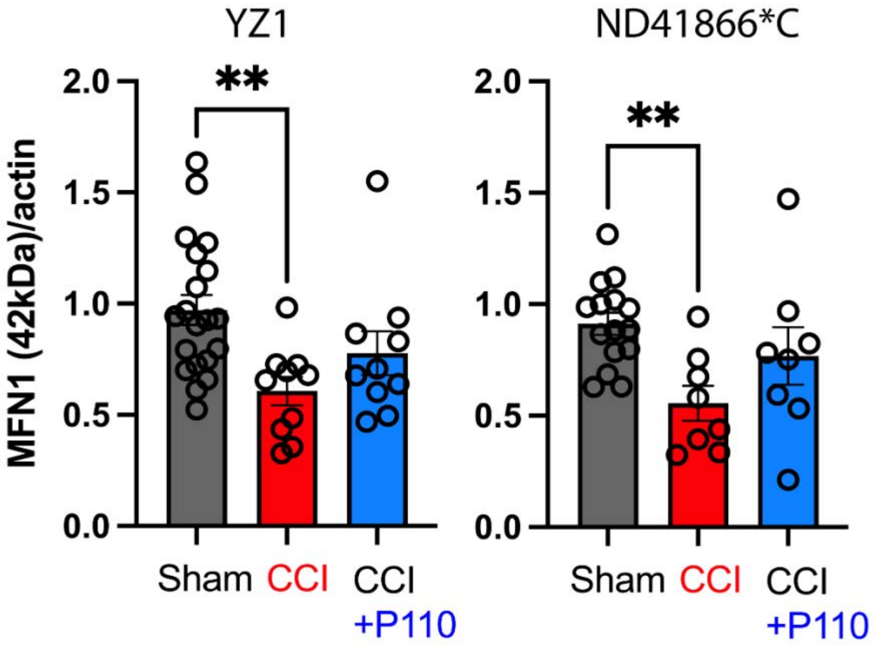


**Supplementary Fig. 22. Secondary neurodegeneration in tricultures with iPSC-derived microglia (both donors) was associated with decreased mitochondria fusion.** Mitofusin 1, 2 (42 kDa isoform) protein amount quantification of Western blots 24 hours after injury. Data presented as mean ±SEM of three independent experiments with n=2-4 scaffolds per condition. *, **, ***, **** indicates significant difference (p<0.05, 0.01, 0.001, 0.0001 respectively; two-way ANOVA (analysis of variance) between control and experimental groups).

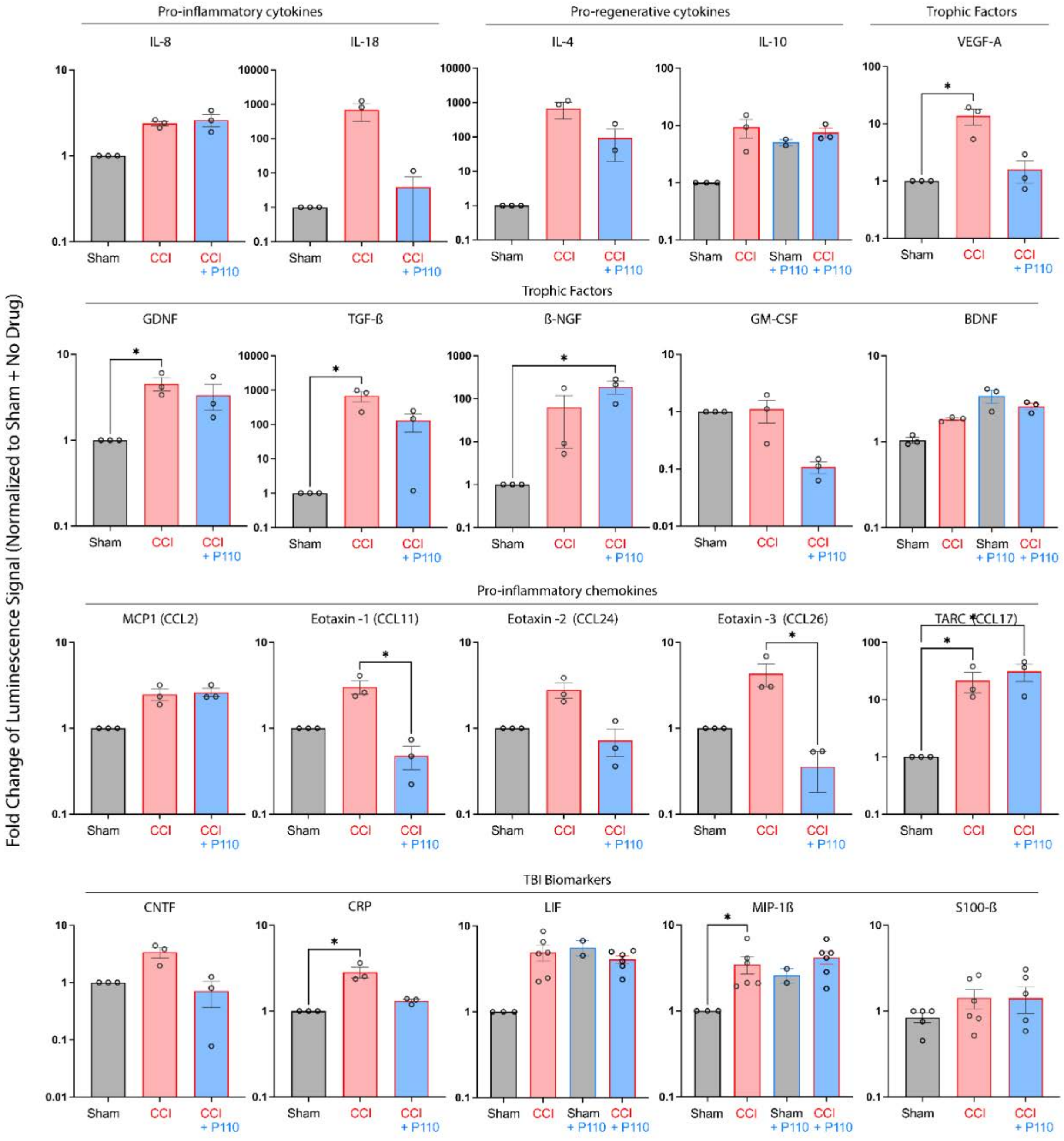


**Supplementary Fig. 23. Increased mitochondria fission was associated with neuroinflammation in tricultures with iPSC-derived microglia (YZ1 line) 24 hours after the injury.** The production level of: Pro-inflammatory and pro-regenerative cytokines, trophic factors, pro-Inflammatory chemokines, and TBI biomarkers. Data presented as mean ± SEM of four independent experiments with each data point is the average for n=2-4 scaffolds per condition. *, **, ***, **** indicate significant differences (p<0.05, 0.01, 0.001, 0.0001 respectively; two-way ANOVA (analysis of variance) between control and experimental groups).

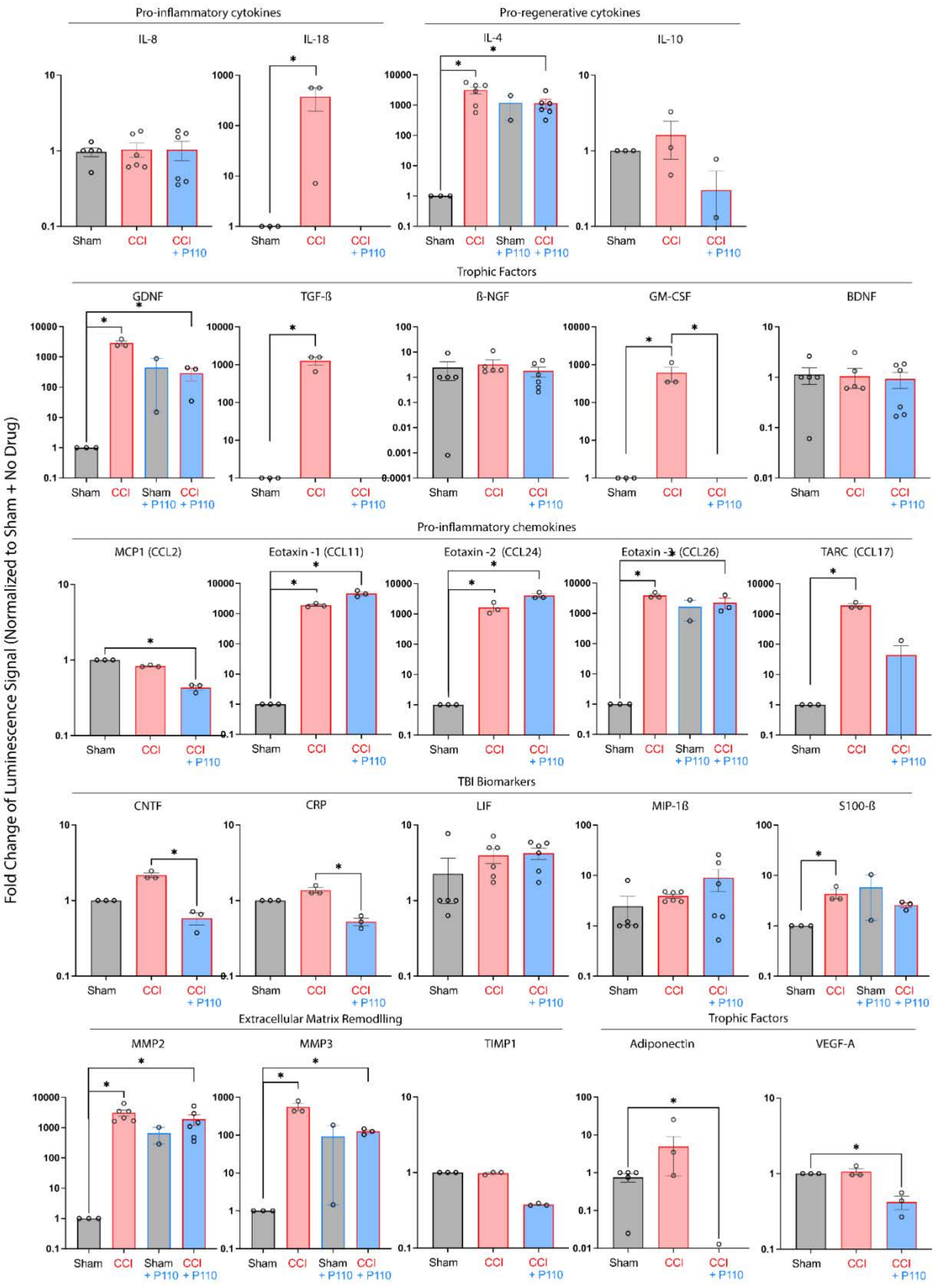

Pro-inflammatory cytokines
Pro-regenerative cytokines
IL-8
IL-18
IL-4
IL-10
Trophic Factors
GDNF
TGF-ß
ß-NGF
GM-CSF
BDNF
Pro-inflammatory chemokines
MCP1 (CCL2)
Eotaxin -1 (CCL11)
Eotaxin -2 (CCL24)
Eotaxin -3 (CCL26)
TARC (CCL17)
TBI Biomarkers
CNTF
CRP
LIF
MIP-1ß
S100-ß
Extracellular Matrix Remodlling
Trophic Factors
MMP2
MMP3
TIMP1
Adiponectin
VEGF-A
Fold Change of Luminescence Signal (Normalized to Sham + No Drug)
Sham
CCI
CCI + P110
Sham + P110

**Supplementary Fig. 24. Increased mitochondria fission was associated with neuroinflammation in tricultures with iPSC-derived microglia (ND41866*C line) 24 hours after the injury.** The production level of: Pro-inflammatory and pro-regenerative cytokines, trophic factors, pro-Inflammatory chemokines, and TBI biomarkers. Data presented as mean ± SEM of four independent experiments with each data point is the average for n=2-4 scaffolds per condition. *, **, ***, **** indicate significant differences (p<0.05, 0.01, 0.001, 0.0001 respectively; two-way ANOVA (analysis of variance) between control and experimental groups).

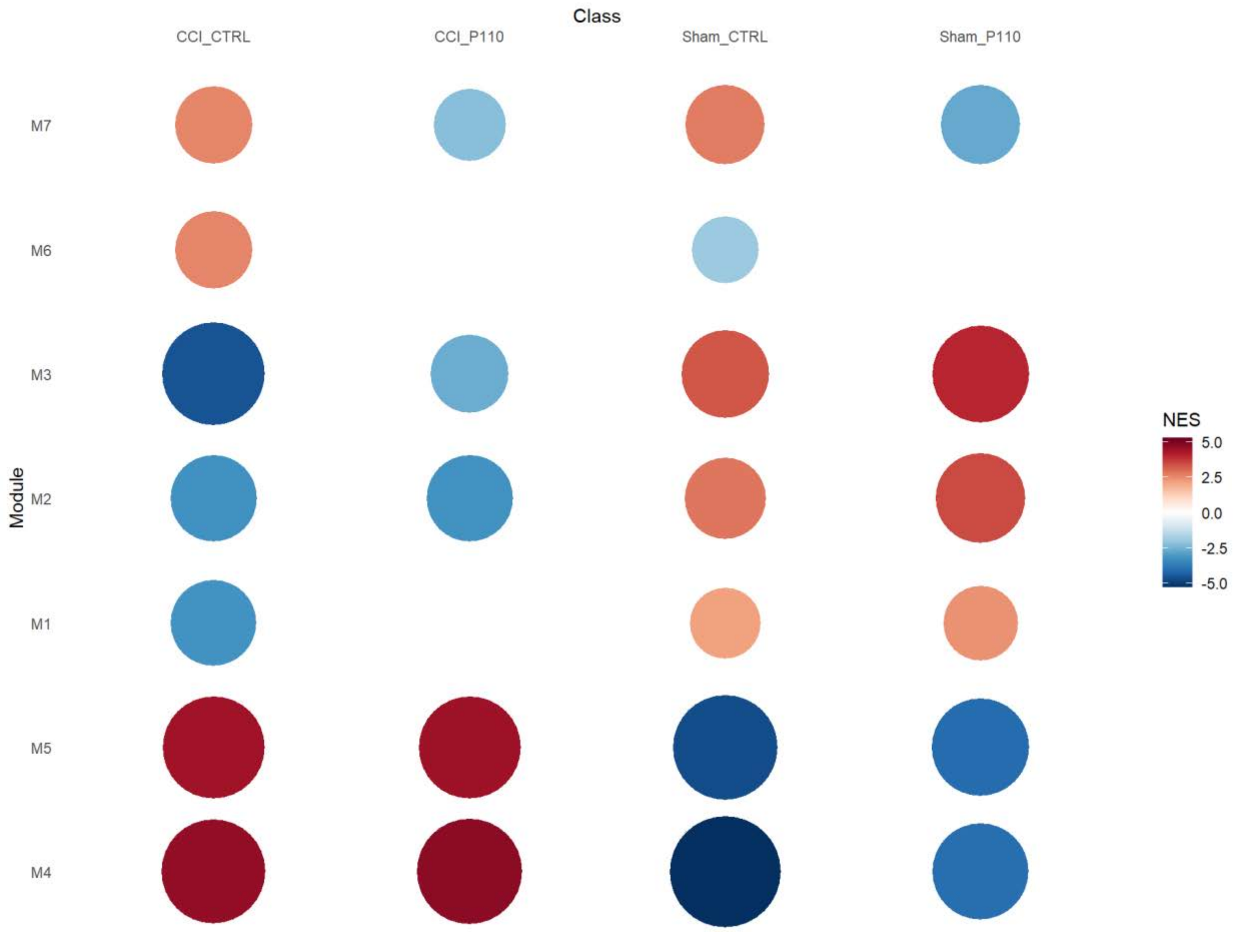


**Supplementary Fig. 25. Co-expression network analysis revealed a significant shift in gene expression associated with cellular proliferation (Module 5), extracellular matrix remodeling (Modules 2, 4), interferon signaling (Module 6), and neuronal systems (Modules 1, 3) in human 3D triculture model in response to contusion injury.** P110 treatment removed the positive association of contusion injury with interferon-associated inflammation in Module 6, while In Module 7, P110 treatment was negatively associated with collagen formation and assembly but not injury. In modules 1-3, gene expression was negatively associated with a contusion, while in Modules 4-5, the association was positive. Changes in differentially expressed genes 24 hours post-contusion injury in human 3D triculture model composed of induced neural stem cells derived neurons, primary astrocytes, and iPSC-derived microglia from two healthy donors YZ1 and ND418664*F (n=6).

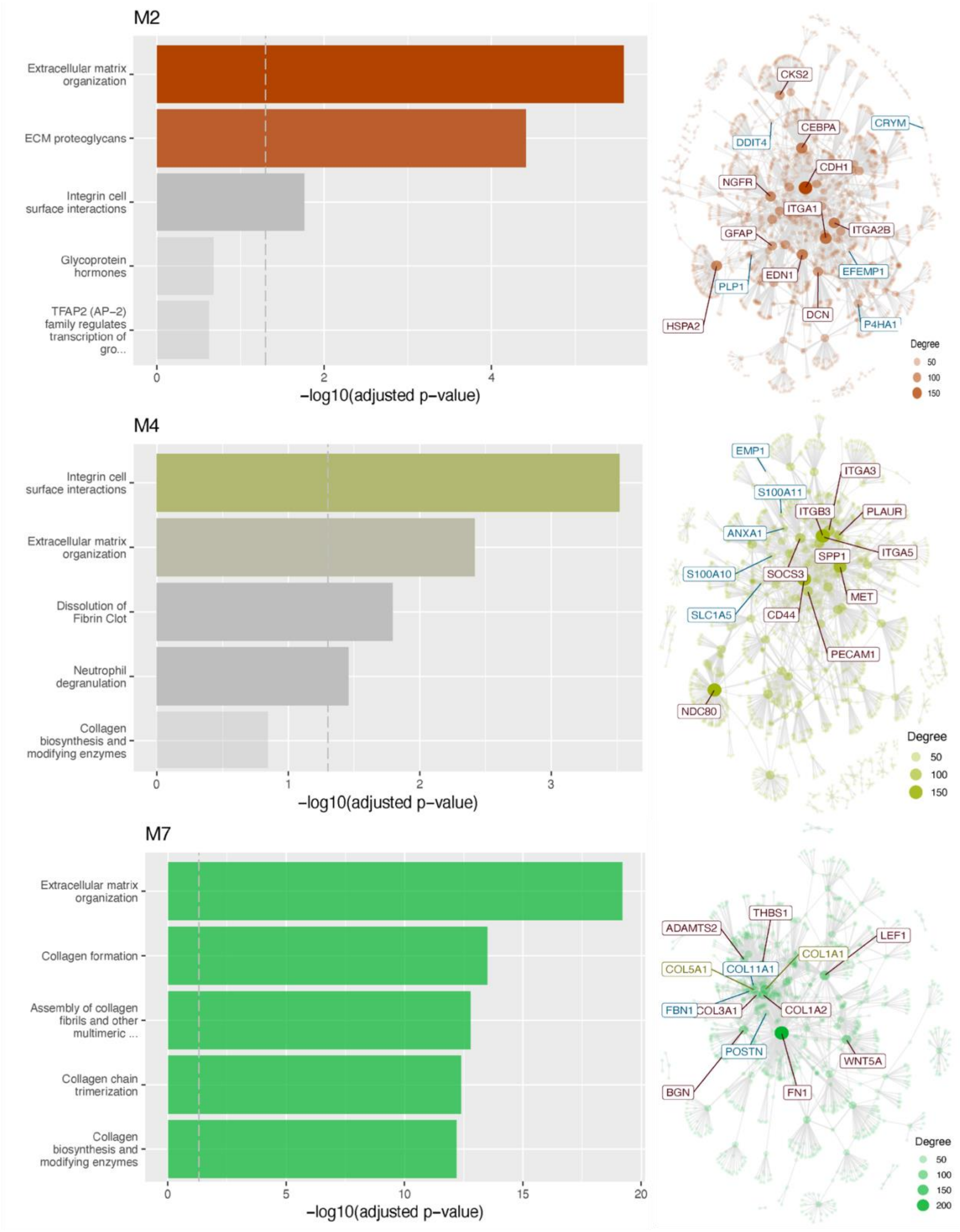


**Supplementary Fig. 26. Contusion injury-induced shift in the gene expression of extracellular matrix components in Modules 2 and 4.** Gene set enrichment analysis revealed distinct expression modules associated with M2 – ECM proteoglycans (negatively associated with a contusion in CTRL and P110); M4

– integrin cell surface interactions (positively associated with a contusion in CTRL and P110); M7 – collagen formation and assembly (positively associated with P110 treatment, but not the injury) and their corresponding regulators. Changes in differentially expressed genes 24 hours post-contusion injury in human 3D triculture model composed of induced neural stem cells derived neurons, primary astrocytes, and iPSC-derived microglia from two healthy donors YZ1 and ND418664*F (n=6).

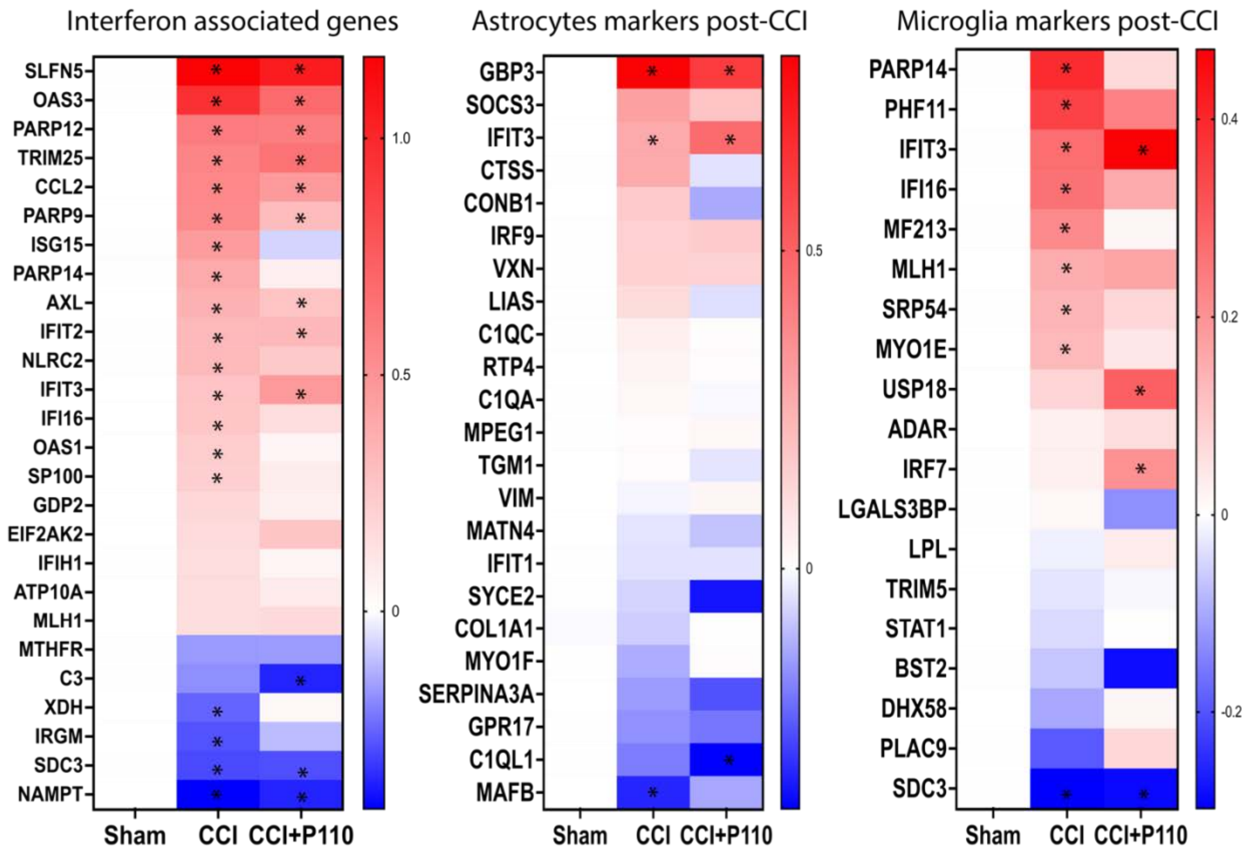


**Supplementary Fig. 27**. **Contusion injury induces activation of microglia-driven interferon-associated inflammation.** Differentially expressed genes in sham (sham vs sham P110), CCI (CCI vs sham), and CCI+P110 (CCI+P110 vs Sham+110) groups were detected (abs(Log2(Fold change))> 0.585 and q < 0.01). Changes in differentially expressed genes 24 hours post-contusion injury in human 3D triculture model composed of induced neural stem cells derived neurons, primary astrocytes, and iPSC-derived microglia from two healthy donors YZ1 and ND418664*F (n=6).

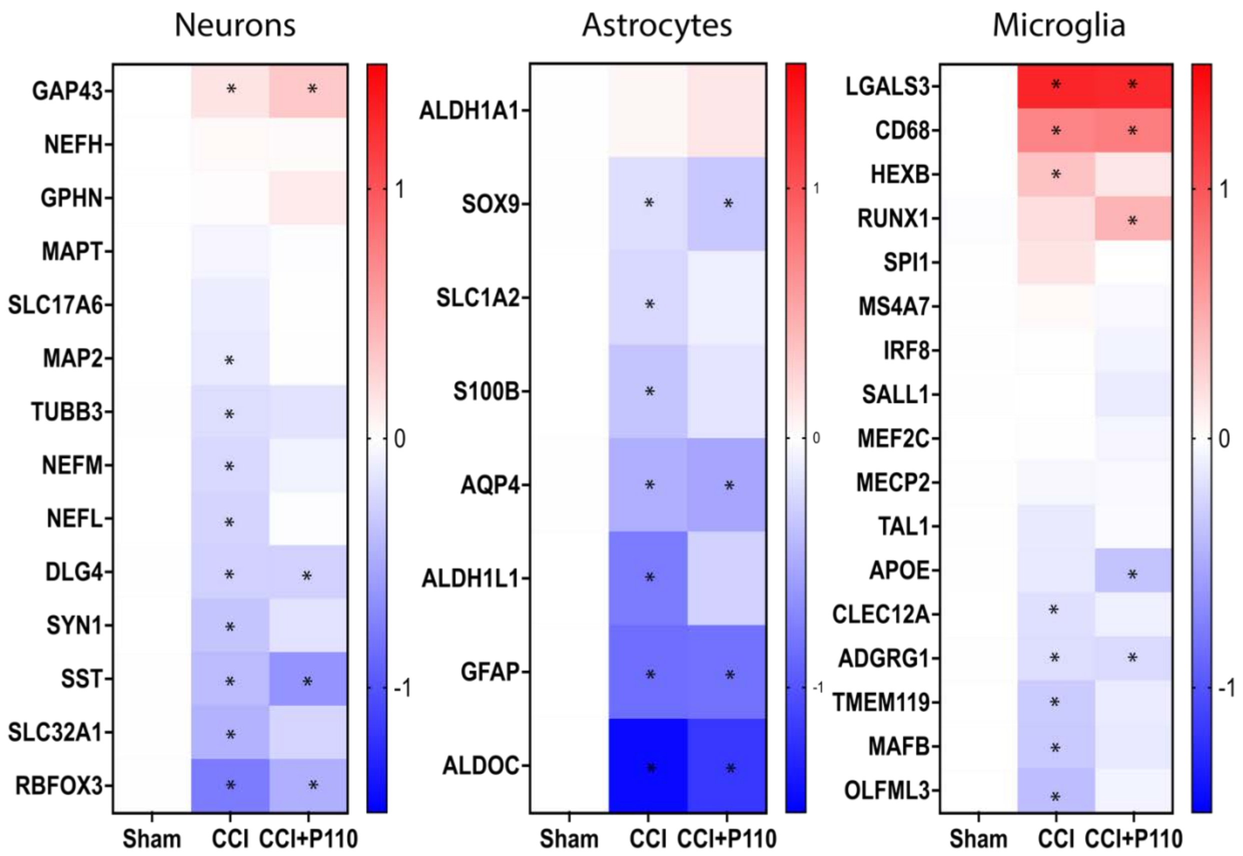


**Supplementary Fig. 28**. **Contusion injury induces a negative shift in neuronal and astrocytic pan-markers but a positive shift in microglial.** Differentially expressed genes in sham (sham vs sham P110), CCI (CCI vs sham), and CCI+P110 (CCI+P110 vs Sham+110) groups were detected (abs(Log2(Fold change))> 0.585 and q < 0.01). Changes in differentially expressed genes 24 hours post-contusion injury in human 3D triculture model composed of induced neural stem cells derived neurons, primary astrocytes, and iPSC-derived microglia from two healthy donors YZ1 and ND418664*F (n=6).

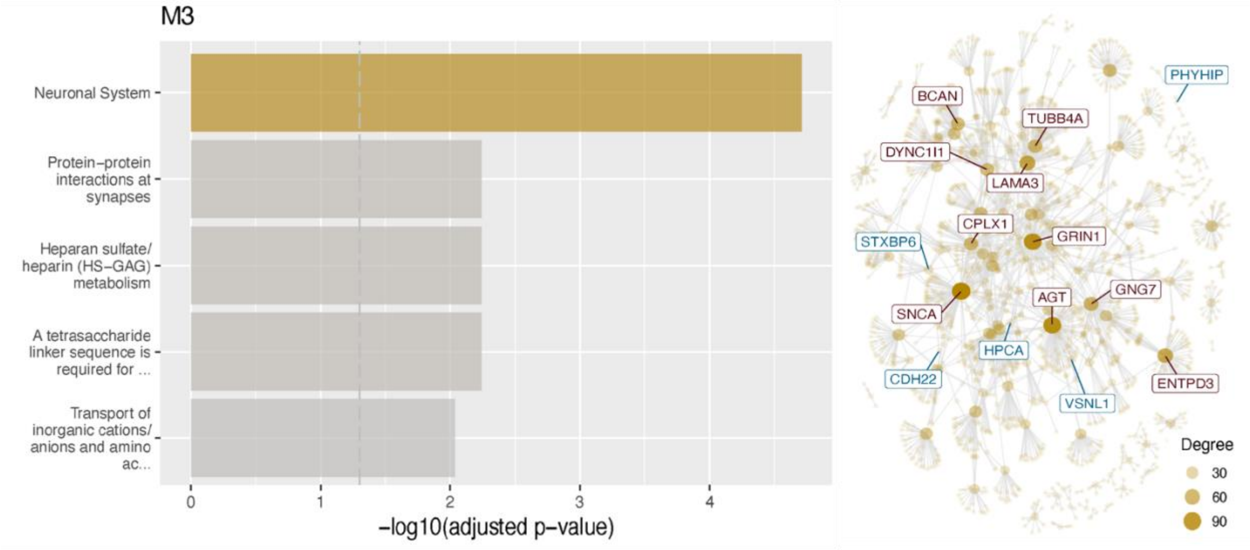


**Supplementary Fig. 29. Contusion injury-induced shift in the gene expression of extracellular matrix components.** Gene set enrichment analysis revealed distinct expression modules associated with M3 – neuronal systems (negatively associated with a contusion in both CTRL and P110 groups) and their gene -regulators. Changes in differentially expressed genes 24 hours post-contusion injury in human 3D triculture model composed of induced neural stem cells derived neurons, primary astrocytes, and iPSC-derived microglia from two healthy donors YZ1 and ND418664*F (n=6).

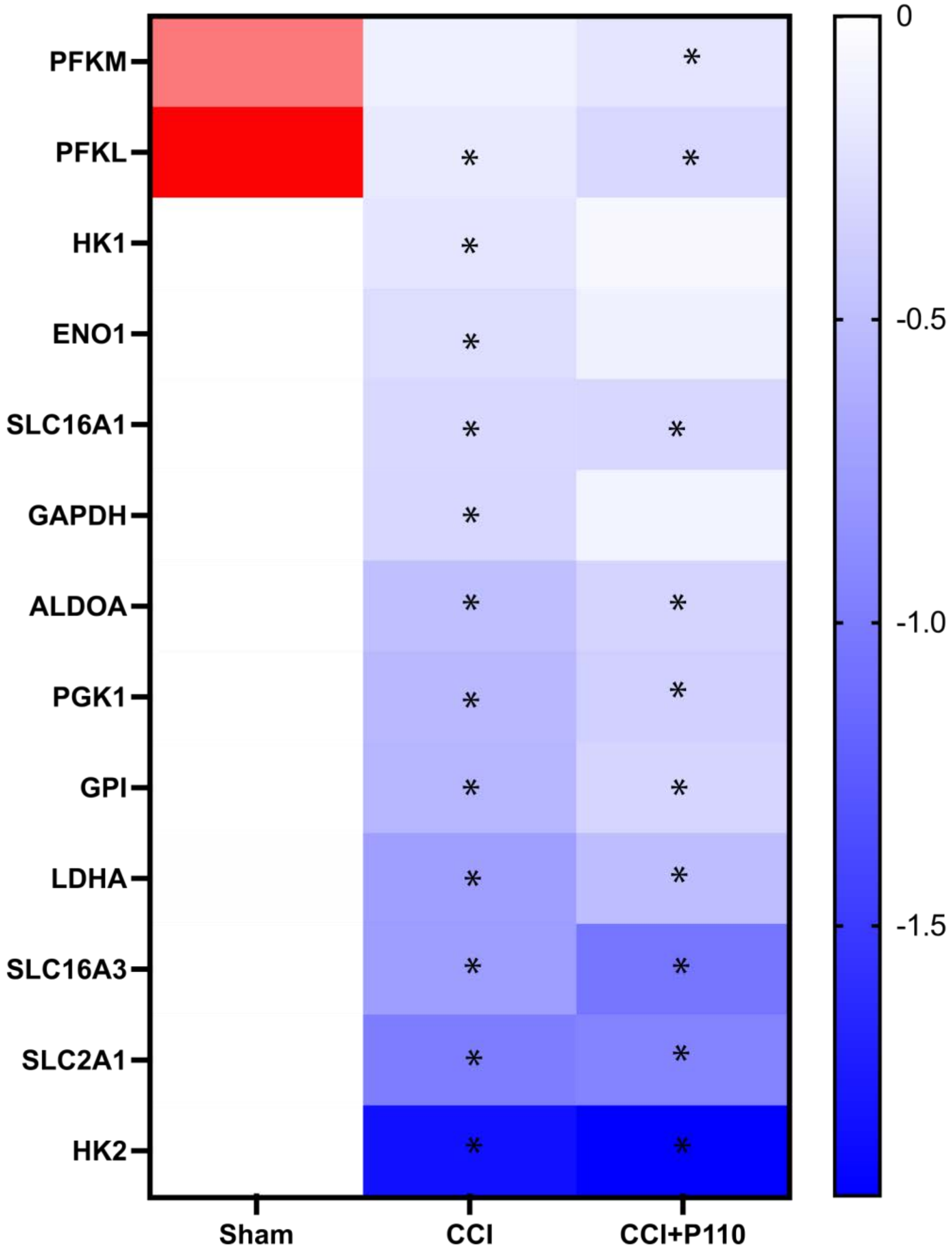


**Supplementary Fig. 30. Contusion injury induces glycolysis downregulation in the short term after the injury.** Differentially expressed genes in sham (sham vs sham P110), CCI (CCI vs sham), and CCI+P110 (CCI+P110 vs Sham+110) groups were detected (abs(Log2(Fold change))> 0.585 and q < 0.01). Changes in differentially expressed genes 24 hours post-contusion injury in human 3D triculture model composed of induced neural stem cells derived neurons, primary astrocytes, and iPSC-derived microglia from two healthy donors YZ1 and ND418664*F (n=6).

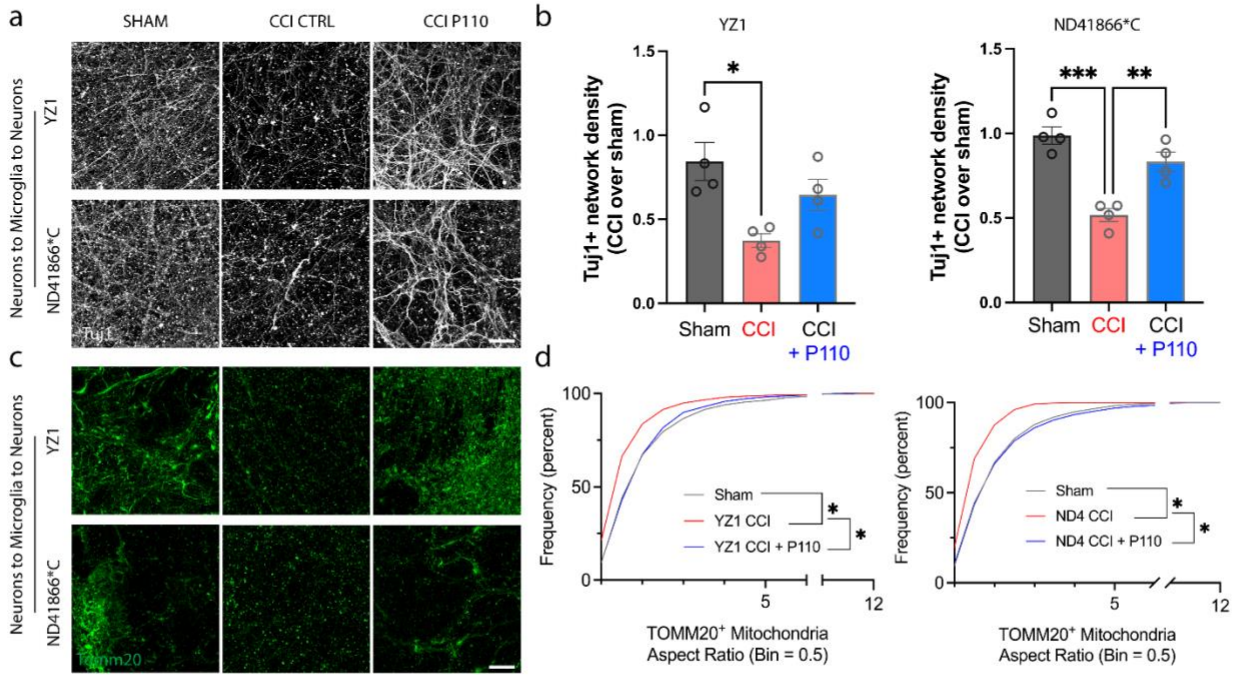


**Supplementary Fig. 31. Fragmented mitochondria released from iPSCs derived microglia (ND41866*C line from a healthy donor) induced neurodegeneration and neuroinflammation progression 24 hours after controlled cortical impact injury.** Representative images of **a,** Tuj1 and **c,** TOMM20 staining of naïve neurons treated for 24 hours with mitochondria isolated from microglia treated with conditioned media collected from 24h injured neurons with quantification of **b,** Tuj1 positive neuronal network density and **d,** TOMM20 positive mitochondria aspect ratios. Data presented in **(b, d)** mean ±SEM of n=2-10 scaffolds per condition. *, **, ***, **** indicates significant difference (p<0.05, 0.01, 0.001, 0.0001 respectively; one-way or two-way ANOVA (analysis of variance) between experimental groups). Experiments were replicated at least three times. Scale bar: 50 µm.

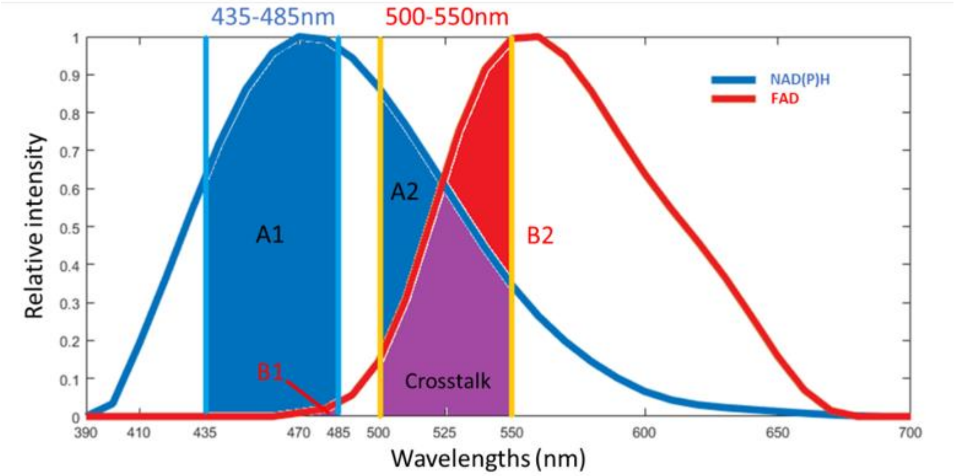

**Supplementary Fig. 32.** Spectra shapes of NAD(P)H and FAD. A1, A2. B1. B2 represents the area under the curve (AUC) of NAD(P)H and FAD at two different emission bands (435-485nm and 500-550nm), respectively. Purple color highlighted the crosstalk of NAD(P)H and FAD in both channels.

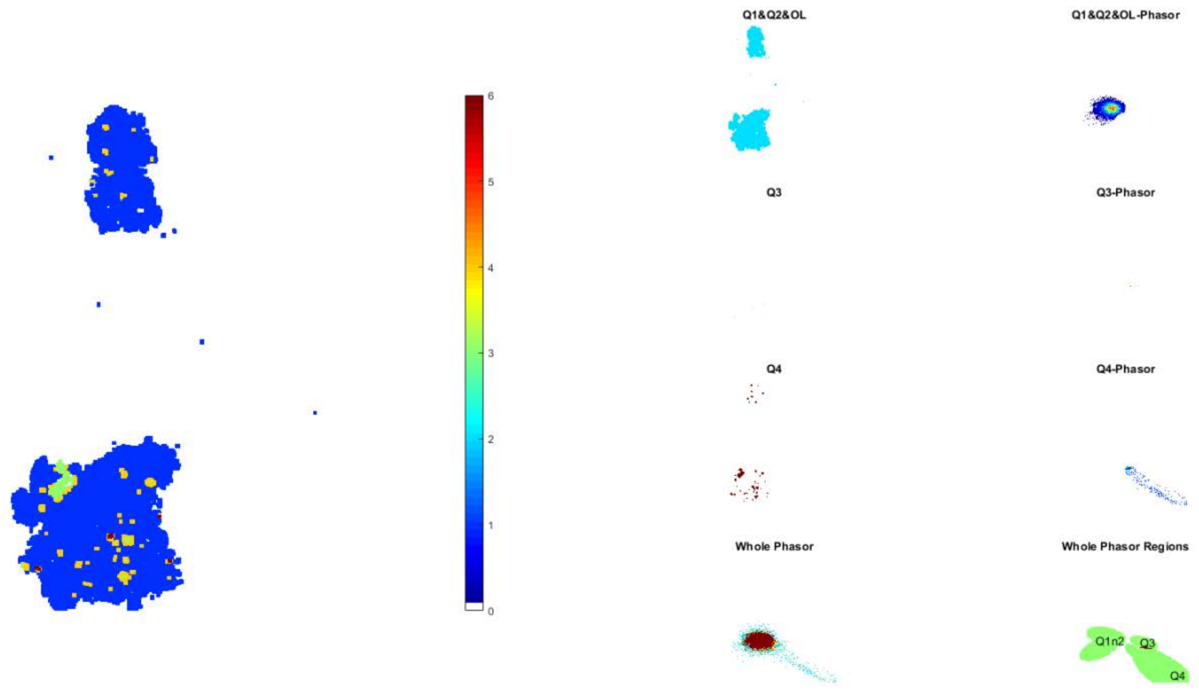


**Supplementary Fig. 33.** Representative Phasor-guided redox ratio area used for analysis.

**Redox distributions at Q3 and Q4 Phasor locations very different from Other three locations; The highly overlapped features among A-C indicate they may be NADH and NADPH**

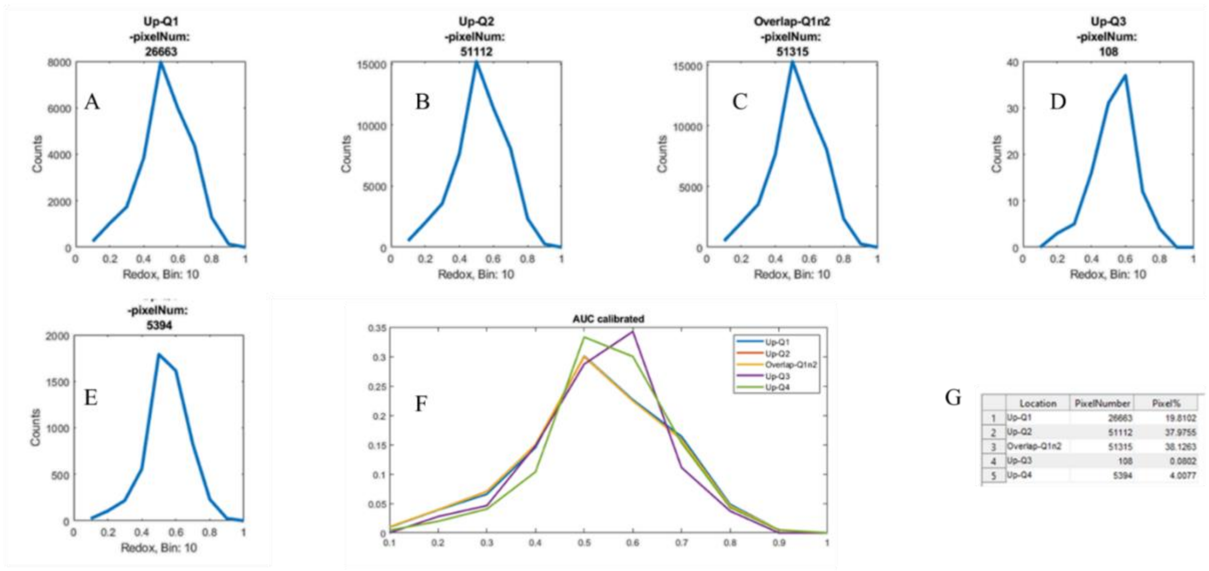


| | Location | PixelNumber | Pixel% |
|---|---|---|---|
| 1 | Up-Q1 | 26663 | 19.8102 |
| 2 | Up-Q2 | 51112 | 37.9755 |
| 3 | Overlap-Q1n2 | 51315 | 38.1263 |
| 4 | Up-Q3 | 108 | 0.0802 |
| 5 | Up-Q4 | 5394 | 4.0077 |

Example from iPSCs, neuron transfer, P110m Sham sample 3, all the pixels from different ROIs

**Supplementary Fig. 34.** Redox distribution for each phasor location (A, B, D, E) and overlapped region between Q1 and Q2(C). Super-imposed AUC calibrated redox distribution from every different phasor location (F). (G) the pixel number percentage of different phasor locations.
